# The maternal microbiome regulates infant respiratory disease susceptibility via intestinal Flt3L expression and plasmacytoid dendritic cell hematopoiesis

**DOI:** 10.1101/2023.01.05.522516

**Authors:** Md. Al Amin Sikder, Ridwan B. Rashid, Tufael Ahmed, Ismail Sebina, Daniel Howard, Md. Ashik Ullah, Muhammed Mahfuzur Rahman, Jason P. Lynch, Bodie Curren, Rhiannon B. Werder, Jennifer Simpson, Alec Bissell, Mark Morrison, Carina Walpole, Kristen J. Radford, Vinod Kumar, Trent M. Woodruff, Tan HuiYing, Ayesha Ali, Gerard E. Kaiko, John W. Upham, Robert D. Hoelzle, Páraic Ó Cuív, Patrick G. Holt, Paul G. Dennis, Simon Phipps

## Abstract

Severe lower respiratory infection (sLRI) are a major cause of infant morbidity and mortality, and predispose to later chronic respiratory diseases such as asthma. Poor maternal diet during pregnancy is a risk factor for sLRI in the offspring. Here we demonstrate in mice that a maternal low-fibre diet (LFD) disrupts plasmacytoid and conventional dendritic cell (DC) hematopoiesis in the offspring, predisposing to sLRI and subsequent asthma. The LFD alters the composition of the maternal milk microbiome and assembling infant gut microbiome, ablating the induction of a developmental wave of the non-redundant DC growth factor Flt3L by neonatal intestinal epithelial cells. Therapy with a propionate-producing bacteria isolated from the milk of high-fibre diet-fed mothers, or supplementation with propionate, confers protection against sLRI by restoring gut Flt3L expression and pDC hematopoiesis. Our findings identify a microbiome-dependent Flt3L axis in the gut that regulates pDC hematopoiesis in early life and confers disease resistance.

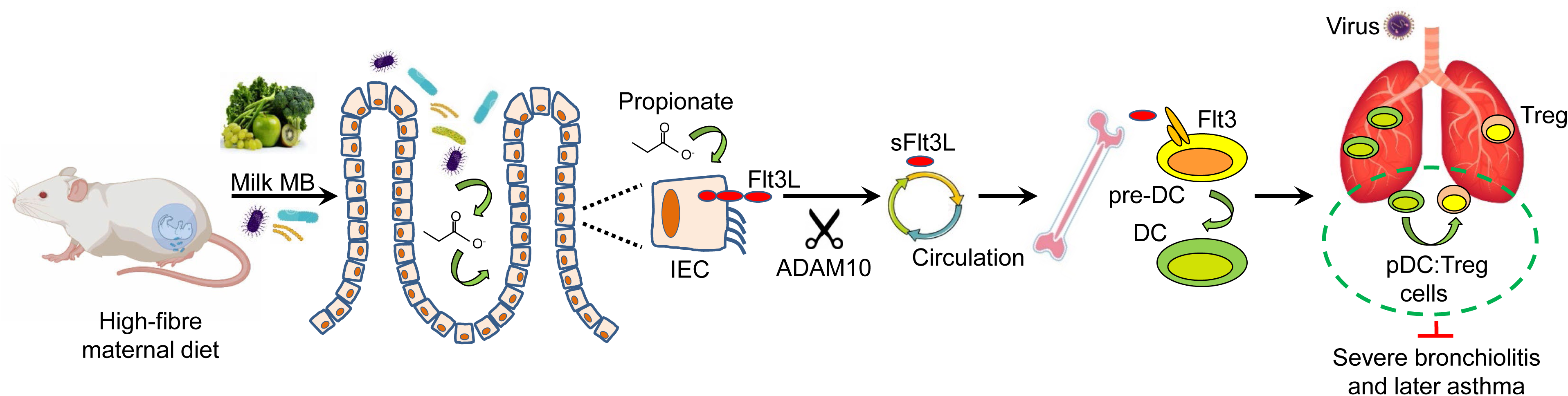

## Introduction

Severe lower respiratory tract infections (sLRI) such as bronchiolitis and pneumonia are the leading cause of childhood mortality globally, and a major cause of infant hospitalisations in high income societies ^1, 2^. Early and recurrent LRIs also impact lung growth and function ^3, 4^, and increase the risk of developing chronic lung diseases, such as asthma, bronchiectasis and chronic obstructive pulmonary disease ^5–10^. sLRIs are highly prevalent in infancy at a time when diverse microbial communities are being assembled in the gut. These microbes and their products are essential for the development of a healthy immune system ^11^. Accordingly, perturbations to the composition and maturation of the microbiota in early life have been linked to LRI severity and the development of chronic inflammatory diseases that commence in childhood ^12–16^.

While the underlying mechanisms remain incompletely understood, it is widely appreciated that fermentable dietary fibre influences gut microbiome composition, and that the resultant immunomodulatory metabolites (e.g. short chain fatty acids, SCFAs), enhance homeostasis of immune cells, most notably dendritic cells (DC) and regulatory T (Treg) cells ^17–21^. For example, DC numbers are lower in germ-free mice and can be increased by microbiota transplantation or SCFA supplementation, implicating an important role for the microbiome ^17, 19^. DCs are integral to the healthy functioning of the immune system; they contribute to innate immunity, shape the adaptive response, and promote Treg cell differentiation. In a neonatal mouse model of human early-onset asthma, we recently demonstrated that the temporal depletion of plasmacytoid DC (pDC), a specialized subset of DCs that initiate antiviral immunity, perturbs Treg cell expansion, predisposing to a severe viral LRI in infancy and the subsequent development of experimental asthma in later life ^22, 23^. While these findings elucidated an important pDC-Treg interaction that confers protection, and provided an explanation as to why infants with low numbers of pDC are at risk of developing greater LRI symptoms, wheeze, and childhood asthma ^24, 25^, the processes that promote healthy DC homeostasis or give rise to the ‘pDC-low’ phenotype in infancy remain ill- defined.

Risk factors associated with infant sLRI include never breastfeeding, malnutrition, and age below two months ^26, 27^. However, in addition to risk factors that operate postnatally, it has become evident that a range of maternal environmental exposures, particularly those that modify the microbiome, can also affect sLRI and asthma risk in the offspring ^26, 28–31^. Of major interest, a large population study found that a carbohydrate-rich, fruit/vegetable-poor maternal diet predisposes the infant to sLRI with respiratory syncytial virus (RSV)^32^. In this study, we demonstrate that a maternal low-fibre diet (LFD) during pregnancy and the subsequent pre-weaning period predisposes the offspring to sLRI during the high-risk early postnatal period by disturbing the microbiome’s influence on neonatal DC development and maturation. Critically, we discovered that these effects are associated with microbial metabolite-mediated fluctuations in the production of the non-redundant DC growth factor Flt3L (Fms-like tyrosine kinase 3 ligand), which is produced at high levels by intestinal epithelial cells in the neonatal gut. Our findings identify a critical pathway by which the maternal microbiota affects neonatal pDC and Treg cell homeostasis and susceptibility to lung disease in early life.

## Results

### A maternal low-fibre diet (LFD) predisposes the offspring to sLRI and later asthma in mice

To assess the impact of poor maternal diet on the severity of infant bronchiolitis, the offspring of breeding-age mice fed a low-fibre diet (LFD) or high-fibre diet (HFD) were inoculated at postnatal day (PND) 7 with PVM (pneumonia virus of mice) – the murine equivalent of RSV, which is also in the genus Pneumovirus (Fig.S1A). Diet did not alter the weight of the pregnant mothers or pups at birth (Fig.1A and Fig.S1B,C). However, post-PVM infection, pups reared by LFD-fed mothers (hereafter referred to ‘LFD-reared pups’) gained less weight (Fig. 1B), had increased viral load in the airway epithelium, and produced lower amounts of the antiviral cytokine IFN-λ (Fig.1C,D).

**Figure 1.**
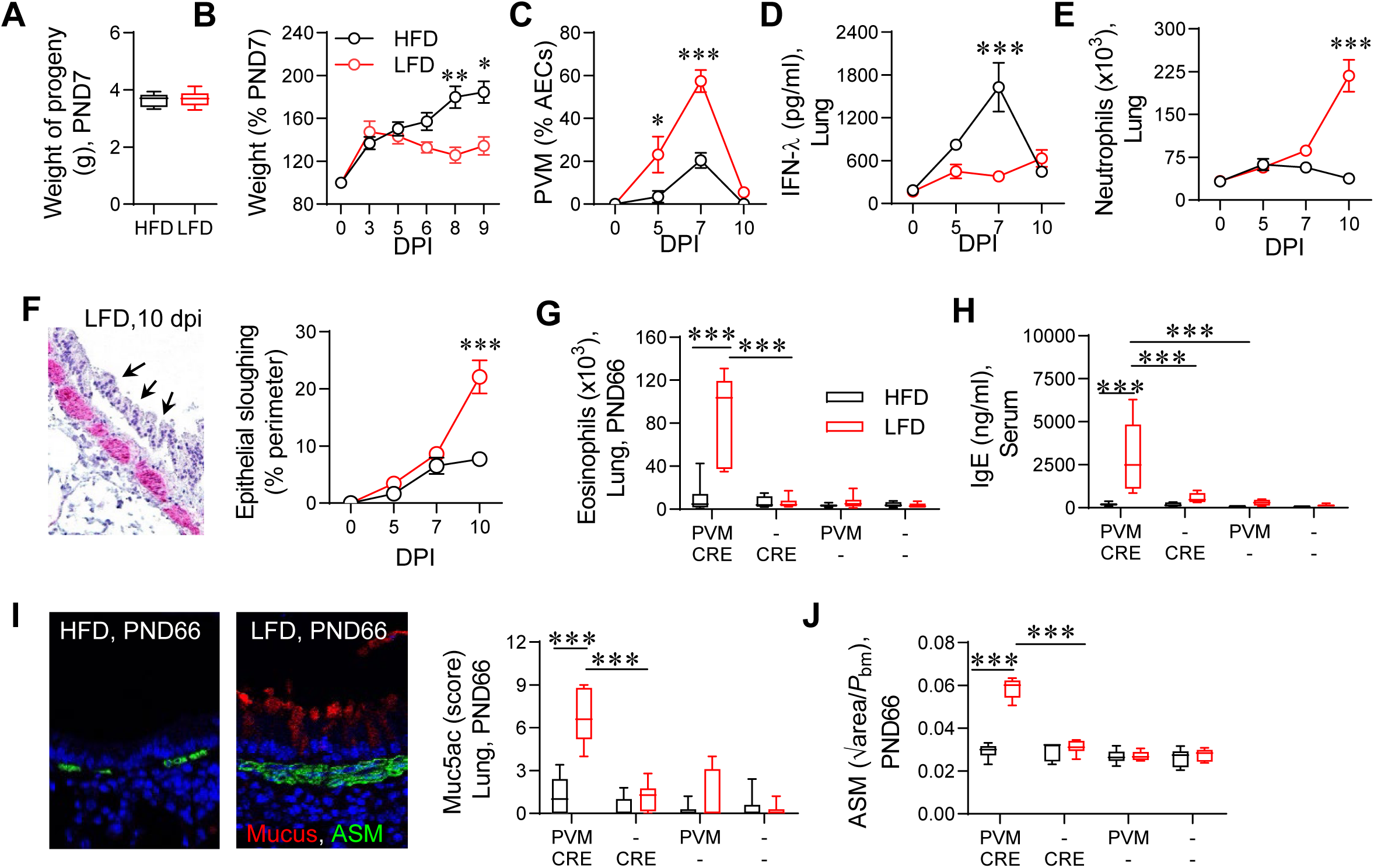
A maternal low-fibre diet (LFD) predisposes the offspring to sLRI and later asthma in mice. (A-F) Data from neonatal LRI phase. (A) Weight of the pups at PND7. (B) Pup weight following virus infection. (C) PVM immunoreactivity as a percentage of AECs. (D) IFN-λ protein in lung homogenate. (E) Lung neutrophils. (F) Representative image of epithelial sloughing in infected LFD-reared pup and quantification of epithelial sloughing. (G-J) Data from later-life asthma phase (for experimental study design see Fig.S1A. (G) Lung eosinophils. (H) Total IgE concentration in serum. (I) Representative image of Muc5ac (red) and ASM (green) immunofluorescence in lungs of HFD- and LFD-reared mice (left panel), quantification of muc5ac (right panel) at PND66. (J) ASM area at PND66. Black boxes or circles represents HFD-reared pups, red boxes or circles represent LFD-reared pups. Data are represented as mean ± SEM (B-F) or box-and-whisker plots (median, quartiles, and range) (A, G-J), n=4-15 mice per group. *P* values (* p<0.05, ** p<0.01, *** p<0.001) were derived by Mann Whitney U test (A) comparing HFD-reared with LFD-reared mice or two-way ANOVA (C-J) followed by Tukey’s multiple comparisons test.

Notably, the LFD-reared pups developed the hallmark features of a sLRI, including elevated pro-inflammatory cytokine expression, neutrophilic inflammation, epithelial sloughing, and mucus hyper-production (Fig.1E,F and Fig.S1D,E). This excessive inflammatory response and tissue damage in LFD-reared pups was associated with the onset of type-2 inflammation and airway remodeling (Fig.S1F-J). Interestingly, the LFD-reared pups developed sLRIs upon inoculation with rhinovirus (RV), another causative agent of bronchiolitis (Fig.S1K-Q). In light of observations from human birth cohort studies demonstrating that aeroallergen sensitization enhances the asthmatogenic effects of early viral bronchiolitis, we exposed the pups to PVM in early life and then low-dose cockroach extract (CRE) in later life (Fig.S1A)^23^. Simulating the human epidemiology, mice with sLRI in infancy (i.e. LFD-reared pups) progressed to an asthma-like phenotype, characterised by increases in lung eosinophils, airway smooth muscle area, mucus, serum IgE, and type-2 cytokine production (Fig.1G-J and Fig.S2A-D). In contrast, exposure to low-dose CRE in the absence of sLRI was not sufficient to induce experimental asthma, congruent with findings that show the majority of children, including those sensitized to ubiquitous perennial aeroallergens present virtually continuously in the indoor environment, remain clinically unresponsive ^33, 34^. Similar findings were observed when mice were inoculated with RV (rather than PVM) in early-life and then CRE in later-life (Fig.S2E-I).

### Maternal LFD decreases neonatal pDC and Treg cell expansion

Treg cells calibrate inflammatory processes. We previously identified that Neuropilin-1^+^ (Nrp-1) Treg cells, putative thymic (t)Treg cells ^35^, are expanded by pDCs and prevent immune-mediated tissue damage during PVM infection ^22^. Here, in response to infection, Treg cell numbers were significantly higher in the mediastinal lymph nodes (medLN) and lungs of HFD- compared to LFD- reared pups, consistent with disease severity (Fig.2A,B; gating strategy in Fig.S3A). As Treg cells were significantly lower in the lungs and medLNs of LFD-reared pups prior to infection (Fig.2A,B), we next assessed other lymphoid tissues and the gut, an important site of Treg cell differentiation and accumulation ^18, 20, 21^. In LFD-reared pups, Treg cells numbers were lower in the bone marrow (BM), mesenteric lymph nodes (mesLN), small intestine (SI), and colon (Fig.2C) - a phenotype that was unrelated to proliferation (Fig. S3B). Of note, Treg cells in the SI and colon were negative for Nrp-1 (∼50% Helios^+^; Fig.S3C), suggestive of peripheral (p)Treg cells ^36^, whereas those in the medLN and lung were >90% and ∼60% Nrp-1^+^ respectively and predominantly Helios^+^ (Fig.S3C), suggestive of thymic (t)Treg cells ^35^. Lower CD39 expression on lung Treg cells, and attenuated IL- 10 expression in the BALF indicated a lack of immunoregulation in the LFD-reared pups (Fig.S3D), a phenocopy of pDC-depleted neonatal mice infected with PVM ^22^, suggesting an upstream defect in the pDC compartment. As postulated, pDC numbers (gating strategy in Fig.S3E) in the lungs and medLN were significantly lower at 5 and 7 dpi in LFD-reared pups (Fig.2D,E), an effect that was unrelated to proliferation (Fig.S3F). As with the neonatal Treg cells, pDC numbers were lower in LFD-reared pups in the medLN, lungs, spleen, SI, colon, and mesLN at steady-state (Fig.2F). To investigate whether the diet affected the expression of co-stimulatory molecules on pDCs we performed spectral flow cytometry. Prior to infection, only minor shifts in expression were apparent in the bone marrow and lung pDCs. At 5 dpi, semaphorin4a (the cognate ligand of Nrp-1 and an effector of Treg cell stability ^22, 37^) and PD-L1 expression was significantly higher on pDCs in the medLN of HFD-reared pups, whereas CD86 and OX40L were more highly expressed on lung pDCs of LFD-reared pups (Fig.2G), suggesting that the pDC compartment is altered functionally as well as numerically. Conventional (c)DC1 and cDC2 numbers (gating strategy in Fig.S4)^38^ were also numerically lower in the lungs and medLN of LFD-reared pups at 5 dpi, although both subsets increased by 10 dpi (Fig.S5A,B). This observation was confirmed using an alternative immunophenotyping strategy (Fig.S5C)^39, 40^. The lower numbers of all three DC subsets was indicative of impaired DC hematopoiesis in the BM, and consistent with this, the numbers of cDC1 and cDC2s, as well as the major DC progenitor populations (*viz*. macrophage-DC progenitors, common DC progenitors, common lymphoid progenitors and pre-DCs; gating strategy in Fig.S5E) were all lower in the BM of LFD-reared pups (Fig.S5D,F).

**Figure 2.**
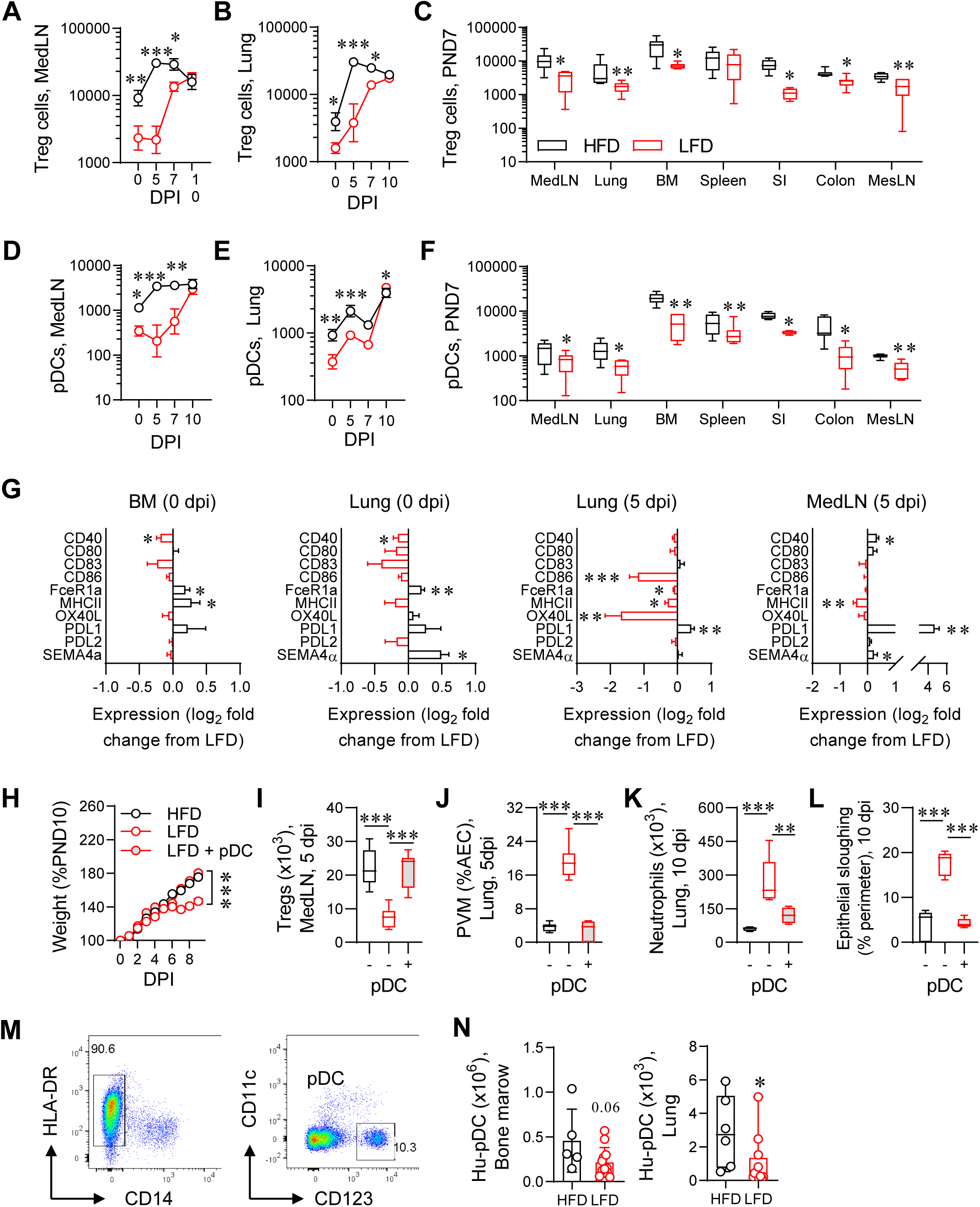
Maternal LFD decreases neonatal pDC and Treg cell expansion. (A and B) Treg cells in the MedLN (A) and lungs (B) following PVM infection. (C) Treg cells in various organs at PND7. (D and E) pDC numbers in the MedLN (D) and lungs (E) following PVM infection. (F) pDC numbers in various organs at PND7. (G) Expression of co-stimulatory molecules on pDCs in BM and lung prior to infection (left two panels), and lung and MedLN at 5 dpi (right two panels) represented as the log_2_ fold change (L2FC) from LFD-reared neonates. Markers with negative L2FC are more highly expressed in LFD-reared pDCs, while those with positive L2FC are higher in HFD-reared pDCs. (H) Weight gain in PVM infected neonates (study design of pDC adoptive transfer study shown in Fig.S6A). (I) Treg cells in the MedLN at 5 dpi. (J) PVM immunoreactivity as a percentage of AECs at 5 dpi. (K) Lung neutrophils at 10 dpi. (L) Epithelial sloughing at 10 dpi. (M) Representative FACS plots showing human pDC in the BM of humanized mice (full gating strategy shown in Fig.S6G). (N) Human pDC numbers in BM and lung. Black boxes or circles represents HFD-reared pups, red boxes or circles represent LFD-reared pups unless mentioned otherwise. Data are represented as mean ± SEM (A, B, D, E, G and H) or box-and-whisker plots (median, quartiles, and range) (C, F, I-L and N), n=5-13 mice per group. *P* values (* p<0.05, ** p<0.01, *** p<0.001) were derived by Mann Whitney U test, comparing HFD-reared with LFD-reared mice (C, F, G and N) and one-way (I-L) or two-way ANOVA (A, B, D, E and H) followed by Sidak’s or Tukey’s multiple comparisons test as appropriate.

To directly implicate pDC in conferring protection, we adoptively transferred FACS-purified pDC (isolated from HFD-reared pups) to recipient LFD-reared pups during the acute PVM infection (Fig.S6A). Of note, donor pDCs markedly increased Treg cell numbers, decreased viral load, and decreased LRI severity (Fig 2H-L and Fig.S6B-D). Similarly, the adoptive transfer of Treg cells (isolated from HFD-reared pups) to recipient LFD-reared pups ameliorated LRI severity (Fig.S6E-F), confirming the importance of these two immunoregulatory cell types. To extend our findings to the development of human pDCs, we grafted CD34+ hematopoietic stem cells into HFD- and LFD-reared pups to generate humanized mice and assessed pDC numbers at PND25. Human CD45^+^ cell numbers in BM and lung were not affected by maternal diet, however, human pDCs were numerically lower in the BM and significantly diminished in the lungs of LFD- relative to HFD-reared pups (Fig.2M-N and Fig.S6G-H). Collectively, our data indicate that maternal intake of dietary fibre positively impacts DC and Treg cell homoeostasis in infancy.

### Maternal diet influences the milk and infant microbiota, SCFA levels, and LRI severity

The fecal microbiome of mothers differed significantly between diets throughout pregnancy (Fig. S7A). In addition, dietary fibre content was positively associated with levels of propionate and acetate in the mother’s feces (Fig.S7B). Similarly, for pups at PND7, the HFD diet altered fecal microbial community composition (Fig.3A), and increased SCFA concentrations, particularly propionate and butyrate, in feces and serum (Fig.3B and Fig.S7C-D). By switching the newly-born pups of HFD-fed to LFD-fed mothers, and *vice versa* (i.e. cross-fostering), we found that the neonatal microbiome and SCFA profiles were strongly influenced by the diet of the fostering mother post-birth (Fig.S7E-G). Critically, switching pups from LFD- to HFD-fed mothers conferred protection against sLRI, whereas the opposite led to disease (Fig.3C-D and Fig.S7H-I).

**Figure 3.**
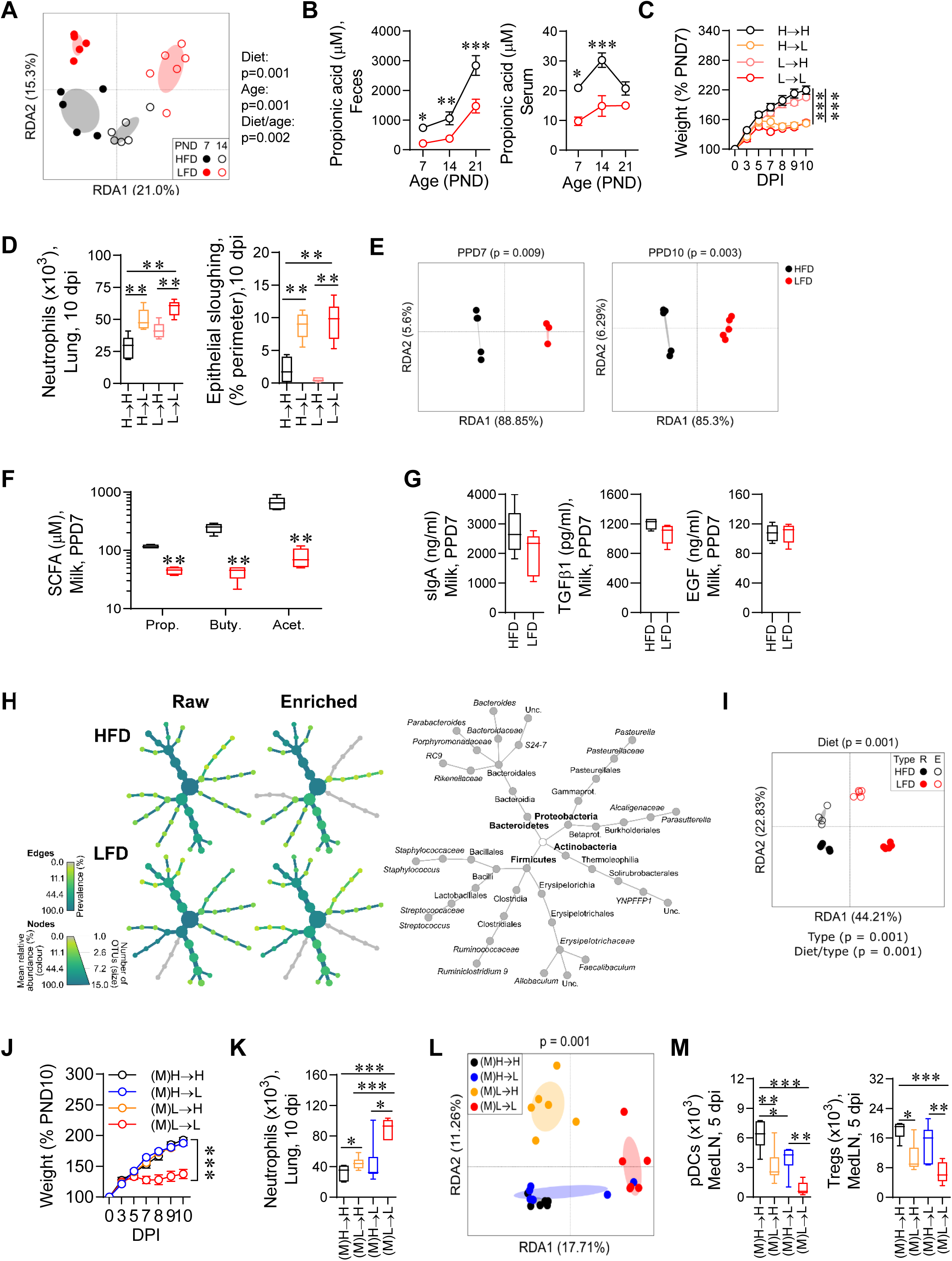
Maternal diet influences milk and infant microbiota, SCFA levels and severity of LRI. (A) RDA ordination highlighting differences in the composition of fecal microbiomes of HFD and LFD-reared pups at PND7 and PND14. The circles are samples and the ellipses represent one standard deviation around group centroid. The p values derive from a PERMANOVA model of the main and interactive effects of diet and age. (B) Propionic acid concentration in feces (left panel) and serum (right panel). (C) Weight gain in PVM infected neonates. (D) Lung neutrophils (left panel) and epithelial sloughing (right panel) at 10 dpi. (E) RDA ordination highlighting the effect of diet on the composition of milk microbiomes at PND7 (left) and at PND10 (right) (p values represent the main effect of diet according to PERMANOVA). Circles are individual samples and ellipses represent the standard deviation around the centroid of each group. (F) SCFAs concentration in milk at PPD7. (G) sIgA, TGF-β1, and EGF concentration in milk at (postpartum day) PPD7. (H) Heat trees of milk microbial communities (left panel). Trees are rooted to kingdom (Bacteria), and are aligned to the taxonomic structure of the grey skeleton tree (right panel). Edge color, node color, and node size represent the prevalence, mean relative abundance, and number of associated OTUs, respectively, for each taxa across each replicate set. (I) RDA ordination highlighting differences in the composition of microbial communities between raw (Type R) and enriched (Type E) milk microbiota from HFD- and LFD-fed mothers (p=0.001, PERMANOVA) at PPD10. Circles are individual samples and ellipses represent the standard deviation around the centroid of each group. (J) Weight gain in PVM infected pups post enriched milk microbiota transplantation (MMT; full study design in Fig.S8J). (K) Lung neutrophils (left panel) and epithelial sloughing (right panel) at 10 dpi. (L) RDA ordination highlighting differences in the composition of MMT pup fecal microbiomes (p=0.001, PERMANOVA) at 5 dpi. The circles are samples and the ellipses represent one standard deviation around group centroid. (M) pDCs (left panel) and Treg cells (right panel) in MedLN at 5 dpi. Black boxes or circles represents HFD-reared pups, red boxes or circles represent LFD-reared pups unless stated otherwise. Data are represented as mean ± SEM (B, C and J) or box-and-whisker plots (median, quartiles, and range) (D, F, G, J, K and M), n=4-15 mice per group. *P* values (* p<0.05, ** p<0.01, *** p<0.001) were derived by Mann Whitney U test, comparing HFD-reared with LFD- reared mice (F and G) and one-way (D, K and M) or two-way (B, C and J) ANOVA followed by Sidak’s or Tukey’s multiple comparisons tests as appropriate.

Next, we sought to assess the likely paths of transmission of microbes/effectors from mothers to infants. While mice tend not to engage in coprophagy until PND17 ^41^, we excluded this as a means of conferring protection to pups by performing fecal microbiome transplants (FMTs). Notably, oral gavage of HFD-fed mothers’ feces to susceptible LFD-reared pups failed to reduce LRI severity (Fig.S8A-E), whereas the feces of HFD-reared pups conferred protection against sLRI (lowering viral load, lung neutrophils, and epithelial sloughing; Fig.S8A,F-I). This suggests that the transfer of protective microbiota from mother to infant is not mediated through fecal exposure.

Hence, we next considered breastmilk as a potential source of protective microbes/effectors ^42^. We found that the composition of milk microbiomes and SCFA concentrations differed significantly in response to variation in maternal diet (Fig. 3E-F). In contrast, potential effectors, including concentrations of epidermal growth factor (EGF), TGF-β, and secretory IgA were similar, irrespective of maternal diet (Fig.3G). To more definitively attribute protection to milk microbes, we enriched milk microbiomes in anaerobic culture, confirmed that these enrichments were representative of raw milk (raw vs. enrichment, Mantel p = 0.016), and then gavaged the spin-purified samples into susceptible pups prior to PVM inoculation (Fig.S8J). Post-enrichment, the composition of milk microbiomes remained significantly different between maternal diets, with ≥75% of the dominant taxa in raw milk successfully recovered (Fig.3H-I). Importantly, LFD-reared pups were protected against sLRI following transfer of HFD-enriched milk microbes, and this was associated with an altered gut microbiome and increases in fecal and serum propionate (Fig.3J-L and Fig.S8K-M). In contrast, transfer of LFD-enriched milk microbes to HFD-reared pups did not significantly influence weight gain, and had only minor effects on other pathologies, potentially due to the continued supply of beneficial milk microbes from the HFD-fed mother. In these milk microbiota transplantation (MMT) studies and after cross-fostering, resistance to sLRI was again associated with an increase in both pDC and Treg cell numbers, consistent with an important protective role for these two immunoregulatory cell types (Fig.3M and Fig.S7I).

### Bacterial isolates from the milk of HFD-fed mothers or propionate supplementation confer protection against sLRI

Next, we questioned whether individual isolates from the enriched milk microbiota of HFD-fed mothers would promote resistance. Colonies formed on agar plates inoculated with enriched milk exhibited two dominant morphotypes. Individual colonies from each of these morphotypes were isolated and then sequenced. Genome sequencing identified the isolates as three *Enterobacter steigerwaltii* and five *Enterococcus_A avium* cultures (Fig.S9A). Enterobacter and Enterococcus have previously been reported in milk ^43–46^, and *in silico* metabolic reconstruction revealed that both *Enterobacter steigerwaltii* and *Enterococcus_A avium* possess multiple propionate pathways originating from both carbohydrate and amino acid degradation (Fig.S9B). The ability of these isolates to produce propionate was confirmed via *in vitro* culturing experiments (Fig.S9C).

Importantly, inoculation of LFD-reared pups with individual isolates conferred protection against sLRI (Fig.4A-B and Fig.S10A-C), and was associated with an increase in pDCs, Treg cells, and fecal propionate levels (Fig.4C-D and Fig.S10D-E). The relative abundances of OTUs most closely affiliated with the isolate genomes increased in response to isolate transplantation (Fig.S10F). In addition, introduction of the isolates led to significant changes in the resident gut microbiome (Fig.S10G), as well as their predicted propionate-associated gene profiles (p<0.001). This suggests that the reduction in morbidity may be directly attributable to the isolate, the isolate-induced changes to the microbiome, or a combination of the two.

**Figure 4.**
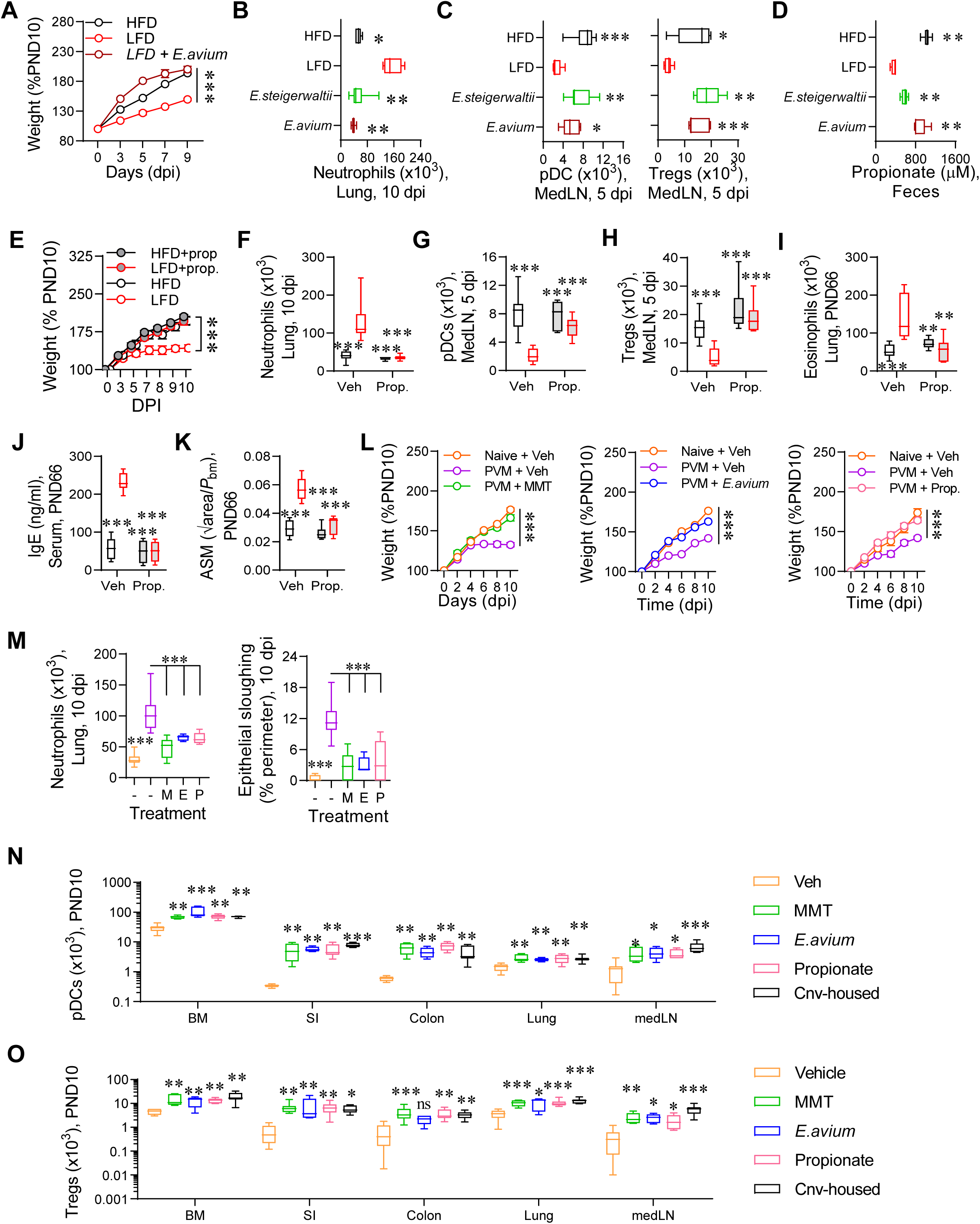
Bacterial isolates form milk of HFD-fed mothers confer protection against sLRI. (A) Weight gain in PVM-infected neonates following transplantation of *Enterobacter steigerwaltii* or *Enterococcus_A avium*. (B) Neutrophils in lung at 10 dpi. (C) pDCs and Treg cells in MedLN at 5 dpi. (D) Propionic acid concentration in feces at PND10. (E) Weight gain in PVM-infected LFD-reared pups supplemented with propionate (full study design in Fig.S11a) (F) Lung neutrophils at 10 dpi. (G and H) pDCs (G) and Treg cells (H) in MedLN at 5 dpi. (I) Lung eosinophils at PND66. (J) Total IgE concentration in serum at PND66. (K) ASM area at PND66. (L) Weight gain in PVM infected GF neonates. (M) Lung neutrophils (left panel) and epithelial sloughing (right panel) at 10 dpi. (N and O) pDC (N) and Treg cell (O) numbers in BM, small intestine, colon, lung, and MedLN in naïve mice at PND10. Black boxes or circles represents HFD-reared pups, red boxes or circles represent LFD-reared pups and other treatment groups are labeled in the graphs. Data are represented as mean ± SEM (A, E and L) or box-and-whisker plots (median, quartiles, and range) (B-D, F-K, and M-O), n=5-12 mice per group. *P* values (* p<0.05, ** p<0.01, *** p<0.001) were derived by one-way (B-D and M-O) or two-way (A, and E-L) ANOVA followed by Tukey’s multiple comparisons test. The LFD-reared group was compared with other treatment groups (A-K), PVM infected vehicle treated GF group was compared with other treatment groups (L and M) and naive vehicle treated GF group was compared with other treatment groups within individual compartment (N and O).

As protection from sLRI following cross-fostering, MMT or monobacterial transplantation was more consistently associated with increased levels of propionate (rather than acetate or butyrate), we hypothesised that propionate would confer protection to LFD-reared pups. As anticipated, propionate supplementation (oral gavage) restored propionate levels and diminished LRI severity (Fig.4E-F and Fig.S11A-E), and this was associated with an increase in pDCs, cDCs, and Treg cells in the medLNs and lungs (Fig.4G-H and Fig.S11F-H). In support of a causal association between sLRI in early-life and subsequent asthma, treatment with propionate in early-life averted the development of type-2 inflammation, IgE production, and airway remodelling in later-life (Fig.4I-K and Fig.S11I-L). To confirm the importance of the gut microbiome to lung immunity, we next employed neonatal germ-free (GF) mice, and assessed the effect of (i) enriched MMT from a HFD-fed mother (‘M’), (ii) the *E. avium* isolate (‘E’), or (iii) propionate supplementation (‘P’) on LRI severity. Despite being reared to HFD-fed mothers, GF mice developed sLRI. However, oral gavage with any of the three treatments ameliorated LRI severity and increased fecal propionate levels (Fig. 4l-M and Fig.S11M). At steady state, pDC and Treg cell numbers were significantly lower in the BM, SI, colon, lung, and medLN of GF compared to conventionally-housed neonatal mice (Fig.4N-O). However, this multi-tissue defect was rescued by MMT, mono-colonisation with *E. avium*, or propionate supplementation (Fig.4N-O). As the maternal LFD affected pDC, cDC1, and cDC2 in the offspring, we next assessed whether propionate-mediated protection from sLRI would be ablated in pDC-diptheria toxin receptor ^22, 47^ transgenic pups reared to LFD-fed mothers. The induced depletion of pDCs reversed the beneficial effects of propionate supplementation on LRI severity (with the exception of epithelial sloughing), and this was associated with an inability to increase Treg cells in the medLN (Fig.S12A-D). Together, these findings indicate that the microbiome-mediated protection from sLRI occurs via the expansion and recruitment of Treg cells to the lung, and that this is indirectly mediated via signals provided by pDCs. The effect of Flt3L-deficiency on the contribution of cDC1 and cDC2 to host defence against PVM remains to be formally assessed.

### Maternal diet affects Flt3L expression in the neonatal intestine and bone marrow

Propionate has been linked to DC hematopoiesis ^17^, however the underlying mechanisms remain obscure ^48^. As the hematopoietic cytokine Flt3L is integral to pDC development ^49^, we measured *Flt3L* gene expression in the organs where pDC numbers were affected by maternal LFD. This revealed lower *Flt3L* expression in the SI, colon and BM (Fig.5A). Immunofluorescent staining of the SI, colon and BM in cross-section confirmed these findings at the protein level (Fig.5B). Pre-incubation of anti-Flt3L with exogenous Flt3L completely ablated the immunofluorescent signal, validating the specificity of the fluorescent signal (Fig.S13A). The attenuated Flt3L expression in the SI of LFD-reared pups was confirmed by ELISA (Fig.5C). As Flt3L (red) was expressed on the apical surface of villin+ (green) intestinal epithelial cells (IECs), and evident in the gut lumen (Fig.5B), we measured Flt3L in fecal pellets. Notably, Flt3L levels were lower in the stool of LFD-reared pups (Fig.5D). By PND42, however, Flt3L expression was barely detectable in IECs or feces, in contrast to BM cell Flt3L expression, which increased with age (Fig.5D and Fig.S13B-C). As RSV bronchiolitis primarily affects infants aged <4 months ^32^, we assessed Flt3L expression in the gut and serum throughout the neonatal period. In HFD-reared pups, fecal Flt3L levels increased in the first week of life, before waning to adult levels at around PND21 (Fig.5E), a pattern that was repeated in the serum (Fig.5F). Indeed, the tight correlation in Flt3L levels between the two compartments (Fig.5G) suggests that the gut is the primary source of serum Flt3L in early life. Critically, the postnatal elevation in fecal and serum Flt3L expression was completely absent in LFD-reared pups (Fig.5E-F), consistent with the aberrant expansion of neonatal DCs.

**Figure 5.**
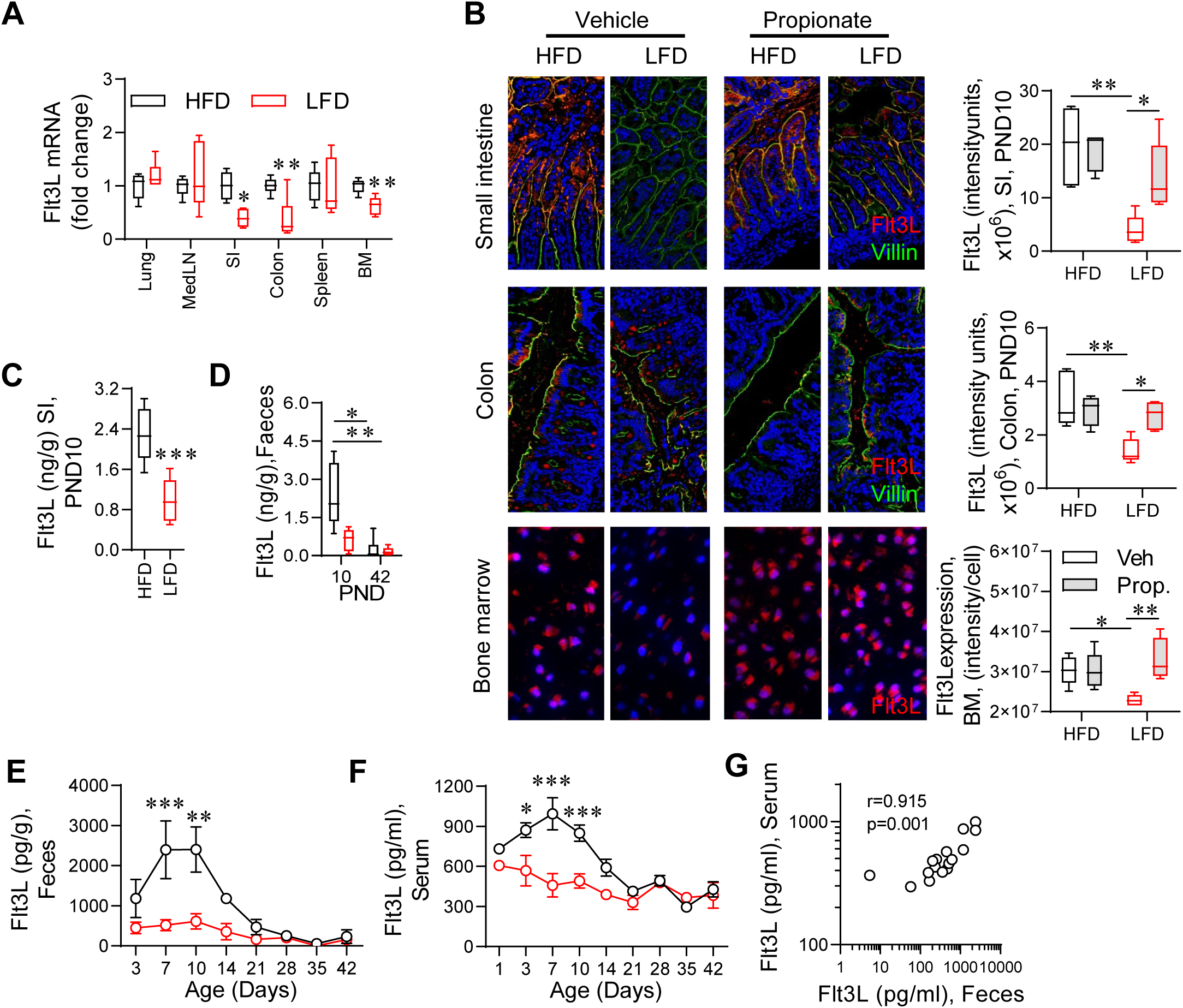
Maternal diet affects neonatal Flt3L expression in the intestine. (A) *Flt3L* gene expression in various tissues. (B) Flt3L expression in small intestine colon and bone marrow (BM). Flt3L immunofluorescence (red), villin (gut epithelial marker; green) and DAPI counterstain (blue). Quantification of Flt3L expression (intensity/cell) shown on the right hand side. (C and D) Flt3L production in small intestinal homogenate (C) and fecal pellet (D) at PND10. (E and F) Flt3L concentration in feces (E) and serum (F). (G) Correlation of feces and serum Flt3L concentrations. Black boxes represent HFD-reared pups, red boxes represent LFD-reared pups, grey shaded boxes represent propionate supplementation. Data are represented as mean ± SEM (E and F) or box-and-whisker plots (median, quartiles, and range) (A-D), n=4-8 mice per group. *P* values (* p<0.05, ** p<0.01, *** p<0.001) were derived by Mann Whitney U test, comparing HFD-reared with LFD- reared mice (A, C and D) or two-way ANOVA followed by Sidak’s (E and F) or Tukey’s (B) multiple comparisons test as appropriate.

### ADAM10 inhibition decreases soluble Flt3L and predisposes to sLRI

Surface-expressed Flt3L is biologically active, however, its cleavage by enzymes such as ADAM10 ^50^ enables soluble Flt3L to act as a cytokine. Consistent with the immunofluorescence images (Fig.5B), flow cytometric detection of Flt3L expression by IECs did not require permeabilisation, in stark contrast to BM cells, indicating that IECs express surface-bound Flt3L (Fig.6A-C). Hence, we reasoned that ADAM10 inhibition in HFD-reared pups would decrease Flt3L levels and increase susceptibility to sLRI. As hypothesized, oral delivery of an ADAM10 inhibitor increased surface-bound Flt3L in SI IECs, but not BM cells (Fig.S13D-E). Furthermore, the ADAM10 inhibitor lowered Flt3L levels in the feces, gut lumen, and serum (Fig.6D-E), with consequent decreases in BM and lung associated pDCs (Fig.6F and Fig.S13G). In response to acute infection, ADAM10 inhibition ablated pDC and Treg cell expansion (Fig.6G and Fig.S13H), predisposing otherwise healthy HFD-reared pups to a sLRI (Fig.6H-I and Fig.S13I). Importantly, ADAM10 affects a broad array of proteins ^51^, and hence there are limitations to the systemic use of an ADAM10 inhibitor, however these data are consistent with the notion that gut-derived Flt3L promotes postnatal DC hematopoiesis and confers protection against sLRI.

**Figure 6.**
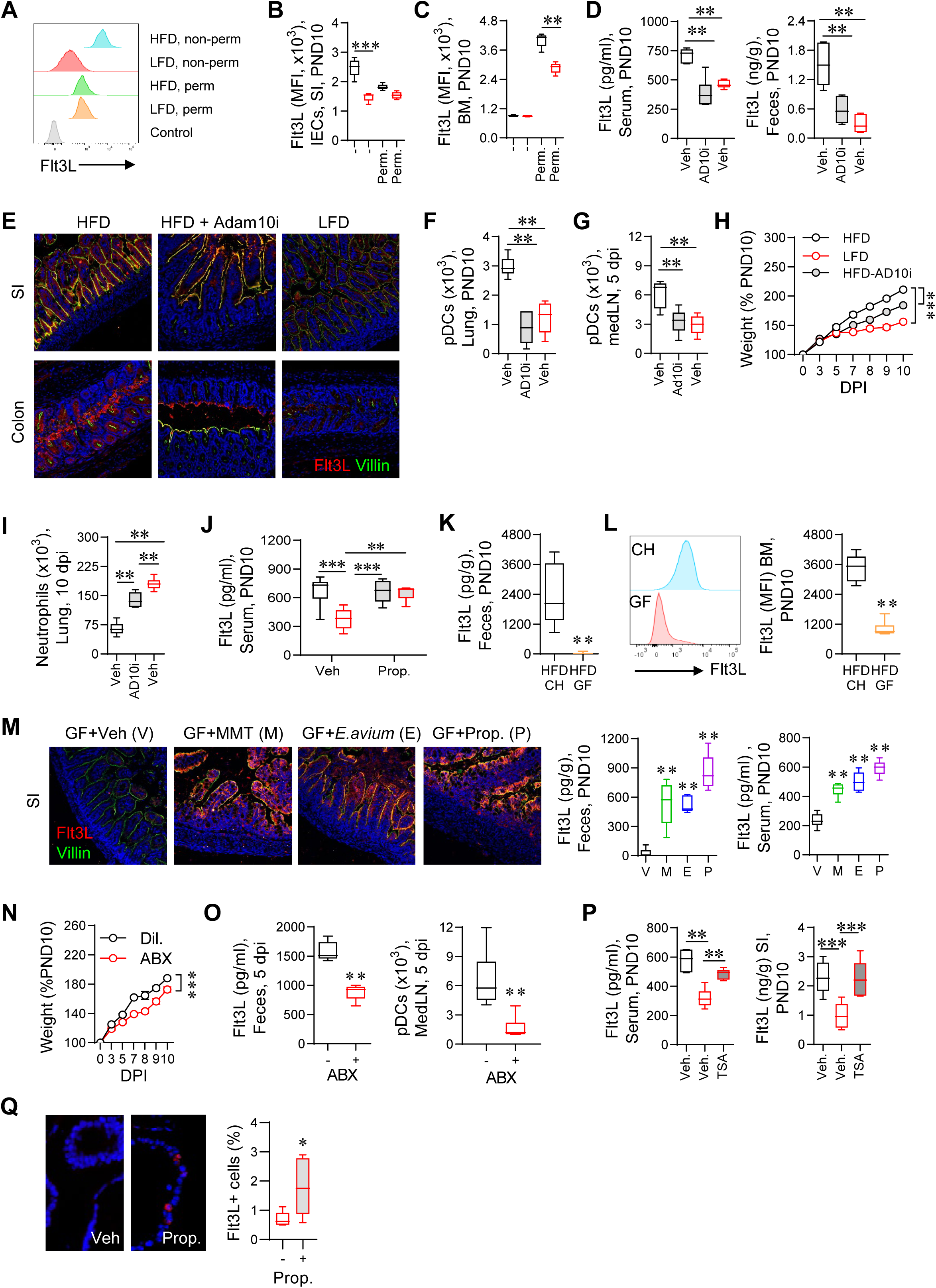
Inhibition of ADAM10 decreases soluble Flt3L and predisposes to sLRI. (A-C) Flt3L expression in intestinal epithelial cells (IECs) (A and B) and BM cells (C) at PND10. (D and E) Flt3L concentration in serum (left panel) and feces (right panel)(D), and small intestine (SI, upper panel) and colon (lower panel) at PND10 following ADAM10 inhibition (AD10i; full study design in Fig.S13D); Flt3L (red), villin (epithelial protein, green) and DAPI (blue). (F) pDCs in lung at PND10. (G) pDCs in medLN at 5 dpi. (H) Weight gain in virus-infected pups. (I) Lung neutrophils at 10 dpi. (J) Serum Flt3L concentration at PND10. (K-M) Flt3L production in GF and conventionally-housed (CH) mice in feces and BM cells, detected by ELISA and flow cytometry respectively. (K) Flt3L expression in feces. (L) Flt3L expression (MFI) in BM cells. (M) Flt3L expression in neonatal SI at PND10 detected by immunofluorescent microscopy and quantification of Flt3L in feces and serum following treatments as indicated. (N) Weight gain in PVM-infected neonates following maternal antibiotic (ABX) exposure. (O) Fecal Flt3L production (left panel) and pDCs in MedLN (right panel) at 5 dpi. (P) Effect of TSA (full study design in Fig. S13N) on Flt3L concentration in serum (left panel) and SI homogenate (right panel) at PND10. (Q) Effect of propionate on Flt3L expression by enteroids generated from SI of LFD-reared pups assessed by immunohistochemistry (left) and flow cytometry (right). Black boxes or circles represents HFD-reared pups, red boxes or circles represent LFD-reared pups. Data are represented as mean ± SEM (H, N) or box-and-whisker plots (median, quartiles, and range) (B-D, F, G, I-M, O-Q), n=5-8 mice per group. *P* values (* p<0.05, ** p<0.01, *** p<0.001) were derived by Mann Whitney U test, comparing CH with GF (K and L), vehicle- and antibiotic- treated (N and O) and vehicle- and propionate-treated enteroids (Q), and one-way (B-D, F, G, I, M, P) and two-way (H, J, and N) ANOVA followed by Tukey’s multiple comparisons test.

We next assessed serum Flt3L levels in the various intervention studies. In all cases, e.g. cross-fostering, MMT, bacterial monotherapy (Fig.S13J-L) and propionate supplementation (Fig.6J), serum Flt3L expression negatively associated with disease severity. In neonatal GF mice, Flt3L levels were ablated in IECs, feces, and serum (Fig.6K-M), and restored following MMT, bacterial monotherapy, or propionate supplementation, further highlighting the importance of the milk microbiota to the assembling enteric microbiota and downstream production of bioactive metabolites. Pups reared to antibiotic-treated mothers, which were predisposed to sLRI, expressed low levels of Flt3L and had attenuated numbers of pDC and Treg cells (Fig.6N-O and Fig.S13M). As propionate can mediate its anti-inflammatory effects through the inhibition of histone deacetylases (HDACs)^18^, we treated mice with the pan-HDAC inhibitor, TSA (Fig.S13N).

Following oral gavage with TSA, Flt3L expression in the serum and homogenized SI of LFD-reared pups was restored to levels observed in HFD-reared pups (Fig.6P), implicating a mechanism of action by which propionate regulates Flt3L in IECs. To assess whether propionate acts directly on IECs, we grew enteroids from the SI of LFD-reared pups ^52^. After a three day culture in the presence of propionate, the gut enteroids expressed significantly higher levels of Flt3L (Fig.6Q), implicating a direct effect. Collectively, these observations indicate the existence of a postnatal burst of Flt3L expression, affected by fermentable fibre in the maternal diet, which regulates the neonatal development of pDCs and Treg cells.

### Anti-Flt3L ablates propionate-induced protection against sLRI

To establish whether Flt3L is the molecular factor by which propionate promotes the development of pDCs and confers protection against sLRI, we next treated the propionate supplemented LFD- reared pups with anti-Flt3L or an isotype-matched control (Fig.S14A). Anti-Flt3L abolished the propionate-induced increases in pDCs, cDC1s, and cDC2s in the BM, medLN, lung, SI and colon (Fig. 7A-C and Fig.S14B-D). Although Treg cells were similarly ablated in the BM, medLN and lung, anti-Flt3L had no effect on propionate-induced increases in gut Treg cells (Fig.7D-F and Fig.S14E). This indicates that the propionate-induced expression of gut-derived Flt3L facilitates pDC-mediated expansion of extra-enteric Treg cells, and suggest that Flt3L neutralization would abrogate the protective effect of propionate supplementation to LFD-reared pups. As hypothesised, anti-Flt3L ablated the restorative effects of propionate supplementation on pDC and Treg cell numbers in the medLN and lungs of PVM-infected mice. This reversed the beneficial effects of propionate supplementation on LRI severity in PVM-infected LFD-reared pups, and the later development of experimental asthma, although eosinophil numbers remained attenuated (Fig.7G-P and Fig.S14F-G). Lastly, we assessed whether exogenous Flt3L would confer protection to LFD- reared pups. As predicted by the findings above, exogenous Flt3L expanded pDC numbers in the BM, lungs and MedLN prior to infection (Fig.S14H-I), and decreased the severity of LRI in LFD- reared pups (Fig.S14J-K). Taken together, these findings show that propionate confers protection against sLRI via Flt3L-mediated hematopoiesis of DCs.

**Figure 7.**
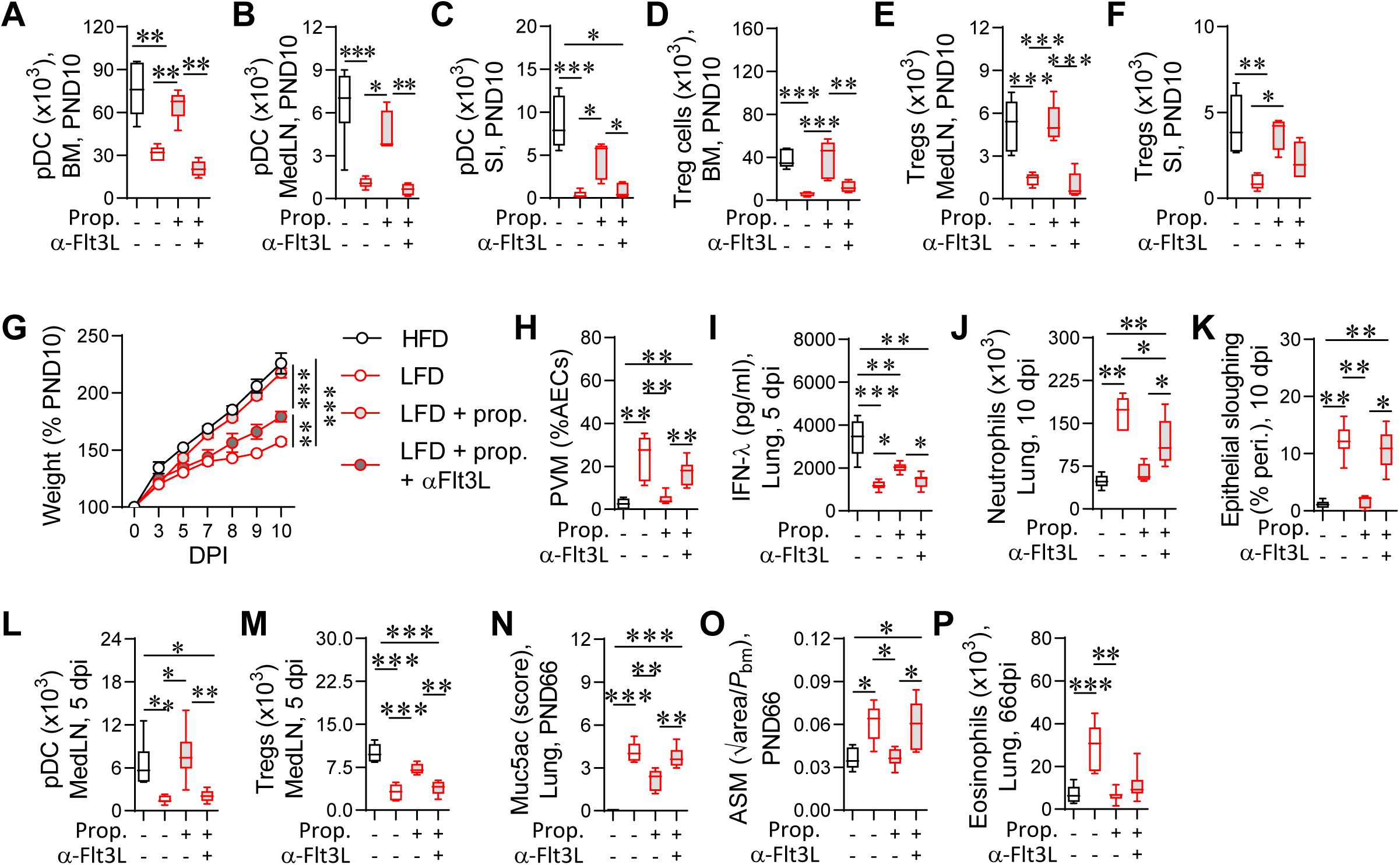
Anti-Flt3L ablates propionate-induced protection against sLRI. (A-C) pDCs numbers in BM (A), MedLN (B), and SI (C) at PND10, following propionate and anti- Flt3L Ab treatment (full study design in Fig.S14A). (D-F) Treg cells in BM (D), MedLN (E), and SI (F) at PND10. (G) Weight gain in virus-infected pups. (H) PVM immunoreactivity as a percentage of AECs at 5 dpi. (I) IFN-λ protein in lung homogenate at 5 dpi. (J) Lung neutrophils at 10 dpi. (K) Epithelial sloughing at 10 dpi. (L and M) pDCs and Treg cells in MedLN at 5 dpi. (N) Quantification of muc5ac at PND66. (O) ASM area at PND66. (P) Lung eosinophils at PND66. Black boxes represent HFD-reared pups, red boxes represent LFD-reared pups, grey shaded boxes represent propionate supplementation. Data are represented as mean ± SEM (G) or box-and-whisker plots (median, quartiles, and range) (A-F, H-P), n=5-7 mice per group. *P* values (* p<0.05, ** p<0.01, *** p<0.001) were derived by one-way (A-F, and H-P) or two-way (G) ANOVA followed by Tukey’s multiple comparisons test.

## Discussion

sLRIs during infancy are well-recognized risk factors for subsequent asthma development ^6, 7, 9, 10^, and recent evidence has identified poor maternal diet as an important determinant of infant susceptibility to these infections ^32^. Here, we demonstrate in a mouse model that fermentable fibre in the mother’s diet confers protection to her offspring against sLRI and experimental asthma by promoting the production of the DC growth factor Flt3L. As a major source of Flt3L production in early life, we identify the intestinal epithelium as a critical regulator of DC ontogeny and homeostasis. Induced by the microbial metabolite propionate, surface-bound Flt3L on IECs is cleaved by ADAM10 then enters the circulation, where after it promotes DC differentiation and the establishment of tissue DCs in readiness for immune surveillance and host defense. Critically, perturbations to this temporally choreographed wave of developmental Flt3L production massively disrupts pDC and Treg cell homeostasis, predisposing infants to sLRI and the ensuing expression of asthma-associated immunopathology.

It is well established that the production of SCFAs in the gut following the fermentation of dietary fibre increases gut Treg cell numbers. This occurs via the accumulation of tTreg cells and the local differentiation of pTreg cells, which is in part mediated by increased TGF-β expression by IECs ^18, 20, 21^. Here, we identify IEC production of Flt3L as another critical pathway, operational in the neonatal period, through which microbiome-IEC interactions promote Treg cell expansion.

Notably, this axis is particularly important for extra-intestinal (i.e. BM, lung, and medLN) Treg cell homeostasis. The production of Flt3L is induced by propionate and could be replicated via treatment with a pan-HDAC inhibitor, suggesting that other SCFAs that inhibit HDACs also regulate intestinal Flt3L expression.

Critically, in our preclinical model, propionate-induced resistance against sLRI was ablated in pDC-depleted or anti-Flt3L-treated mice, suggesting that intestinal Flt3L confers protection via the development of pDCs and their extra-enteric support of tTreg cell expansion via a Sema4a-Nrp- 1 interaction. Here, using spectral flow cytometry, we found that the perturbed neonatal microbiome affected the expression of various co-stimulatory molecules on pDCs, including Sema4a and PD-L1, which were expressed at lower levels in LFD-reared pups. This is consistent with the attenuated Treg cell expansion, although we did not formally interrogate this. Low Treg cell numbers during infancy and/or childhood has been previously identified as an important risk factor for sLRI and the development of asthma in a number of clinical settings ^53–59^, and thus future studies will need to determine whether this disruption to key DC-dependent Treg cell immunoregulatory mechanisms during the neonatal period is underpinned by Flt3L deficiency in early life. Moreover, since the absence of intestinal Flt3L in pups reared to LFD-fed mothers or following antibiotic treatment affected both pDC and cDC (and downstream Treg cell) numbers in multiple tissue compartments, disturbances to this microbiome-Flt3L pathway may also contribute to other diseases whose developmental origin commences in infancy.

Environmental and lifestyle factors in the perinatal period that influence the microbiome, such as never breastfeeding, caesarean birth, antibiotic use, and poor diet, heighten childhood susceptibility to severe infections ^26, 27^ and are increasing linked to the rising prevalence of allergic and chronic diseases ^12–14, 28, 29, 60^. In the context of asthma, maternal environmental (particularly microbial) exposures during pregnancy have been linked with variations in disease susceptibility of their offspring in both human and animal model settings ^30, 31, 61^. Collectively, these studies suggest that both antenatal and postnatal exposures to microbial products likely contribute to infant immune maturation and expression of the asthma phenotype. Here, we found that a maternal LFD affects the microbiota composition of the mother’s milk and the infant gut, lowering neonatal gut Flt3L expression. Similarly, perinatal maternal antibiotic exposure lowered Flt3L expression and predisposed the infant to sLRI. Thus, our findings identify a critical microbiome-host interaction and a downstream immunological pathway to explain how lifestyle factors that adversely influence the composition of the milk and neonatal gut microbiome, perturb immune maturation and increase disease susceptibility in early life. Although the provenance of milk-associated microorganisms remains controversial; should the entero-mammary pathway participate ^62^, then this would partly explain how antenatal exposures can impinge on events that coordinate postnatal vertical transmission (and associated neonatal immune development). Rehabilitating this pathway via manipulation of the maternal diet during pregnancy and during the pre-weaning period, or through the provision of defined tailor-made prebiotics and/or probiotics directly to infants (which we show is sufficient to confer protection), may present novel opportunities for enhancing resistance to inflammation-mediated diseases including (but not restricted to) acute LRIs, asthma and allergies. In summary, we identify Flt3L expression by IECs as a novel mechanism, operational in early-life, by which metabolic by-products of the gut microbiota promote DC and Treg cell homeostasis, and decrease susceptibility to sLRI and subsequent asthma.

## Acknowledgments

This work was supported by an NHMRC of Australia project grant awarded to S.P., P.G.D. and P.ÓC., and Mater Foundation support of K.J.R. The authors thank QIMR Berghofer Flow Cytometry Facility and Histology Facility for technical assistance, and Dr. Colonna for providing BDAC2-DTR mice.

## Author Contributions

S.P. and M.A.S. conceived the project and designed the experiments. S.P., P.G.D. and P.G.H. prepared the early manuscript drafts; all authors contributed thereafter. The majority of the *in vivo* experiments were performed by M.A.S. and R.R., with support from T.A., I.S., D.R.H., M.A.U., M.M.R., J.P.L., B.C., R.B.W., J.S., and A.B. The studies with humanized mice were performed by KR, C.W., and M.A.S. Mass spectrometry analyses performed by V.K., and T.M.W. Enteroid cultures and experiments were performed by G.E.K., T.H., and A.A. Microbiota profiling analyses performed by R.D.H. and P.G.D. Intellectual input and reagents were provided by P.ÓC., J.W.U. and M.M.

## STAR Methods

### STAR*METHODS

#### KEY RESOURCES TABLE

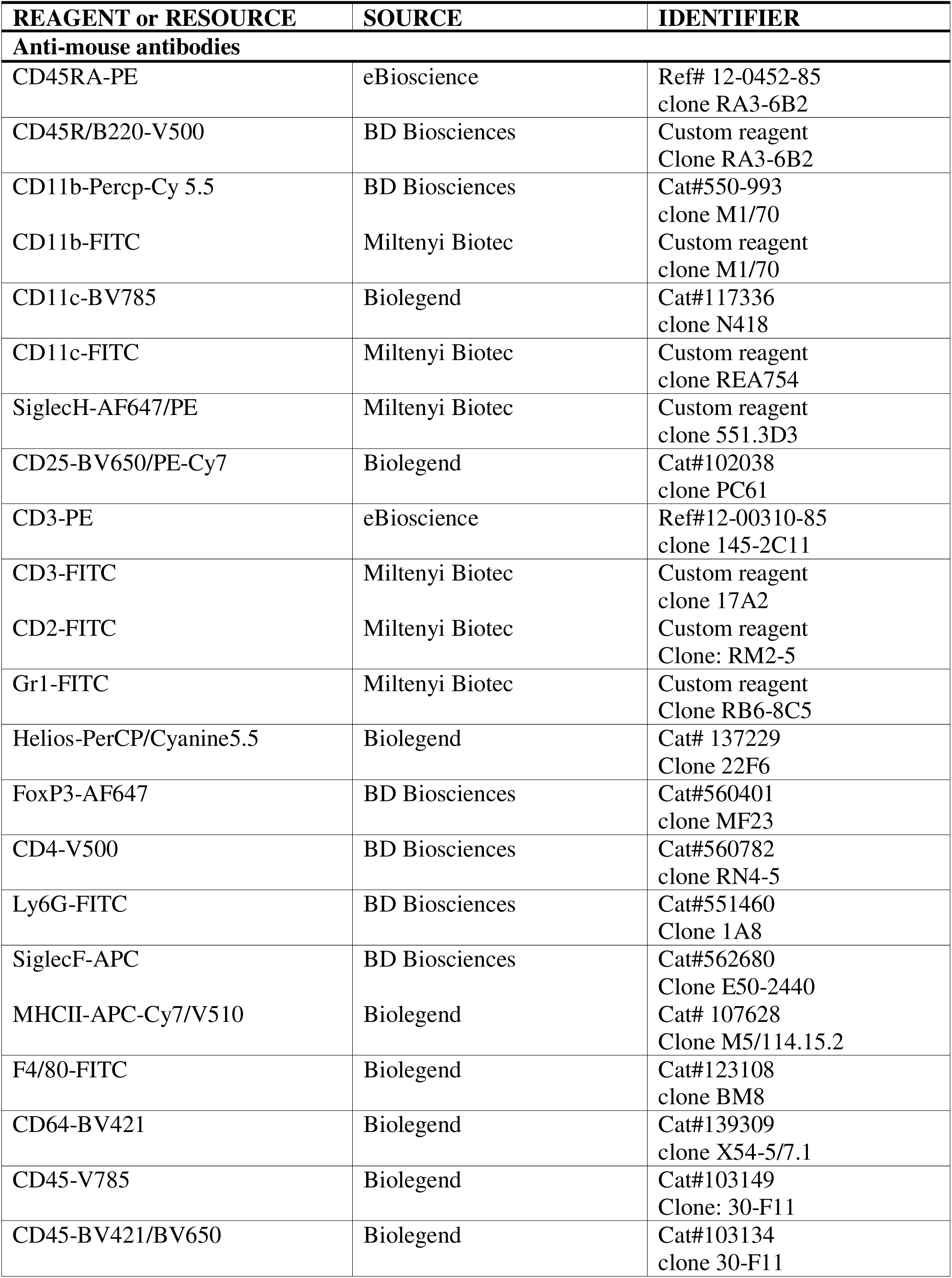
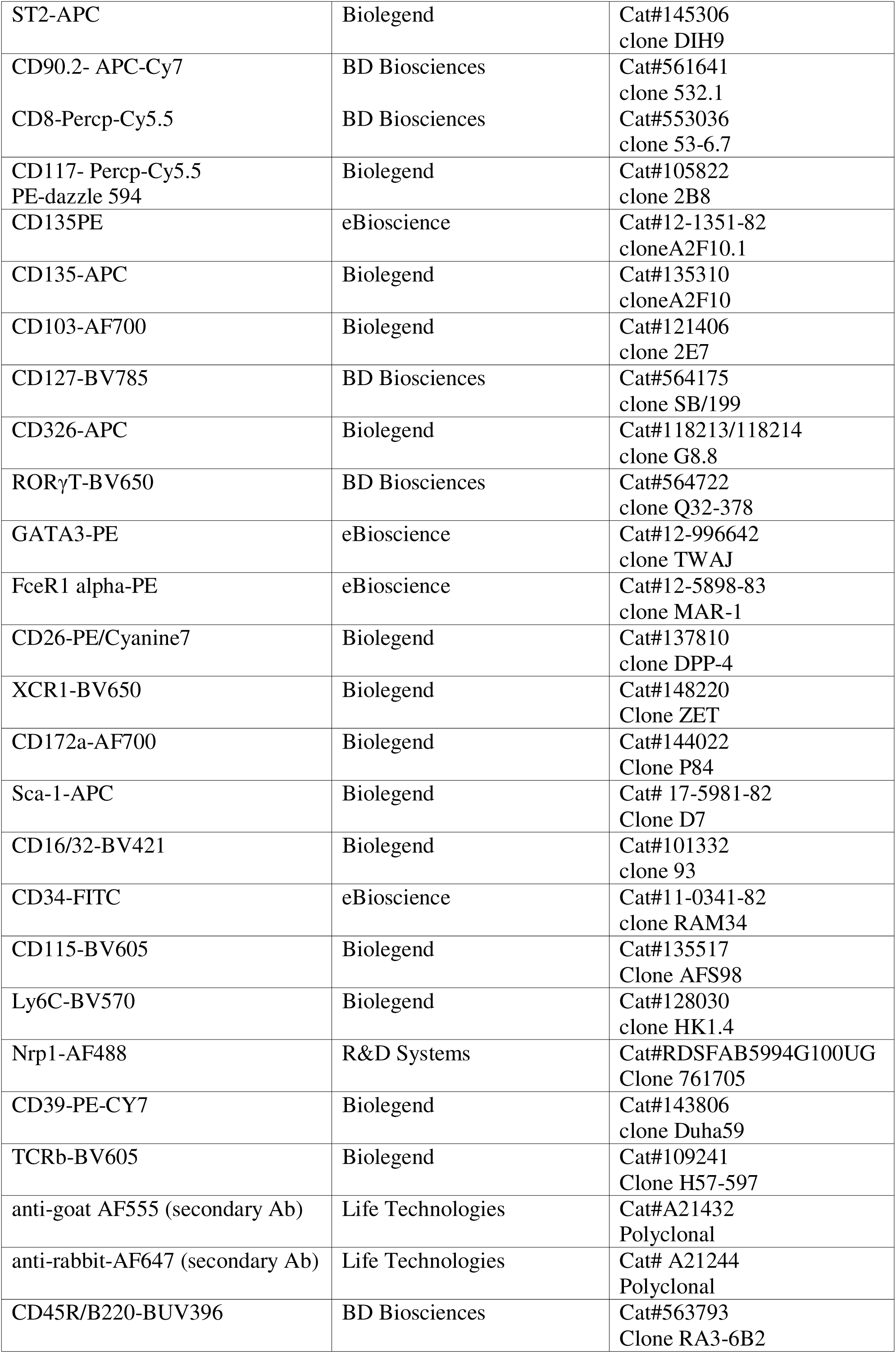
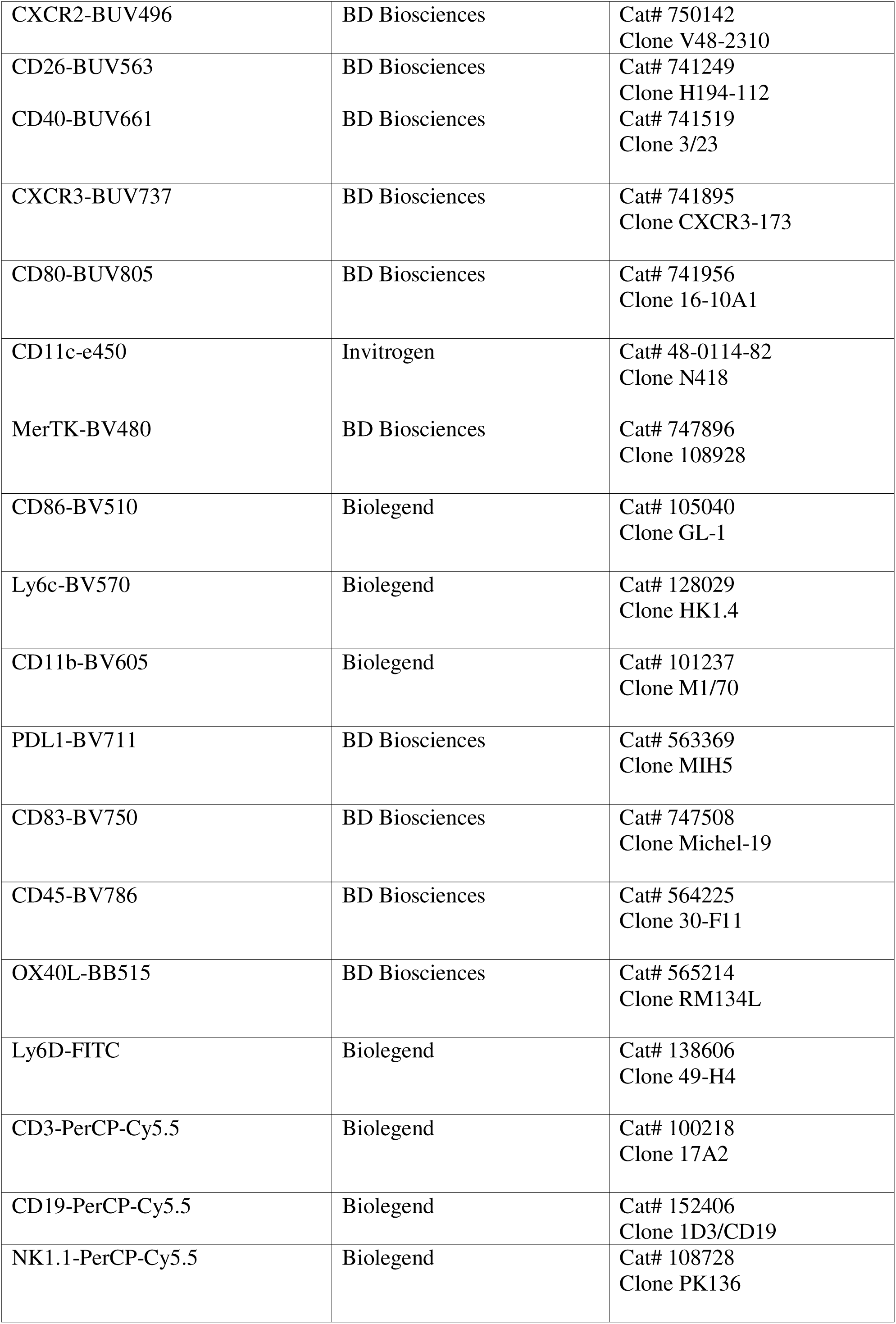
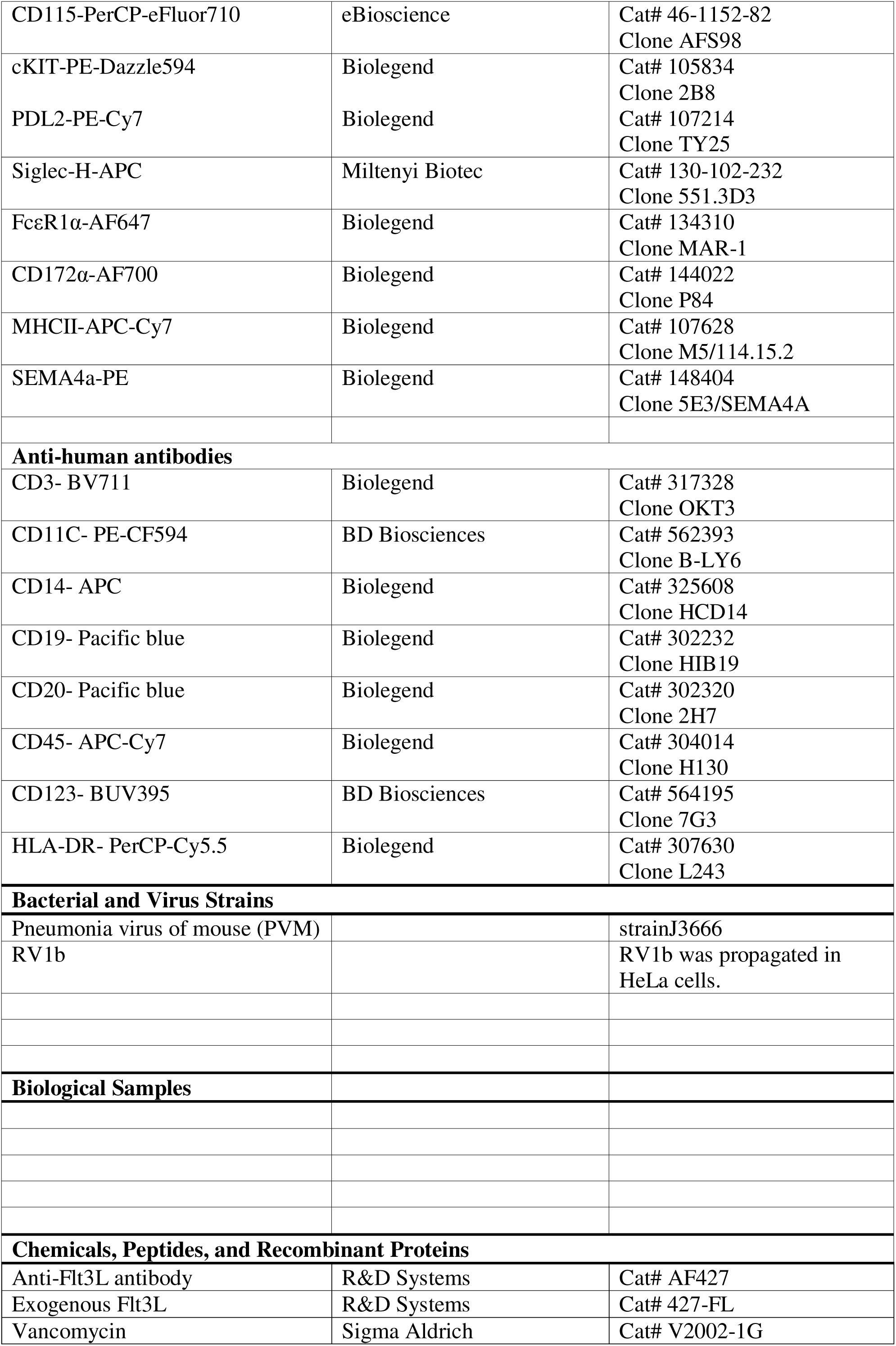
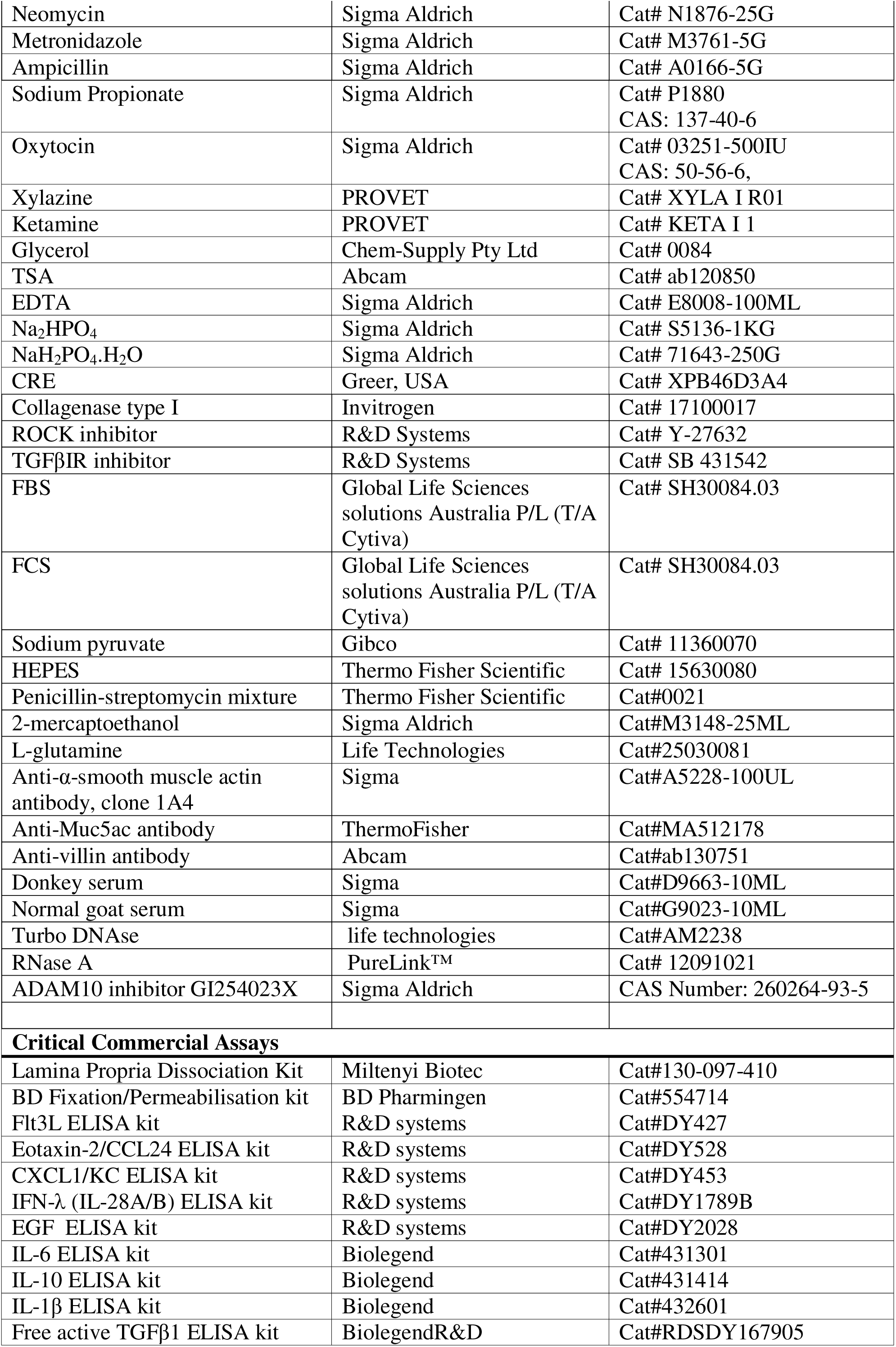
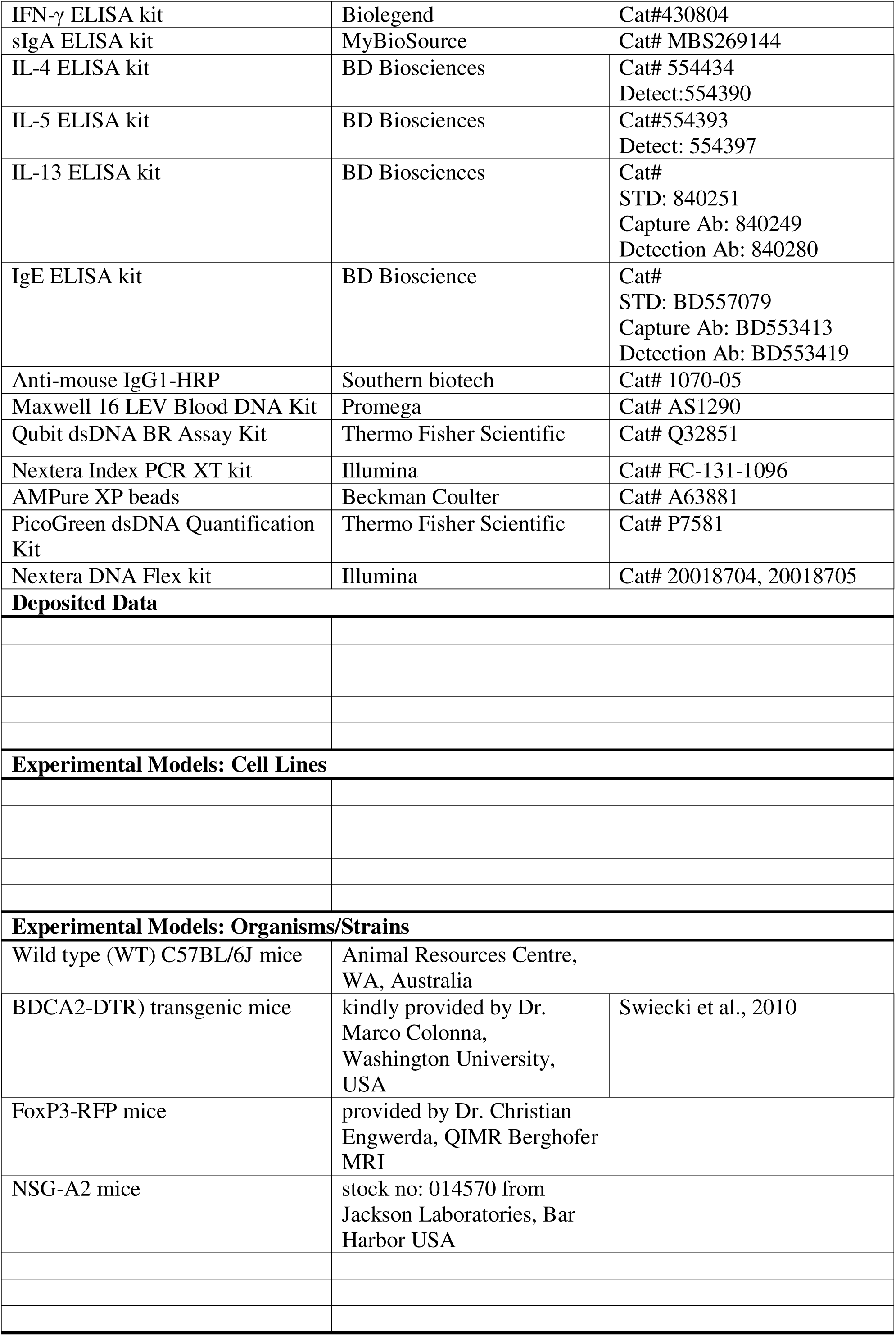
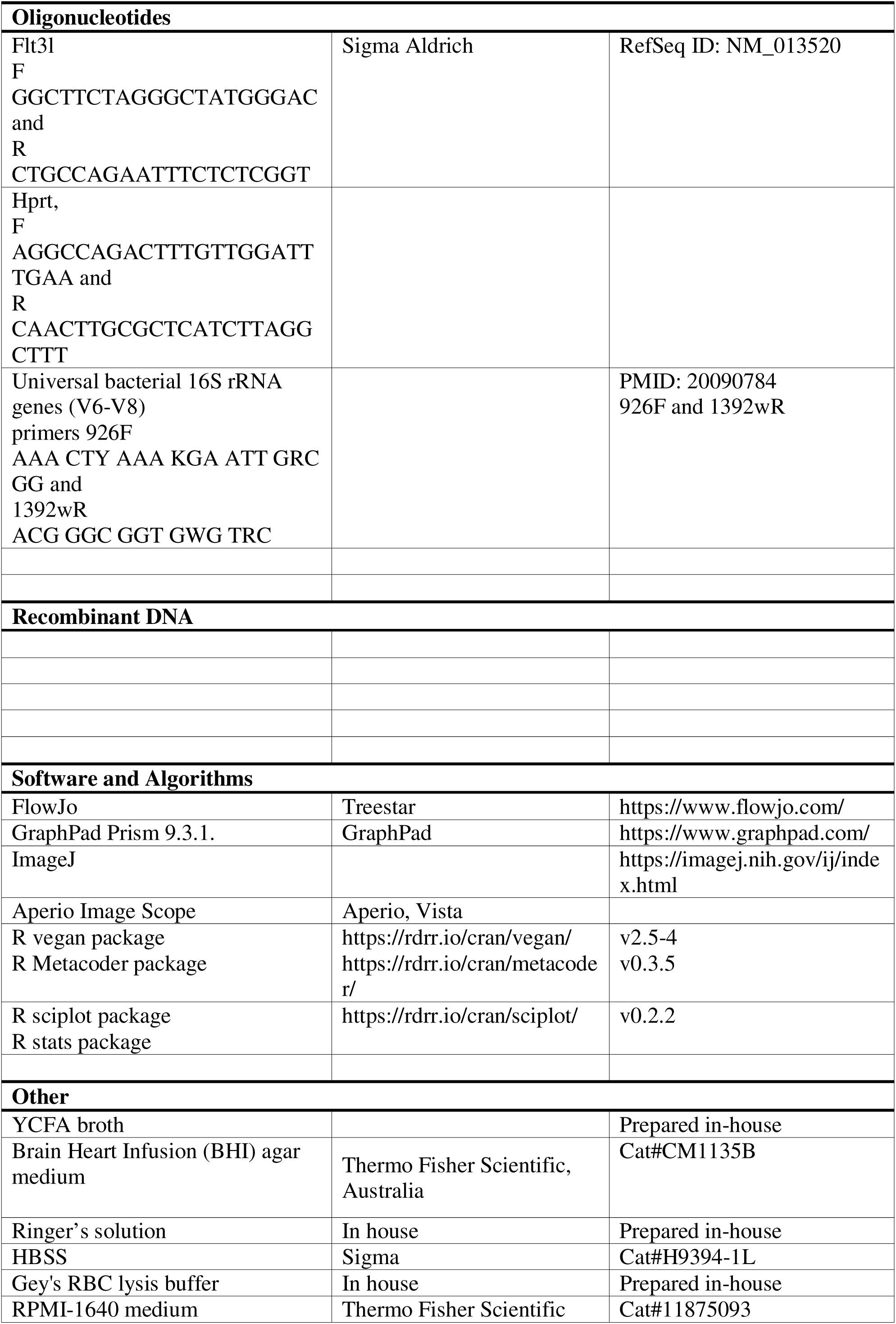
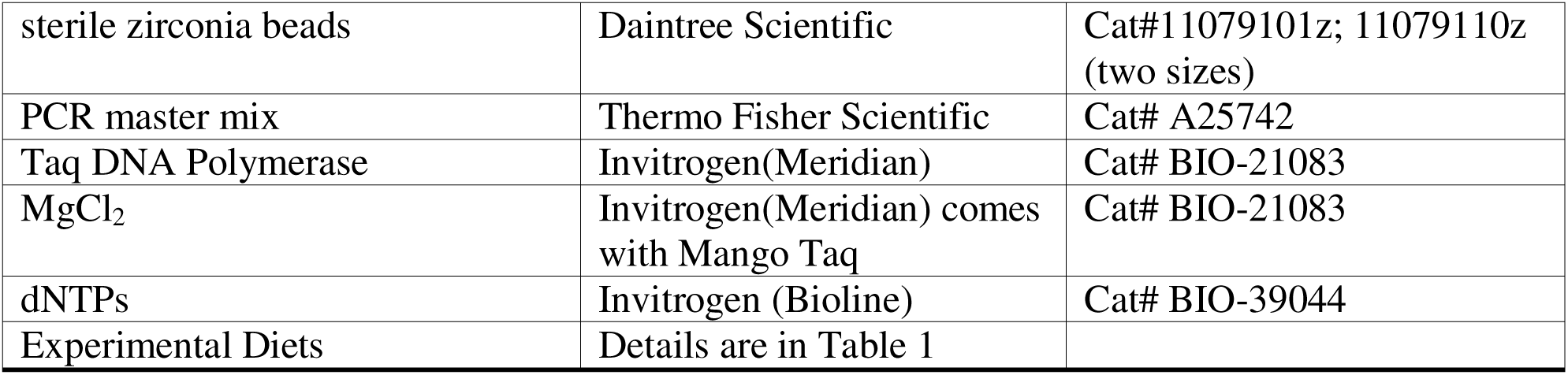

##### Contact for Reagent and Resource Sharing

Any further information and requests for resources and reagents should be directed to and will be fulfilled by the Lead Contact, Simon Phipps.

##### Mouse strains

Wildtype (WT) C57BL/6J mice (from Animal Resources Centre, WA, Australia) blood dendritic cell antigen 2-diptheria toxin receptor (BDCA2-DTR) transgenic mice (kindly provided by Dr. Marco Colonna, Washington University, USA)^47^, and FoxP3-RFP mice (provided by Dr. Christian Engwerda, QIMR Berghofer MRI) were housed at the specific pathogen free animal facility at QIMR Berghofer MRI. C57BL/6J germ-free (GF) mice and NSG-A2 mice (stock no: 014570 from Jackson Laboratories, Bar Harbor USA) were housed at the Translational Research Institute Biological Research Facility, University of Queensland. All experiments were approved by the animal ethics committees of QIMR Berghofer MRI, or University of Queensland, Australia.

##### Mouse diets

Breeding-age female and male mice were fed a high fibre diet (HFD) or low fibre diet (LFD) from 3 weeks prior to time mating and until the end of the study. In the studies of experimental asthma, the neonatal mice were weaned at 3 weeks of age, and fed the same diet as their parents had received. After plugging the studs were removed. HFD and LFD chow was purchased from Specialty Feeds (Western Australia, Australia). The composition of the diets is shown in Table 1.

**Table 1.** Experimental Diets.

| Composition | High-fibre diet | Low-fibre diet |
| --- | --- | --- |
| Protein | 19.4% | 19.4% |
| Total Fat | 7.0% | 7.0% |
| Crude Fibre | 4.7% | 0.0% |
| Acid Detergent Fibre | 4.7% | 0.0% |
| Neutral Detergent Fibre | 0.0% | 0.0% |
| Cellulose | 50 g/Kg | 0.0% |
| Wheat Starch | 404 g/Kg | 0.0% |
| Dextrinised Starch | 132 g/Kg | 0.0% |
| Dextrose Monohydrate | 0.0% | 686 g/Kg |
| % total digestible energy calculated from protein | 21.0% | 21.0% |
| % total digestible energy calculated from lipids | 16.0% | 16.0% |
| % total digestible energy calculated from carbohydrate | 56.8% | 62% |
| Digestible Energy | 16.1 MJ/ Kg | 16.9 MJ/ Kg |

All other ingredients including vitamins and mineral content were equivalent.

##### Virus infection, allergen exposure, and treatment of mice

Pneumonia virus of mouse (PVM) stock J3666 was prepared as previously described ^63^. A single dose of 10 plaque forming unit (PFU) of PVM in 10 µl or the same volume of diluent (10%, v/v, FCS in DMEM; Gibco) was administered (i.n. route) at PND7 or PND10 (as indicated in the study designs). The age of infection did not influence the effect of any given intervention. In two experiments, mice were inoculated (i.n. route) with RV1b (1x10^6^ TCID50) instead of PVM. RV1b was prepared, purified and titrated as previously described ^63^. In the model of experimental asthma, mice were inoculated with PVM or diluent at PND7, then exposed to low-dose cockroach allergen (CRE, Greer, USA; 1 µg, i.n. route) or diluent (PBS) at PND42, 49, 56, 63 ^64^. For the pDC adoptive transfer experiments, FACS-sorted pDC purified from the medLN of infected (6 dpi) HFD-reared donor pups were transferred to recipient LFD-reared WT pups (3,500 cells at 3 dpi, and 7,500 cells at 7 dpi).

For Treg cell adoptive transfer experiments, FACS-sorted Treg cells from medLN of infected (5 dpi) HFD-reared FoxP3^RFP+^ pups were transferred to recipient LFD-reared WT pups (25,000 cells at 5 dpi). For microbe (x10^8^ cfu) transplantation or propionate (1 g/kg) supplementation studies, the treatment was administered via oral gavage between PND5 to PND9 (see specific study designs), followed by inoculation with PVM at PND10. For pDC depletion, pDC-DTR transgenic mice and WT control mice, were injected (i.p. route) with DT (2 μg/kg, Sigma-Aldrich) at PND9, 11, 13, and 15. To inhibit ADAM10, mice were treated daily (oral gavage) with GI254023X (0.2 g/kg; Sigma-Aldrich) at PND5-9. To inhibit HDACs, mice were treated (oral gavage) with TSA (1 mg/kg; Abcam) at PND5, 7 and 9. To neutralise Flt3L, mice were injected (i.p. route) with goat anti-Flt3L Ab (600 μg/kg, R&D Systems) or an isotype-matched control at PND5, 7, and 9. To increase DC numbers, mice were injected (i.p. route) daily with exogenous Flt3L (1 mg/kg) at PND3-9. All treatments were performed under light isoflurane-induced anaesthesia. For antibiotic treatment, vancomycin (0.25 g/L), neomycin (0.5 g/L), metronidazole (0.5 g/L), ampicillin (0.5 g/L) and sweetener (8.0 g/L) were added to the drinking water from E13 until the end of the study.

##### Cross fostering (CF) study

Neonatal mice born to HFD-fed mothers were fostered by HFD- and LFD-fed mothers, and *vice-versa*. The fostered pups were removed from the biological mother and transferred to a different cage with the foster dam within 24 hours of birth.

##### Propionate Supplementation

Neonatal mice were treated (oral gavage) daily with a sodium salt of propionic acid (1 g/kg) at PND5-10 as previously described ^17, 18^. PBS was used as the vehicle control.

##### Milk collection

Milk was collected from lactating HFD- or LFD-fed mothers at post-partum day 7 (PPD7) and PPD10. Briefly, dams were separated from their litter for 2 hours then injected (i.p. route) with oxytocin (2 IU/kg). After induction of anaesthesia via injection (i.p. route) of a mixture of xylazine (10 mg/kg) and ketamine (100 mg/kg), the fur was removed and the teats and surrounding area cleaned with ethanol swabs to reduce contamination. Gentle pressure was applied to the nipple to release the milk, which was collected into sealed serum bottles containing 30% anaerobic glycerol in PBS (for later enrichment) or without glycerol (for later protein and 16S analysis), and stored at -80°C prior to further processing.

##### Preparation of enriched milk microbial transplantation (MMT)

Anaerobic YCFA broth was prepared as described ^65^. The broth was inoculated separately with milk collected from HFD- or LFD-fed mothers and incubated anaerobically at 37°C for 48 hours. The cultures were centrifuged at 7,500 rpm for 5 minutes to obtain bacterial pellets, then washed twice with anaerobic Ringer’s solution. The resulting bacterial pellets were resuspended with a sufficient volume of sterile 30% (v/v) glycerol in PBS to produce an optical density at 600 nm of 1.0, then stored at -80°C for future experiments.

##### Enriched Milk Microbial Transplantation (MMT)

Neonatal mice from HFD- and LFD-fed mothers were lightly anaesthetised, then treated (oral gavage) with the purified milk-enriched microbiota (10^8^ cfu in 50 µl) at PND5, 7, and 9. The following experimental groups were established:

1. (M)H H: MMT from HFD-fed mother’s milk to HFD-reared neonates

2. (M)H L: MMT from HFD-fed mother’s milk to LFD-reared neonates

3. (M)L H: MMT from LFD-fed mother’s milk to HFD-reared neonates

4. (M)L L: MMT from LFD-fed mother’s milk to LFD-reared neonates.

##### Isolation of axenic bacteria isolates from enriched milk microbiota

To isolate individual strains from the enriched milk microbiota of HFD-fed mothers, the enrichment cultures were plated on Brain Heart Infusion (BHI) agar medium and incubated overnight at 37°C in an anaerobic chamber. Discrete colonies with distinct morphologies were picked and anaerobically cultured with BHI broth. The cultures were centrifuged, washed twice with anaerobic Ringer’s solution, and the bacterial pellets resuspended with a sufficient volume of sterile 30% (v/v) glycerol in PBS to produce an optical density at 600 nm of 1.0, then were stored at -80°C for future experiments.

##### Preparation of anaerobic fecal suspensions and fecal microbiota transplant (FMT)

Fecal pellets were collected from culled HFD- and LFD-reared pups at PND7, and from HFD-fed mums at post-partum day 7 (PPD7), and placed separately into 0.5 mL micro centrifuge tubes on ice. The tubes were immediately transferred to an anaerobic chamber where the fecal pellets were transferred to 50 mL falcon tubes containing 5 large beads and 3 mL of anaerobic Ringer’s solution per tube. The 50 mL falcon tube was vortexed for 2 min before transferring the slurry into 1.5 mL Eppendorf tubes. After centrifugation (800xg for 2 min) to remove undigested particulates, the supernatant was carefully collected and transferred to 5 ml serum bottles closed with a butyl rubber stopper and containing sterile anaerobically prepared PBS with 30% (v/v) glycerol added in and stored at -80°C. Neonatal mice were lightly anaesthetised, then gavaged with the purified fecal microbiota (10^8^ cfu in 50 µl) at PND5, 7, and 9.

The following treatments and experimental groups were used in this experiment -
1. (F)H→H: FMT from HFD-reared pups to HFD-reared neonates
2. (F)H→L: FMT from HFD-reared pups to LFD-reared neonates
3. (F)L→H: FMT from LFD-reared pups to HFD-reared neonates
4. (F)L→L: FMT from LFD-reared pups to LFD-reared neonates.

##### Humanized mice

Breeding-age NSG-A2 mice were fed a HFD or LFD for 21 days prior to time mating, then time mated. After plugging the studs were removed, and the females maintained on the respective diet until the end of the experiment. Human cord blood units were obtained from the QLD Cord Blood Bank after written informed consent and approval from the Mater Human Research Ethics Committee. Cord blood mononuclear cells were enriched by Ficoll-Paque (GE Healthcare) density centrifugation and CD34+ hematopoietic stem cells isolated using a human CD34 isolation kit (Miltenyi Biotec). At PND 4, pups were irradiated (1Gy) prior to intrahepatic administration of 2 x10^5^ human CD34+ cells using a 30 g needle. The pups were subsequently euthanized at PND25.

##### Isolation of leukocytes from the gut lamina propria

To prepare a single cell suspension from the gut lamina propria, the small intestine and colon were excised, residual fat and lymphoid tissue trimmed off, and the lumen flushed with HBSS. After being cut into 0.25 cm^2^ pieces, the tissue was incubated in pre-digestion solution (5% FCS, 1 mM DTT and 5 mM EDTA in HBSS) at 37°C for 20 mins under continuous mechanical rotation. After a brief vortex, the sample was passed through a 70 µm cell strainer, and the process repeated using fresh pre-digestion solution, followed by a wash step in HBSS. As per the manufacturer’s guidelines (Lamina Propria Dissociation Kit, Miltenyi Biotec), the tissue samples were transferred to gentle MACS C Tubes and incubated in enzyme mix and digestion solution at 37°C for 30 mins under continuous mechanical rotation. Samples were then transferred to the gentle MACS dissociator (Miltenyi Biotec) and re-suspended in approximately 5 mL of PBS. After another pass through a 70 µm cell strainer, the cell suspension was washed in HBSS and centrifuged at 1600 rpm for 5 mins. To prepare a single cell suspension from the lungs, spleen, or LNs, the tissues were cut into 0.25 cm^2^ pieces then dissociated mechanically through a 70 µm cell strainer. To collect BM cells, the skin and flesh from the hind and upper limbs was removed and the cleaned bones pressed using a mortar and pestle (neonatal bones) or flushed (adult bone) with FACS buffer. To remove erythrocytes, all cell preparations were treated with Gey’s RBC lysis buffer. Single cell suspensions were washed in 2% FACS buffer and kept on ice until further analysis.

##### Flow cytometry

Non-specific binding was blocked by pre-incubation with anti-FcγRIII/II in FACS buffer (or combined rat and mouse serum for humanized mouse work), then specific antibodies conjugated with fluorochromes added to the cells for 30 minutes on ice in the dark. For intracellular staining, cells were further permeabilised for 20 minutes with the BD Fixation/Permeabilisation kit following the manufacturer’s instructions (BD Pharmingen). The cells were washed with BD fixation/permeabilising buffer then incubated with the relevant antibodies.

For detection of Flt3L, cells were incubated with goat anti-Flt3L, followed by a wash and incubation step with anti-goat AF555-conjugated antibody. Cells were then washed and analysed immediately on a BD LSR Fortessa cytometer or Cytek^®^ Aurora. pDC were identified as B220^+^, CD11c^+^, CD11b^−^, Siglec-H^+^. cDC1 and cDC2 were identified as MHCII^+^, CD11c^+^, CD11b^-^, CD64^−^, F4/80^−^, CD103^+^ and MHCII^+^, CD11c^+^, CD11b^+^, CD64^−^, F4/80^−^ respectively, as previously described ^66^. The effect of diet on lung cDC1 (MHCII^+^, CD11c^+^, CD26^+^, CD64^−^, CD172^−^, XCR1^+^) and cDC2 (MHCII^+^, CD11C^+^, CD26^+^, CD64^−^, CD172^+^, XCR1^−^, MAR1^−^) numbers was verified using a different immunophenotyping protocol ^39^. Progenitor populations were gated as follows: macrophage-DC progenitor (Lin^-^, CD11c^-^, MHCII^-^, CD135^+^, Ckit^+^, CD115^+^), common DC progenitors (Lin^-^, CD11c^-^, MHCII^-^, CD135^+^, Ckit^int^, CD115^+/-^), common lymphoid progenitors (Lin^-^, CD172α^+^, CD34^+^, Sca-1^lo^, CD135^+^, Ckit^lo^, Thy-1^-^) and pre-DCs (Lin^-^, MHCII^-^, CD11C^+^, CD135^+^, CD172α^lo^, SigH^+/-^, Ly6C^+/-^), as previously described ^67^. Neutrophils and eosinophils were identified as CD11b^+^Ly6G^+^ and CD3^-^, CD45R^-^, MHCII^-^, SiglecF^+^ respectively. ILC2s, Th2 and Tc2 cells were identified as (Lin^-^, CD90.2^+^, CD25^+^, ST2^+^), Th2 cells (CD3^+^, CD90.2^+^, CD4^+^, ST2^+^) and Tc2 cells (CD3^+^, CD90.2^+^, CD8^+^, ST2^+^), respectively. Treg cells were identified as CD3^+^, CD4^+^, CD25^+^, FoxP3^+^. Cell sorting of pDC (CD11c^+^, B220^+^, CD11b^-^, SiglecH^+^) and Treg cells (TCRb^+^, CD4^+^, CD25^+^, FoxP3^RFP+^) was performed using a BD FACSAria IIIu (Becton Dickinson).

##### Enteroid preparation and culture

Enteroids were prepared as previously described ^68^. Briefly, ileal tissue from individual LFD-reared neonatal mice was digested with collagenase type I (20 mg/ml; Invitrogen), and crypts plated and cultured in L-WRN conditioned media with a ROCK inhibitor (Y-27632, 10 μm; R&D Systems) and a TGFβIR inhibitor (SB 431542, 10 μm; R&D Systems) using 3-dimensional culture in Matrigel droplets (Corning). Enteroids were passaged 4 times for expansion before culture for 1 day in stem cell media and 2 days in differentiation media ^52^ with and without propionate (300μM). Enteroids were analysed for Flt3L expression by immunofluorescent staining of histogel (Thermo-scientific) blocked formalin-fixed paraffin-embedded sections.

##### Mediastinal lymph node recall assay

Excised mediastinal LN were gently pressed through a 70 μm cell strainer using a 1 ml syringe plunger to prepare a single cell suspension. Red blood cells (RBC) were lysed using Gey’s lysis buffer, and washed twice in RPMI-1640 medium containing 10% FBS, 2 mM L-gutamate, 1 mM sodium pyruvate, 20 mM HEPES, 100 U/ml penicillin-streptomycin mixture and 50 μM 2-mercaptoethanol. The cells (10^6^ cells/well) were transferred into a flat-bottom 96-well plate and stimulated with CRE extract (10 μg/ml) or media. After 72 hours, the culture supernatants were collected and stored at -80°C prior to cytokine analysis.

##### Quantification of cytokines and antibodies

The concentration of Flt3L, eotaxin-2/CCL24, CXCL1/KC, IFN-λ (IL-28A/B), EGF (all R&D systems), IL-6, IL-10, IL-1β, free active TGFβ1, IFN-γ (all Biolegend), sIgA (MyBioSource) was analysed by ELISA according to the manufacturer’s protocol. IL-4, IL-5, IL-13 were measured in medLN culture supernatant in asthma phase after PND66 using paired antibodies (BD Biosciences). Total IgE and CRE-specific IgG1 were quantified in the serum at PND66 by ELISA as described previously ^69^.

##### Decalcification of bones for histology

Cleaned bones were stored in 10% neutral buffered formalin for 24 hours before transferring to decalcification solution containing 15% EDTA (EDTA 150 g, sodium hydroxide 10 g, Na_2_HPO_4_ 10.22 g, NaH_2_PO_4_.H_2_O 3.86 g and distilled water q.s. to 1000 ml). The decalcification solution was replaced every 3 days. Decalcification of neonatal mouse thigh bones required between 7-14 days. Once the bones were at a soft, spongy consistency, the sample was washed in water for several hours, then dehydrated, and embedded in paraffin.

##### Immunohistochemistry and histology

Paraffin-embedded sections were sectioned and immunohistochemistry for PVM (1:8000 dilution; kindly provided by Dr Ulla Buchholz, Laboratory of Infectious Diseases, National Institute of Allergy and Infectious Diseases), α-smooth muscle actin (Sigma), and Muc5ac (ThermoFisher) was performed as described previously^70^. Prior to detection of Flt3L in the BM, the bones were decalcified, embedded and sectioned onto slides. To detect Flt3L, the sectioned BM, SI or colon was permeabilized in PBS containing 0.5% Triton for 10 minutes, incubated with a combination of 3% donkey serum and 7% normal goat serum in PBS for 30 minutes, then washed and incubated overnight at room temperature with goat anti-Flt3L (1:200; R&D Systems) in PBS with 10% FCS. To confirm specificity, anti-Flt3L was pre-incubated with Flt3L for 1 hr prior to tissue incubation. To visualize intestinal epithelial cells, the sections of gut tissue were also incubated overnight with anti-villin (Abcam). The next day, sections were washed in PBS/Tween 20 and incubated for 60 minutes with anti-goat-AF555 (Life Technologies) and anti-rabbit-AF647 (Life Technologies). After washing with PBS containing 0.5% triton, the sections were incubated for 5 minutes in DAPI solution (1:10,000) then mounted with fluorescent mounting media (DAKO). PVM positive AECs were counted and expressed as a percentage of total AECs. Sloughing of the airway epithelium was quantified by measuring the length of sloughed epithelium and expressing this as a percentage of perimeter length of the airway. The mucus score was calculated based on the percentage of Muc5ac positive cells and the percentage of obstructed airways as previously described ^23, 71^. Scanscope XT software was used to quantify ASM area around the small airways (a circumference less than 500 µm for neonatal mice and less than 800 µm for adult mice) and expressed as the square root of ASM area per micrometer (µm) of basement membrane as previously described ^23^. Five airways per mouse were enumerated for all pathologies. Flt3L was quantified as average integrated intensity of Flt3L expressed in the tissue of interest. Quantification was conducted with Aperio Image Scope (Aperio, Vista, Calif) and ImageJ software.

##### RNA Extraction and Quantitative reverse transcriptase polymeric chain reaction

RNA was extracted from lung, medLN, SI, colon, spleen and BM cells as previously described^72^. After DNAse treatment with Turbo DNAse (Ambion, Carlsbad, CA), RNA quantity and quality was measured using a Nanodrop spectrophotometer, then stored at −80°C. qRT-PCR was performed as previously described^22, 73, 74^. Primer sequences were as follows: Flt3L, forward 5′- GGCTTCTAGGGCTATGGGAC-3′ and reverse 5′-CTGCCAGAATTTCTCTCGGT-3′; Hprt, forward 5′-AGGCCAGACTTTGTTGGATTTGAA-3′ and reverse 5′- CAACTTGCGCTCATCTTAGGCTTT-3′. Gene expression was calculated using the –ΔΔCt method relative to the housekeeping gene Hprt, and expressed as fold change compared to the respective control.

##### Microbial DNA extraction

DNA was extracted from mouse feces and milk, as well as bacterial enrichments and isolates, with an adaptation to a previously described method^75^ using bead beating plus column purification with the Maxwell 16 MDx Instrument (Promega, USA). Here, 0.02 g to 0.1g of sample was transferred into a sterile 2 mL screw cap tube containing 0.4 g of sterile zirconia beads. After adding 600 µL of lysis buffer (50 mM EDTA, 50 mM Tris-HCl, pH 8.0, 500 mM NaCl, and 4% SDS), tubes were homogenized in a Precellys 24 Tissue Homogenizer at 4°C, 5000 rpm, and 3 x 60 s with 30 s rest. The lysates were incubated for 15 minutes at 70°C, with mixing by inversion once every 5 minutes. The lysate was then centrifuged for 5 min at 13.2k x *g* and 4°C, and the supernatant was transferred to a new 1.5 mL microcentrifuge tube and mixed with 30 µL Proteinase K (Maxwell 16 LEV Blood DNA Kit, Promega) by vortexing for 30 seconds. The mixture was then incubated at 56°C for 20 min, and the remaining extraction steps completed following the Maxwell 16 LEV Blood DNA cartridge protocol. After completion of extraction, the remaining paramagnetic particles were removed by centrifugation for 2 min at 10k x g at 4°C, followed by a 15 min incubation at 37°C in RNase A (10 mg/mL final concentration; QIAGEN). DNA concentration was measured using the Qubit dsDNA BR Assay Kit (ThermoFisher Scientific), and quality was checked visually by agarose gel electrophoresis.

##### Bacterial 16S rRNA gene amplicon sequencing and community profiling

Universal bacterial 16S rRNA genes (V6-V8) were PCR amplified using primers 926F (5’-AAA CTY AAA KGA ATT GRC GG-3’) and 1392wR (5’-ACG GGC GGT GWG TRC-3’)^76^. Both the reverse and forward primers were modified to include an Illumina overhang adapter at their 5’ ends to be compatible with Nextera XT indices i5 (Index 2) and i7 (Index 1), respectively. The subsequent PCR was performed using 1.0 µL bacterial DNA as template, in 1X PCR master mix containing buffer without Mg_2_+ (Invitrogen), Taq DNA Polymerase (0.625U) (Invitrogen), MgCl_2_ (300 µM) (Invitrogen), dNTPs (100 µM of each) (Invitrogen), and 250 µM of each primer. Then ultrapure grade water was added to make up to a total volume of 25 µL for each sample. SimpliAmp 96-well Thermocycler (Applied Biosystems) was used to perform amplifications. Amplification by PCR target DNA was tested using agarose gel electrophoresis to visually confirm the amplicon size and quality. Agencourt AMPure XP beads were used for the purification of all the PCR products, and subsequently index PCR was performed with the Nextera Index PCR XT kit (Illumina), following the manufacturer’s guidelines. Amplification by PCR was verified by agarose gel electrophoresis. After purification with AMPure XP beads, PicoGreen dsDNA Quantification Kit (Invitrogen) was used to quantify indexed amplicons. The indexed amplicons were pooled in equal quantities after normalizing their concentrations. Sequencing of the 16S rRNA gene-based libraries was done at the Queensland Brain Institute (QBI), University of Queensland, using a MiSeq Reagent Kit v3 (Illumina- 600 cycle and Illumina- 30% PhiX Control v3).

The sequencing data was processed via a modified UPASRSE approach^77^. Briefly: i) forward reads were quality filtered and clustered and an OTU table was produced using default parameters using USEARCH (v10.0.240)^78^; ii) taxonomy was assigned using BLASTN (v2.3.0+)^79^ from QIIME2 (v2017.9)^80^ and SILVA SSU (v128)^81^ and; iii) the OTU table was filtered for non-bacterial sequences using BIOM^82^. MAFFT (v7.221)^83^ was then used to produce representative OTU sequences for the 16S rRNA gene, and a masked OTU alignment was used to produce a phylogenetic tree using FastTree (v2.1.9)^84^ in QIIME2. Finally, the OTU table was rarefied to 1200 sequences for each sample, and the mean numbers of predicted (Chao1) and observed (Sobs) OTUs in addition to the Faith’s phylogenetic diversity index (Faith’s PD) were calculated using QIIME2. Heat trees were generated in MetaCoder (v0.3.4)^85^ after filtering the OTU table for only OTUs with ≥0.5% relative abundance in at least one sample.

##### Bacterial isolate shotgun genomic sequencing and metabolic reconstruction.

Isolate bacterial whole genomic DNA (gDNA) was sequenced to enable genome recovery, species identification, and metabolic reconstruction. Libraries from gDNA were prepared using Nextera DNA Flex kit (Illumina). gDNA concentration was normalized for each isolate library, and all isolates were pooled for sequencing on an Illumina NextSeq 550 at the Australian Centre for Ecogenomics (ACE), University of Queensland, generating 0.5 Gbp per isolate. Trimmomatic (v0.36)^86^ was used to trim reads to ≥ Q10 trailing bases and an average of ≥ Q15 across a 4 base window. Trimmed reads of < 75 bases were then discarded. These data were individually assembled using SPAdes (v3.11.0)^87^ with default parameters, and assembly statistics were evaluated using CheckM (v1.0.7)^88^. To determine the taxonomy of the 10 isolates, the resulting genomes were placed in a phylogenetic tree in GTDB-Tk (v0.3.2)^89^ using the *de novo* workflow. The multisequence alignments of the isolate genomes, as well as GTDB representatives of species in the same genus as the isolates, were selected in ARB (v6.0.6)^90^ and exported for inferring final trees using IQ-TREE (v2.0.5)^91–93^. Trees were initially inferred using the Extended model selection function, and then the model generating the best fit was selected to infer the final tree, with bootstrap values generated using 1000 iterations. The final trees were decorated with GTDB-Tk taxonomy, and visualised using ARB. Metabolic reconstruction was completed by uploading genomes to MicroScope (v3.14.0)^94^ for gene identification and functional annotation.

##### Microbiome analysis

Isolate genomes were associated to gut microbiome OTUs by mapping the shotgun genomic reads to OTU reference sequences using the sensitive local alignments preset options in bowtie2 (v2.3.4.1)^95^. OTU coverage profiles were then generated in SAMtools (v1.9)^96^ using the depth function at all positions. Those OTUs with the greatest average coverage across the full 250bp length were selected as the associated OTU for each isolate genome. In all cases, the highest-level taxonomic assignment of the OTU matched the taxonomic classification of the isolate genome. Redundancy analysis (RDA) was performed with the rda function in the R vegan package, and ordination plots were generated with the plots function using ordiellipse to draw the standard deviation elipses around groups. The compositional similarity of raw and enriched milk bacterial communities, as represented by Hellinger distances, were compared using a mantel test as implemented in the R package *vegan*. Heat trees were generated with the heat_tree function in the R Metacoder package^85^. Bar graphs were generated with the bargraph.CI function in the R sciplot package, and ad hoc labels were assigned based on Tukey’s range test results (p<0.05) using the Tukey HSD function in the R stats package.

##### SCFA analysis

Quantification of SCFA in milk was performed by LC-MS/MS following derivatization with O-benzylhydroxylamine (O-BHA) and N-(3-dimethylaminopropyl)-N-ethylcarbodimide (EDC) ^97^. 10µl of milk was vortexed with 10µl of 1µg/ml of internal standard 2- ethyl butyric acid in milliQ water followed by deproteinisation with 60µl methanol. Following centrifugation, supernatant was incubated with 10µl of 0.1M O-BHA in methanol and 10µl of 0.25M of EDC in methanol at 25°C for 1 hour. Samples were then diluted with 100µl of milliQ water and extracted with 400µl of dichloromethane through 10 min of vigorous shaking followed by centrifugation. The organic layer was then extracted, evaporated, reconstituted in 50% methanol, and transferred into vials for LC-MS/MS analysis (Agilent 1200 series HPLC system; AB SCIEX API 3200). The mass spectrometer was operated in multiple reaction monitoring scan mode for identification of fragments in positive ionization mode at unit resolution. For quantitative purposes, one mass/charge (m/z) transition per analyte was monitored, and concentrations of derivatized analytes determined by the peak area ratio of analyte to internal standard using Multiquant software (AB SCIEX). SCFAs were measured in serum and fecal pellets by gas chromatography. For fecal samples, 0.02 - 0.1 g of feces was mixed with 350 µL of ultrapure water in a 2.0 mL Eppendorf tube. The mixture was homogenized to a slurry by vortexing for 5 to 10 minutes. For serum samples, 200 µL of clear serum was mixed with ultrapure water to a final volume of 350 µL. Solids were removed from all samples with a 0.22 µm filter, and 90 µL of filtrate was then mixed with 10 µL of internal standard solution containing 10% (v/v) formic acid and 1 g/L 2-ethylbutyric acid. The mixture was then transferred into a glass vial containing a small glass insert and SCFA concentrations measured by GC-FID (PerkinElmer, Australia) with polar capillary column (DB-FFAP; Agilent Technologies, USA) at 140°C.

##### Statistical analysis

For analyses of data, the Mann–Whitney *U* test was used for two-group comparisons based on the assumption that samples followed a non-normal distribution. For comparisons of > than two groups, statistical analysis was performed with one- or two-way ANOVA followed by Tukey’s post-hoc test for multiple comparisons (unless stated otherwise). For analysis of spectral flow cytometry, the Mann-Whitney *U* test was used to compare the Log_2_ fold-change (L2FC) in marker expression of HFD replicates from the LFD mean with the L2FC of LFD replicates from the LFD mean. For all quantifications, n represents the number of mice. Details of the statistical tests applied are reported in the figure legends. *p* values less than 0.05 were considered significant (* p < 0.05; ** p < 0.01; *** p < 0.001).

##### Supplemental information titles and legends

**Figure S1. A maternal LFD increases the severity of viral LRI in the offspring.**

(A) Experimental design.

(B) Weight gain in mother from three weeks prior to time mating, during gestation and post-partum.

(C) Weight gain in naïve pups (age-matched with PVM infected pups).

(D) IL-6, KC (CXCL1) and IL-1β production in lung.

(E) Representative image of muc5ac immunoreactivity (red) in lung tissue of HFD- and LFD-reared pups at 10 dpi, and quantification of muc5ac score.

(F) Lung ILC2s at 10 dpi.

(G) Eosinophils in BALF at 10 dpi.

(H) Eotaxin-2 concentration in lung homogenate at 10 dpi.

(I) IL-13 production in lung.

(J) ASM area in lung at 10 dpi.

(K) Experimental design for RV infection model.

(L) Weight gain in RV1b-infected pups.

(M) Lung neutrophils at 10 dpi.

(N) Epithelial sloughing at 10 dpi.

(O) Quantification of muc5ac (score) at 10 dpi.

(P) IL-6 production in lung homogenates at 10 dpi.

(Q) CXCL1 production in lung homogenates at 10 dpi.

Black boxes or circles represents HFD-reared pups, red boxes or circles represent LFD-reared pups. Data are represented as mean ± SEM (B-E, H, I and L) or box-and-whisker plots (median, quartiles, and range) (F, J, and M-Q), n=5-10 mice per group. *P* values (* p<0.05, ** p<0.01, *** p<0.001) were derived by two-way ANOVA followed by Sidak’s (B-E, H, I and L) or Tukey’s (F, G, J and M-Q) multiple comparisons tests.

**Figure S2: LFD-reared pups that develop PVM- or RV-induced sLRI in infancy are predisposed to experimental asthma in later life.**

(A) Lung ILC2s in PVM/CRE-exposed mice at PND66 (A-D; study design in Fig. S1A).

(B) CRE-specific IgG1 production in serum at PND66.

(C) Lung neutrophils at PND66.

(D) IL-4 (left panel), IL-13 (middle panel) and IFN-γ (right panel) production in MedLN recall assay.

(E-G) Lung eosinophils (E), neutrophils (F), and ILC2s (G) in RV1b/CRE-exposed mice at PND66 (E-I, study design in Fig. S1A).

(H) IgE concentration in serum at PND66.

(I) Quantification of muc5ac (score) at PND66.

Black boxes represents HFD-reared pups, red boxes represent LFD-reared pups. Data are represented as box-and-whisker plots (median, quartiles, and range), n=4-9 mice per group. *p* values (* p<0.05, ** p<0.01, *** p<0.001) were derived by two-way ANOVA followed by Tukey’s multiple comparisons test.

**Figure S3: Treg cell and pDC gating strategy.**

(A) Representative FACS plots showing the gating strategy for FoxP3+ Tregs in PVM infected lung.

(B) Percent KI67^+^ Treg cells at PND7.

(C) Representative FACS plots of Nrp1 and Helios expression in Treg cells in lungs of HFD- and LFD-reared pups at 5 dpi.

(D) Mean fluorescence intensity (MFI) of CD39 expression on FoxP3+ Treg cells (left panel) and IL-10 production in BALF (right panel).

(E) Representative FACS plots showing the gating strategy for pDC in the lung.

(F) Percent KI67^+^ pDCs.

Black boxes represents HFD-reared pups, red boxes represent LFD-reared pups. Data are represented as box-and-whisker plots (median, quartiles, and range), n=4-7 mice per group. *P* values (* p<0.05, ** p<0.01, *** p<0.001) were derived by Mann Whitney U test (B and D left panel), comparing HFD-reared with LFD-reared mice and by two-way ANOVA followed by Tukey’s multiple comparisons test (D left panel, and F).

**Figure S4: cDC gating strategy.**

Representative FACS plots showing the gating strategy for cDC1 and cDC2 in the lung.

**Figure S5: A maternal LFD affects cDC development and expansion during acute LRI in the offspring.**

(A and B) cDC1 (left panel) and cDC2 (right panel) numbers in the MedLN (A) and lung (B).

(C) cDC1 (left panels) and cDC2 (right panels) numbers at 5 dpi in the MedLN and lung, using a modified gating strategy,

(D) cDC1 (left panels) and cDC2 (right panels) numbers in BM at PND7.

(E) Gating strategy for MDP (macrophage-DC progenitor), CDP (common DC progenitor), CLP (common lymphoid progenitors) and pre-DCs.

(F) MDP (macrophage-DC progenitor), CDP (common DC progenitor), CLP (common lymphoid progenitors) and pre-DCs in BM at PND7.

Black boxes or circles represent HFD-reared pups, red boxes or circles represent LFD-reared pups. Data are represented as mean ± SEM (A and B) or box-and-whisker plots (median, quartiles, and range) (C, D, F), n=5-8 mice per group. *P* values (* p<0.05, ** p<0.01, *** p<0.001) were derived by Mann Whitney U test (C, D, F), comparing HFD-reared with LFD-reared mice and by two-way ANOVA followed by Sidak’s multiple comparisons test (A and B).

**Figure S6: Adoptive transfer of pDCs or Treg cells to LFD-reared pups confers protection against sLRI.**

(A) Study design of pDC adoptive transfer.

(B) Treg cell numbers in lung at 5 dpi.

(C) IFN-λ protein in lung homogenate at 5 dpi.

(D) Quantification of muc5ac (left panel) and ASM area (right panel) at 10 dpi.

(E) Study design of Treg cell adoptive transfer.

(F) Weight gain post PVM infection (upper left panel), lung neutrophils (upper right panel), epithelial sloughing (lower left panel) and ASM area (lower right panel) at 10 dpi.

(G) Representative FACS plots showing the gating strategy for human pDCs in BM.

(H) Human CD45+ cell numbers in BM and lung.

Black boxes or circles represent HFD-reared pups, red boxes or circles represent LFD-reared pups. Data are represented as mean ± SEM (F) or box-and-whisker plots (median, quartiles, and range) (B-D, F and H), n=5-13 mice per group. *P* values (* p<0.05, ** p<0.01, *** p<0.001) were derived by one-way (B-D, and F) and by two-way ANOVA (F upper left panel) followed by Tukey’s multiple comparisons test.

**Figure S7: A LFD alters the maternal microbiome and SCFA production and cross-fostering from an LFD- to a HFD-fed mother protects against sLRI.**

(A) RDA ordination highlighting diet associated differences in the composition of fecal microbiomes mothers at first trimester (E7) (p=0.001, PERMANOVA) (left panel) and third trimester (E18.5) (p=0.001) (right panel). Circles are individual samples and ellipses represent the standard deviation around the centroid of each group.

(B) Propionic acid (left panel), acetic acid (middle panel) and butyric acid (right panel) concentration in feces and serum at E18.5.

(C and D) Acetic acid (C) and butyric acid (D) concentration in feces and serum of neonatal mice.

(E) RDA ordination highlighting differences in the composition of fecal microbiomes after cross-fostering (p=0.001, PERMANOVA). Circles are individual samples and ellipses represent the standard deviation around the centroid of each group.

(F and G) Propionic acid, acetic acid and butyric acid concentration in serum (F) and feces (G) after cross-fostering.

(H) PVM immunoreactivity as a percentage of AECs (left panel) and IFN-λ production (right panel) in lung homogenate at 7 dpi.

(I) pDCs (left panel) and Treg cells (right panel) in MedLN at 5 dpi.

Black boxes or circles represent HFD-reared mothers or pups, red boxes or circles represent LFD- reared mothers or pups. Data are represented as mean ± SEM (C, D) or box-and-whisker plots (median, quartiles, and range) (B, F-I), n=4-11 mice per group. *P* values (* p<0.05, ** p<0.01, *** p<0.001) were derived by Mann Whitney U test (B), comparing HFD-reared with LFD-reared mice or by one-way (F-H) or two-way ANOVA (C) followed by Tukey’s multiple comparisons test.

**Figure S8: A FMT from HFD-reared neonates but not a HFD-fed dam and MMT from HFD-fed mothers protects against sLRI in susceptible LFD-reared pups.**

(A) Experimental design of FMT study. Neonatal mice were transplanted with a fecal slurry obtained from HFD-fed dams (B-E) or HFD- or LFD-reared (groups described on right hand side) neonates (F-I) at PND5, 7 and 9, then inoculated with PVM at PND10.

(B) PVM immunoreactivity as a percentage of AECs at 7 dpi.

(C) IFN-λ production in lung homogenate at 7 dpi.

(D) Lung neutrophils at 10 dpi.

(E) Epithelial sloughing at 10 dpi.

(F) PVM immunoreactivity as a percentage of AECs at 7 dpi.

(G) IFN-λ production in lung homogenate at 7 dpi.

(H) Lung neutrophils at 10 dpi.

(I) Epithelial sloughing at 10 dpi.

(J) Experimental design of MMT study. A MMT was administered to HFD- and LFD-reared neonates (groups described on right hand side) at PND5, 7 and 9, followed by PVM inoculation at PND10.

(K) Epithelial sloughing at 10 dpi.

(L) PVM immunoreactivity as a percentage of AECs (left panel) and IFN-λ production (right panel) in lung homogenate at 7 dpi.

(M) Propionic acid concentration in feces (left panel) and serum (right panel).

Black boxes represent HFD-reared pups, red boxes represent LFD-reared pups. Data are represented as box-and-whisker plots (median, quartiles, and range) (B-I, and K -M), n=5-7 mice per group. *P* values (* p<0.05, ** p<0.01, *** p<0.001) were derived by one-way ANOVA (B-I, and K -M) followed by Tukey’s multiple comparisons test.

**Figure S9. Taxonomic classification of milk-derived bacterial isolates and their multiple metabolic pathways for propionate production.**

(A) Phylogenetic trees for the 8 recovered milk bacterial isolate genomes. Genomes are split between the families *Enterococcaceae* (top) and *Enterobacteriaceae* (bottom). Branches containing the isolate genomes are expanded to show taxonomic classification, shown as brackets to the right. Additional reference branches are collapse into grey clades, with the clade size listed in square brackets after the taxonomic label, and clade max and min branch lengths shown as the top and bottom edges of the clade, respectively. Branch bootstrap support is indicated by circles at the branch points, where open circles indicate ≥ 50%, grey circles indicate ≥ 75%, and black circles indicate ≥ 90% support. Phylogenetic distance is indicated by the scale at the bottom of each tree.

(B) Propionate-producing metabolic pathways originating from central carbon metabolism and amino acid degradation. Central carbon metabolism pathways are listed to the left, with the pathway products that link to propionate pathway intermediates listed below. Pathways to the right are then simplified to key substrates and branch points. Square labels at each branch indicate presence or absence of the given function, where *E. steigerwaltii* functions are identified by the green upper square, and *E. avium* functions by the brown lower square. Open squares indicate absence of the given function for the given lineage.

(C) Propionate production by *E. steigerwaltii* and *E. avium* lineages cultured in vitro in YCFA media.

**Figure S10. Transplantation of milk-derived isolates confers protection against sLRI to LFD- reared pups.**

Bacterial isolates (*E. steigerwaltii,* or *E. avium*) were transplanted to LFD-reared pups at PND5, 7 and 9, and prior to PVM inoculation at PND10.

(A) Weight gain in PVM-infected neonates.

(B) IFN-λ production in lung homogenate (left panel) and PVM immunoreactivity as a percentage of AECs (right panel) at 5 dpi.

(C) Epithelial sloughing at 10 dpi.

(D) Butyrate and acetate in feces.

(E) Butyrate, acetate and propionate in serum.

(F) Hellinger-transformed gut microbiome relative abundances of isolate-associated OTUs in HFD, LFD, and isolate-inoculated pups for *E. steigerwaltii* (left panel) and *E. avium* (right panel).

Error bars represent standard error for each set of samples, and *P* values (* p<0.05, ** p<0.01, *** p<0.001) relative to LFD (A-E)and HFD (F) were derived by one way ANOVA followed by Tukey’s HSD comparing groups as shown.

(G) RDA ordination highlighting differences in the composition of fecal microbiomes of HFD, LFD, and isolate-transplanted pups (all isolates in the left panel (p=0.001), *E. steigerwaltii* in the middle panel (p=0.001), and *E. avium* in the right panel (p=0.001)). Circles are individual samples and ellipses represent the standard deviation around the centroid of each group. Grey ‘x’s are individual OTUs, and black ‘x’s are OTUs which fall ≥2.5 standard deviations from the origin. Isolate-associated OTUs are marked by red arrows.

Data are represented as mean ± SEM (A) or box-and-whisker plots (median, quartiles, and range) (B-E), n=4-11 mice per group.

**Figure S11. Propionate supplementation confers protection against sLRI to LFD-reared pups and MMT, *E. avium* transplantation or propionate supplementation increases SCFAs in GF mice**

(A) Experimental design of propionate supplementation study. B-H: Data from neonatal LRI phase.

I-L: Data from later-life asthma phase.

(B) Propionic acid in serum at 5 dpi.

(C) Epithelial sloughing at 10 dpi.

(D) PVM immunoreactivity as a percentage of AECs at 5 dpi.

(E) IFN-λ concentration in lung homogenate at 5 dpi.

(F) pDCs (F) and Treg cells (G) in lung at 5 dpi.

(H) cDC1 and cDC2 in lung and MedLN at 5 dpi.

(I) Lung neutrophils at PND66.

(J) Muc5ac score at PND66.

(K) IL-13 levels in MedLN culture supernatant.

(L) CRE-specific IgG1 in serum at PND66.

(M) SCFA levels in feces (top panels) and serum (lower panels) in naïve GF mice at PND10.

Black boxes or circles represent HFD-reared pups, red boxes or circles represent LFD-reared pups. Data are represented as mean ± SEM (L) or box-and-whisker plots (median, quartiles, and range) (B-J,M), n=5-12 mice per group. *P* values (* p<0.05, ** p<0.01, *** p<0.001) were derived by one-way (M) and two-way ANOVA (B-L) followed by Tukey’s multiple comparisons test.

**Figure S12. Propionate-induced protection of LFD-reared pups against sLRI is ablated upon pDC depletion.**

(A) Experimental design. pDC-DTR transgenic and non-transgenic (WT) littermate controls reared to LFD-fed mothers were treated daily with sodium propionate (1 g/kg) or vehicle (oral gavage) from PND5 to 10. All mice received low dose diphtheria toxin (DT, 15 ng, i.p. route) at PND9, 11, 13 and 15, and were inoculated with PVM at PND10.

(B) Lung neutrophils (left panel) and epithelial sloughing (right panel) at 10 dpi.

(C) PVM immunoreactivity as a percentage of AECs (left panel) and IFN-λ production in lung homogenate at 5 dpi (right panel).

(D) pDC (left panel) and Treg cell (right panel) numbers in MedLN at 5 dpi.

Data are represented as box-and-whisker plots (median, quartiles, and range) (B-D), n=5-14 mice per group. *P* values (* p<0.05, ** p<0.01, *** p<0.001) were derived by one-way ANOVA followed by Tukey’s multiple comparisons test.

**Figure S13. Maternal diet affects Flt3L expression in the BM and gut of the offspring.**

(A) Immunoflourescence following pre-incubation of anti-Flt3L with exogenous Flt3L.

(B) Representative micrographs of Flt3L expression in the small intestine of HFD- and LFD-reared pups at PND10 and 42 (left panel: Flt3L and villin; middle panel: Flt3L; right panel: villin).

(C) Flt3L expression in BM cells measured by flow cytometry (MFI, mean fluorescent intensity).

(D) Experimental design of Adam10 inhibitor study.

(E and F) Flt3L expression by IECs (E) and BM cells (F) stained by flow cytometry, with and without cellular permeabilisation.

(G) pDCs in BM at PND10.

(H) Treg cells in MedLN at 5 dpi.

(I) Epithelial sloughing at 10 dpi.

(J-L) Serum Flt3L production after cross-fostering (J), MMT (K), or isolate transplantation (L).

(M) Effect of maternal antibiotic (ABX) exposure on epithelial sloughing, lung neutrophils, and Treg cells in MedLN in the offspring,

(N) Experimental design of pan-HDAC inhibitor study.

Black boxes represent HFD-reared pups, red boxes represent LFD-reared pups, grey shaded boxes represent propionate supplementation. Data are represented as box-and-whisker plots (median, quartiles, and range) (C, E-M), n=4-17 mice per group. *P* values (* p<0.05, ** p<0.01, *** p<0.001) were derived by Mann Whitney U test, comparing vehicle and antibiotics treated groups

(M) and by one-way (E-L) and two-way ANOVA (C) followed by Tukey’s multiple comparisons test.

**Figure S14. Anti-Flt3L ablates propionate-induced protection against sLRI whereas exogenous Flt3L confers protection to LFD-reared pups against sLRI.**

(A) Experimental design of Flt3L neutralisation study.

(B) pDC in lung (left panel) and colon (right panel) at PND10. (C and D) cDC1 and cDC2 in BM, SI and MedLN at PND10.

(E) Treg cells in lung (left panel) and colon (right panel) at PND10. (F and G) pDCs (F) and Treg cells (G) in lung at 5 dpi.

(H) Study design, HFD- and LFD-reared pups were treated with exogenous Flt3L (600 µg/kg, i.p.) daily from PND3 to 9, and euthanised at PND10 or inoculated with PVM or diluent at PND10 and killed at 10 dpi.

(I) pDCs in BM, lung, and MedLN at PND10.

(J) Weight gain in PVM infected neonates.

(K) Lung neutrophils (left panel) and epithelial sloughing (right panel) at 10 dpi.

Black boxes represent HFD-reared pups, red boxes represent LFD-reared pups. Data are represented as mean ± SEM (J) or box-and-whisker plots (median, quartiles, and range) (B-G, I and

K), n=5-6 mice per group. *P* values (* p<0.05, ** p<0.01, *** p<0.001) were derived by one-way (BG) and by two-way ANOVA (I-K) followed by Tukey’s multiple comparisons test.

