## Supplemental Figures for "The maternal microbiome regulates infant respiratory disease susceptibility via intestinal Flt3L expression and plasmacytoid dendritic cell hematopoiesis"

#### Figure S1

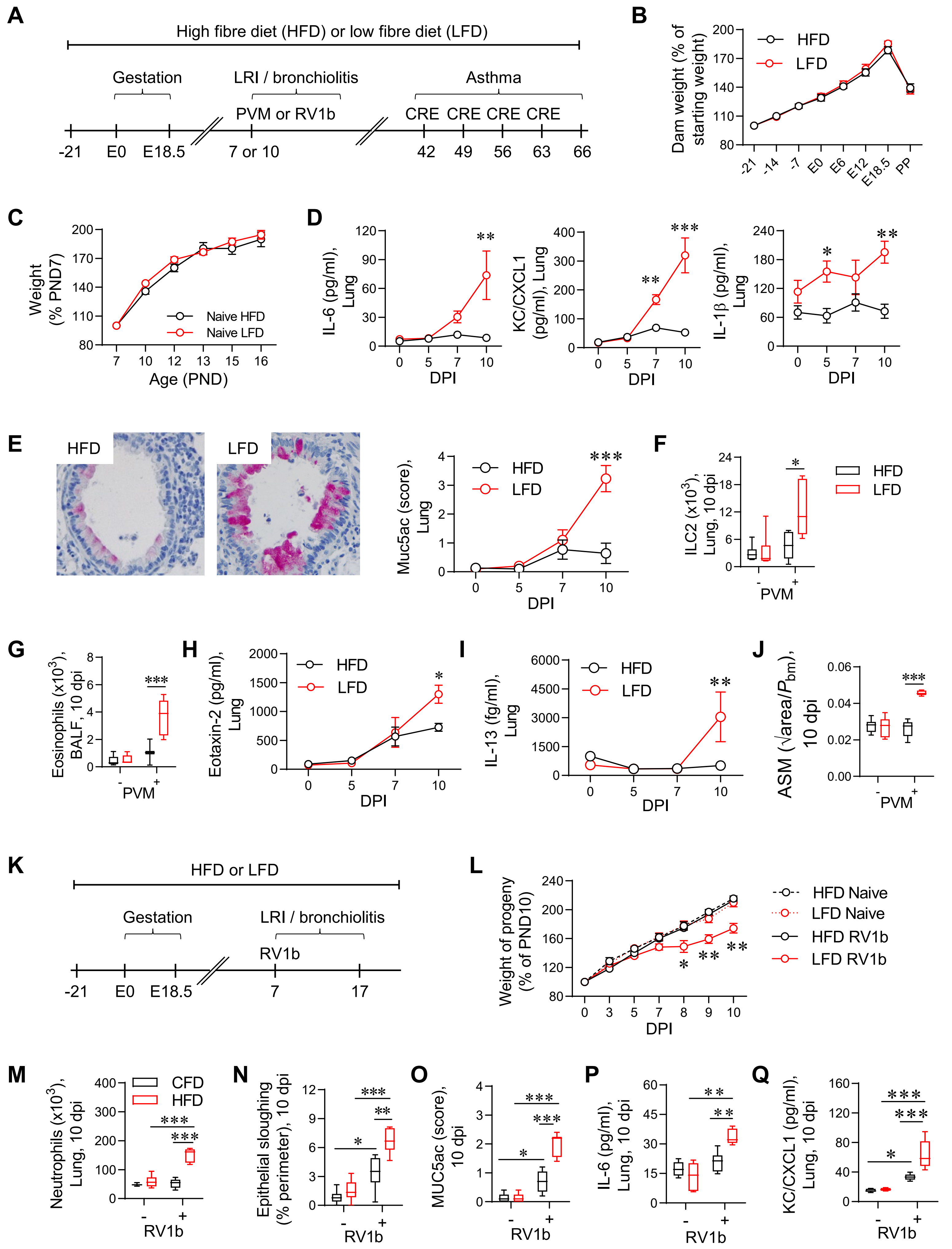
A
High fibre diet (HFD) or low fibre diet (LFD)
Gestation
LRI / bronchiolitis
Asthma
PVM or RV1b
CRE CRE CRE CRE
-21 E0 E18.5 7 or 10 42 49 56 63 66
B
Dam weight (% of starting weight)
HFD
LFD
C
Weight (% PND7)
Naive HFD
Naive LFD
Age (PND)
D
IL-6 (pg/ml), Lung
KC/CXCL1 (pg/ml), Lung
IL-1β (pg/ml), Lung
DPI
E
HFD
LFD
Muc5ac (score), Lung
F
ILC2 (x10³), Lung, 10 dpi
PVM
G
Eosinophils (x10³), BALF, 10 dpi
H
Eotaxin-2 (pg/ml), Lung
I
IL-13 (fg/ml), Lung
J
ASM (√area/Pbm), 10 dpi
K
HFD or LFD
Gestation
LRI / bronchiolitis
RV1b
-21 E0 E18.5 7 17
L
Weight of progeny (% of PND10)
HFD Naive
LFD Naive
HFD RV1b
LFD RV1b
M
Neutrophils (x10³), Lung, 10 dpi
CFD
HFD
RV1b
N
Epithelial sloughing (% perimeter), 10 dpi
O
MUC5ac (score), 10 dpi
P
IL-6 (pg/ml), Lung, 10 dpi
Q
KC/CXCL1 (pg/ml), Lung, 10 dpi

### Figure S2

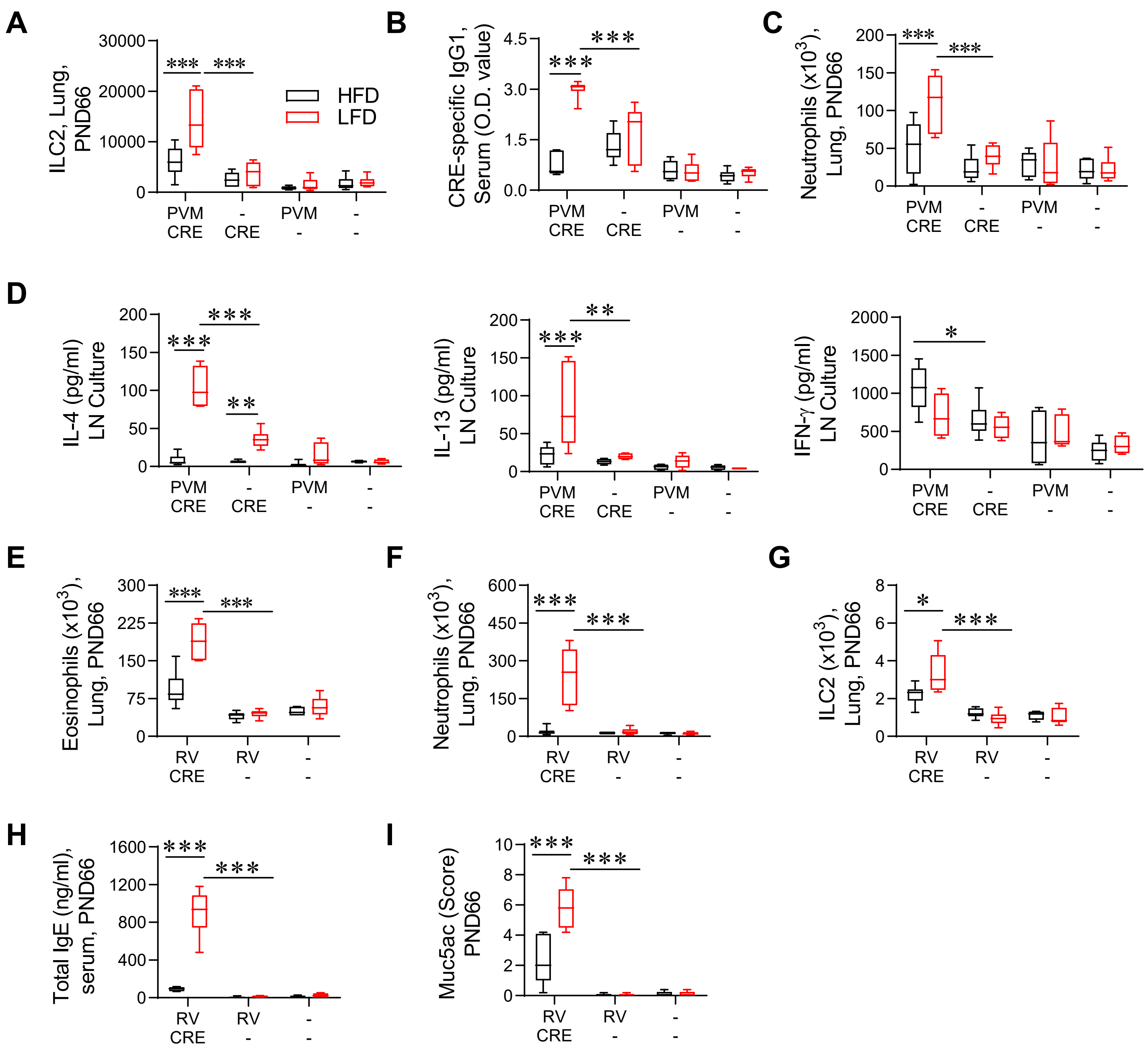
A
ILC2, Lung, PND66
HFD
LFD
PVM CRE
- CRE
PVM -
- -
B
CRE-specific IgG1, Serum (O.D. value)
C
Neutrophils (x10^3), Lung, PND66
D
IL-4 (pg/ml) LN Culture
IL-13 (pg/ml) LN Culture
IFN-γ (pg/ml) LN Culture
E
Eosinophils (x10^3), Lung, PND66
RV CRE
RV -
- -
F
Neutrophils (x10^3), Lung, PND66
G
ILC2 (x10^3), Lung, PND66
H
Total IgE (ng/ml), serum, PND66
I
Muc5ac (Score) PND66

### Figure S3

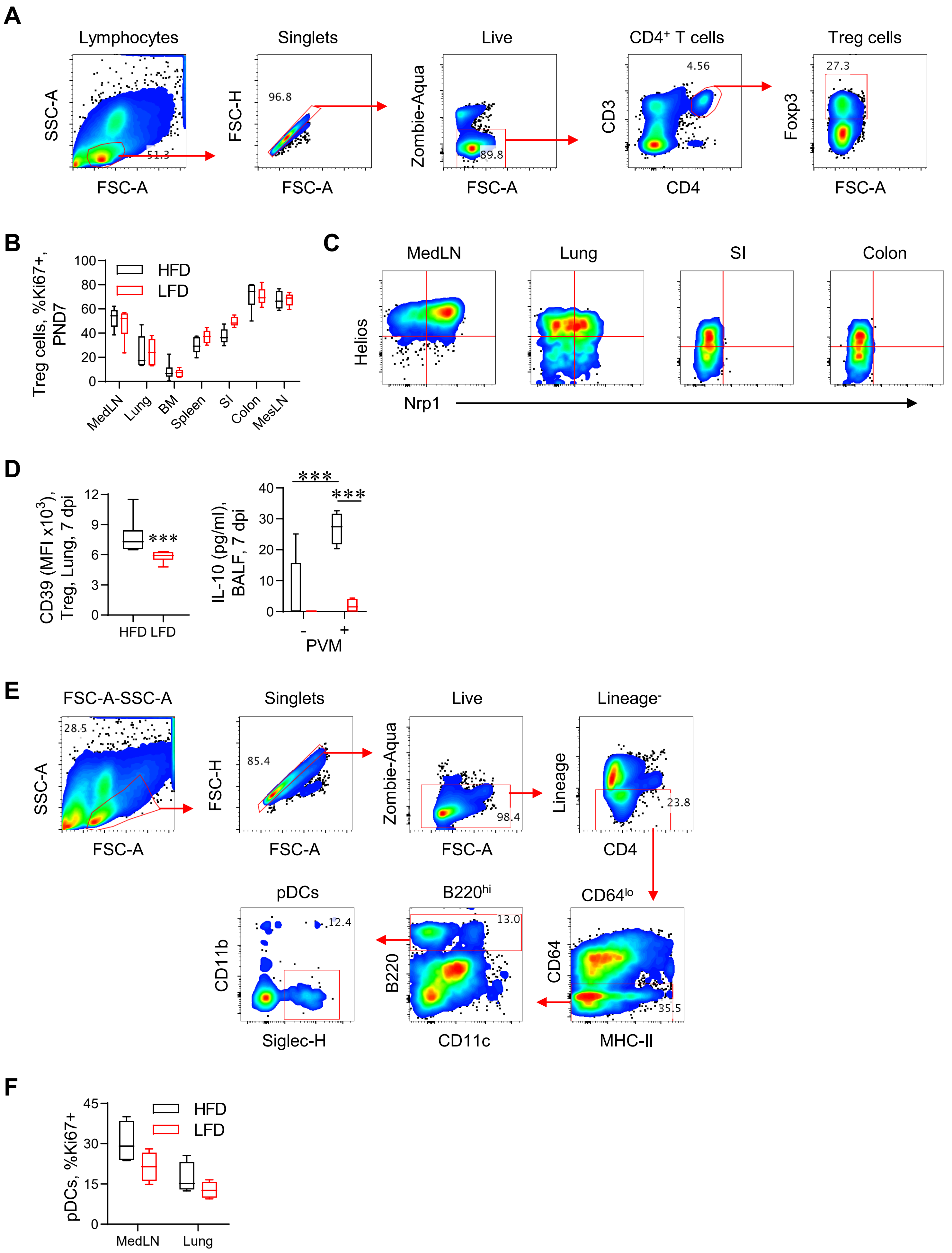
A
Lymphocytes
Singlets
Live
CD4⁺ T cells
Treg cells
SSC-A
FSC-A
51.3
FSC-H
96.8
Zombie-Aqua
89.8
CD3
CD4
4.56
Foxp3
27.3
B
Treg cells, %Ki67+, PND7
HFD
LFD
MedLN
Lung
BM
Spleen
SI
Colon
MesLN
C
MedLN
Lung
SI
Colon
Helios
Nrp1
D
CD39 (MFI x10³), Treg, Lung, 7 dpi
HFD LFD
***
IL-10 (pg/ml), BALF, 7 dpi
PVM
-
+
E
FSC-A-SSC-A
Singlets
Live
Lineage⁻
SSC-A
FSC-A
28.5
FSC-H
85.4
Zombie-Aqua
98.4
Lineage
CD4
23.8
pDCs
B220hi
CD64lo
CD11b
Siglec-H
12.4
B220
CD11c
13.0
CD64
MHC-II
35.5
F
pDCs, %Ki67+
HFD
LFD
MedLN
Lung

**Figure S4**

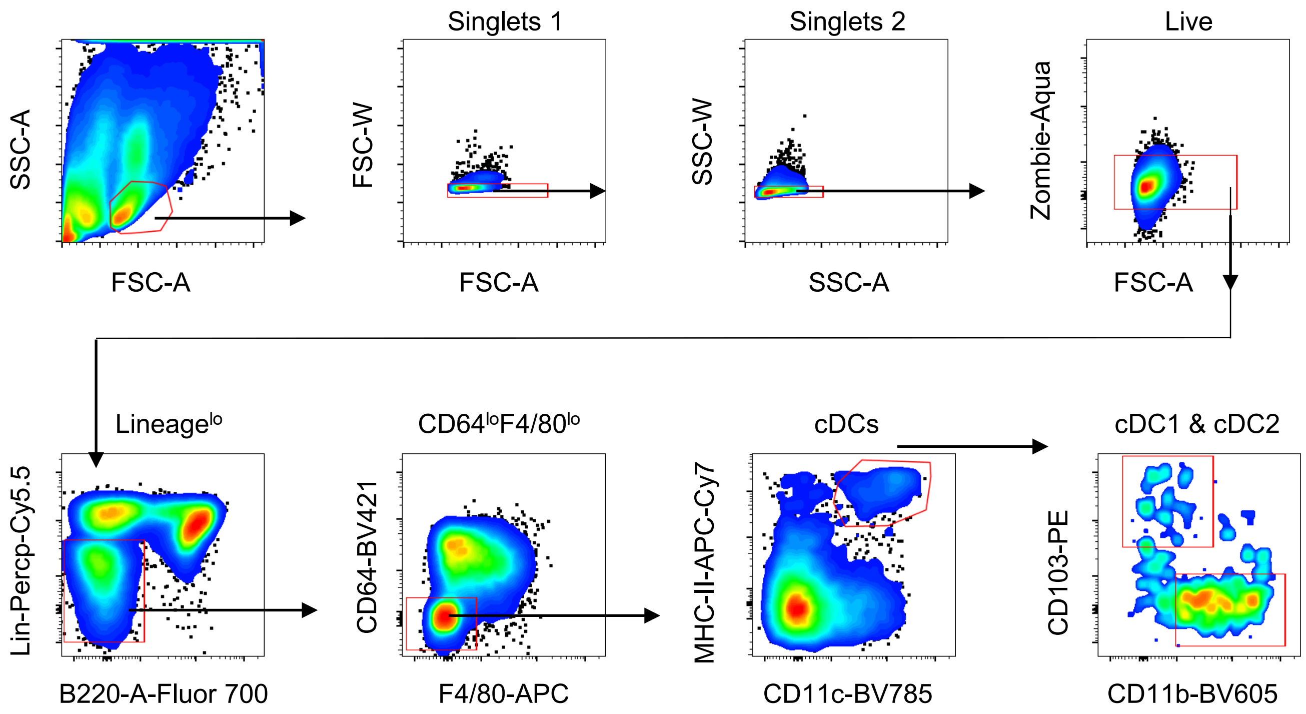

SSC-A
FSC-A
Singlets 1
FSC-W
FSC-A
Singlets 2
SSC-W
SSC-A
Live
Zombie-Aqua
FSC-A
Lineage^lo
Lin-Percp-Cy5.5
B220-A-Fluor 700
CD64^loF4/80^lo
CD64-BV421
F4/80-APC
cDCs
MHC-II-APC-Cy7
CD11c-BV785
cDC1 & cDC2
CD103-PE
CD11b-BV605

### Figure S5

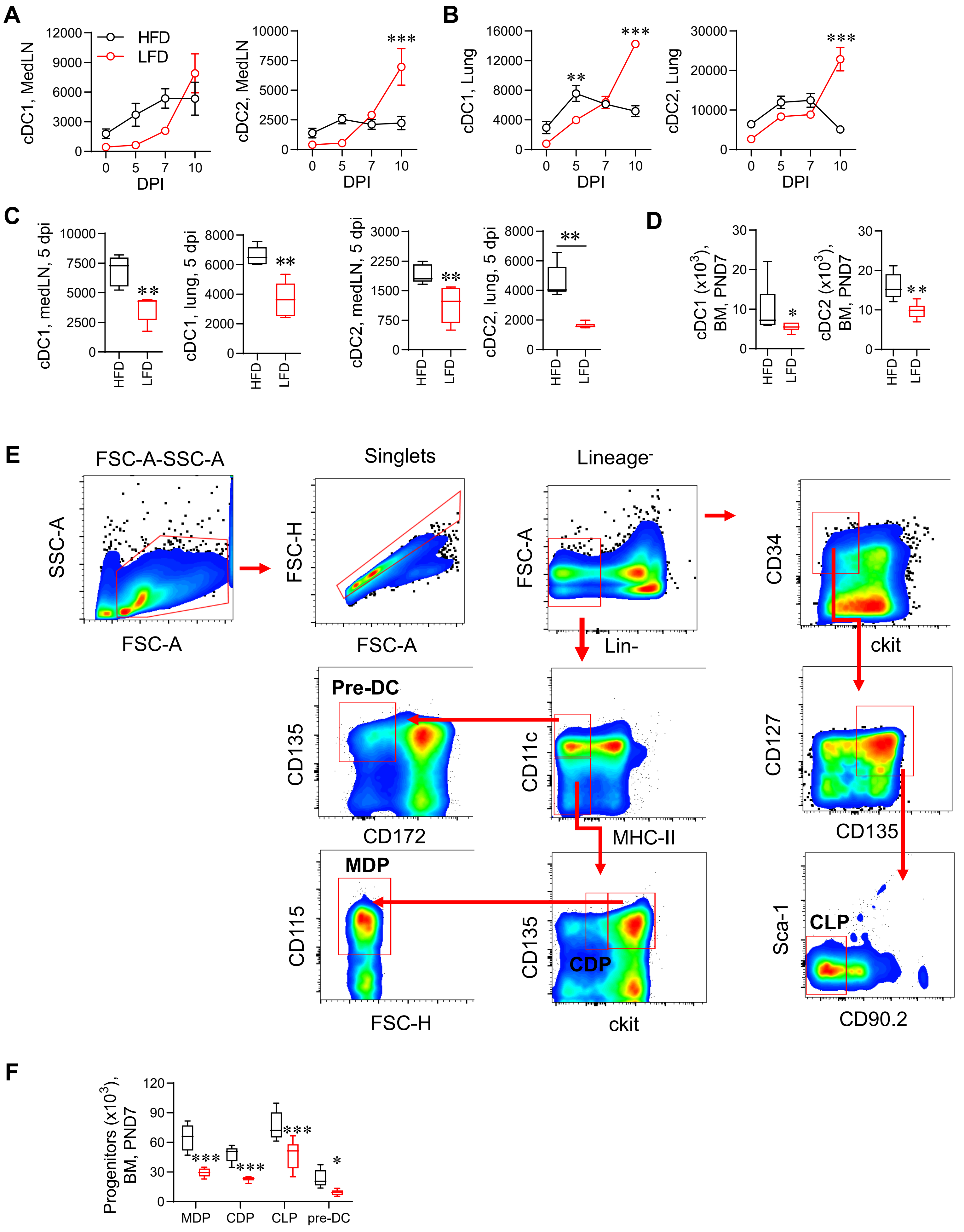

A
cDC1, MedLN
cDC2, MedLN
HFD
LFD
DPI
B
cDC1, Lung
cDC2, Lung
DPI
C
cDC1, medLN, 5 dpi
cDC1, lung, 5 dpi
cDC2, medLN, 5 dpi
cDC2, lung, 5 dpi
HFD
LFD
D
cDC1 (x10^3), BM, PND7
cDC2 (x10^3), BM, PND7
E
FSC-A-SSC-A
Singlets
Lineage-
SSC-A
FSC-A
FSC-H
Lin-
CD34
ckit
Pre-DC
CD135
CD172
CD11c
MHC-II
CD127
MDP
CD115
CDP
Sca-1
CLP
CD90.2
F
Progenitors (x10^3), BM, PND7
MDP
CDP
CLP
pre-DC

### Figure S6

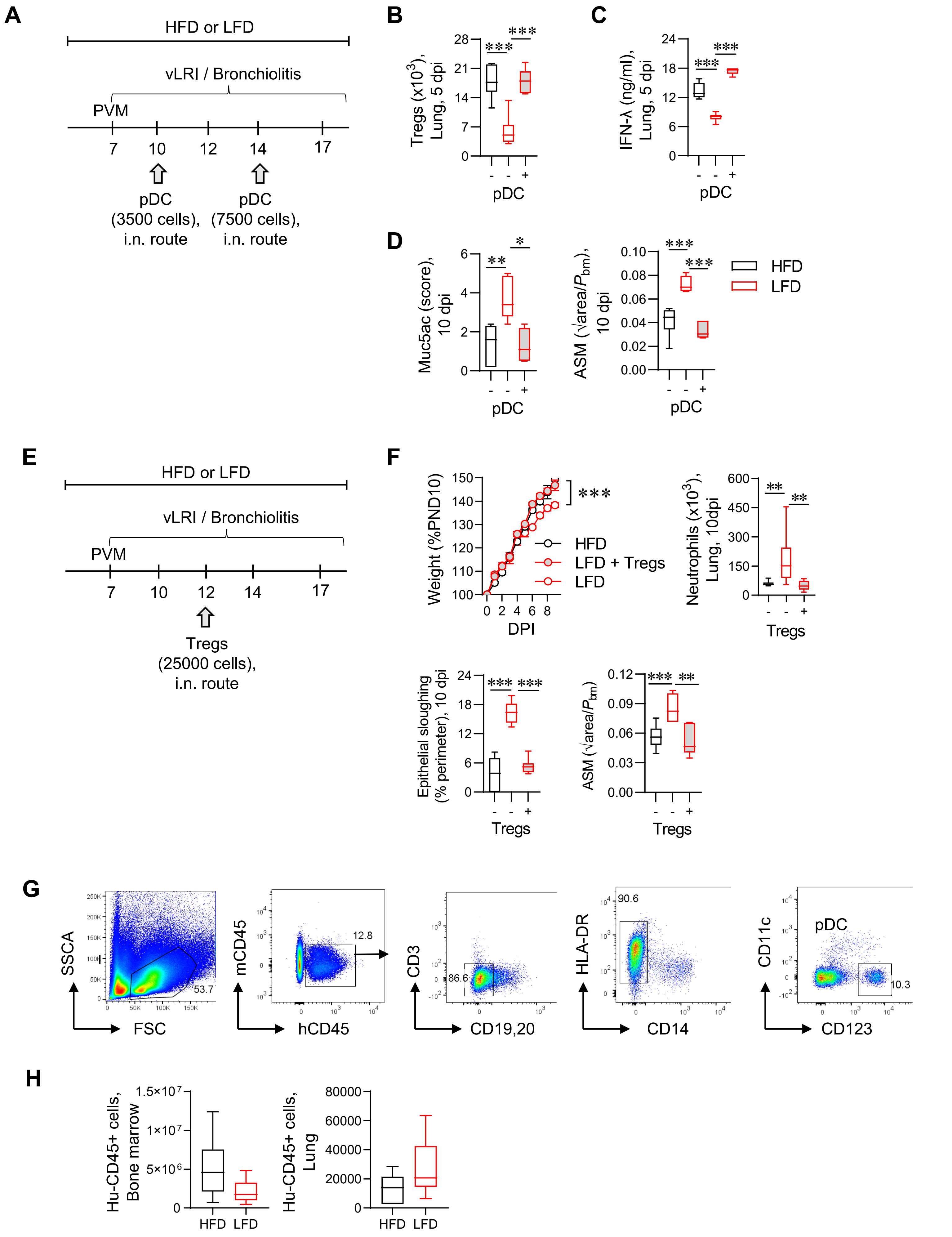
A
HFD or LFD
vLRI / Bronchiolitis
PVM
7 10 12 14 17
pDC (3500 cells), i.n. route
pDC (7500 cells), i.n. route
B
Tregs (x10^3), Lung, 5 dpi
pDC
C
IFN-λ (ng/ml), Lung, 5 dpi
pDC
D
Muc5ac (score), 10 dpi
ASM (√area/Pbm), 10 dpi
pDC
HFD
LFD
E
HFD or LFD
vLRI / Bronchiolitis
PVM
7 10 12 14 17
Tregs (25000 cells), i.n. route
F
Weight (%PND10)
DPI
HFD
LFD + Tregs
LFD
Neutrophils (x10^3), Lung, 10dpi
Tregs
Epithelial sloughing (% perimeter), 10 dpi
ASM (√area/Pbm)
Tregs
G
SSCA
FSC
53.7
mCD45
hCD45
12.8
CD3
CD19,20
86.6
HLA-DR
CD14
90.6
CD11c
CD123
pDC
10.3
H
Hu-CD45+ cells, Bone marrow
Hu-CD45+ cells, Lung
HFD
LFD

**Figure S7**

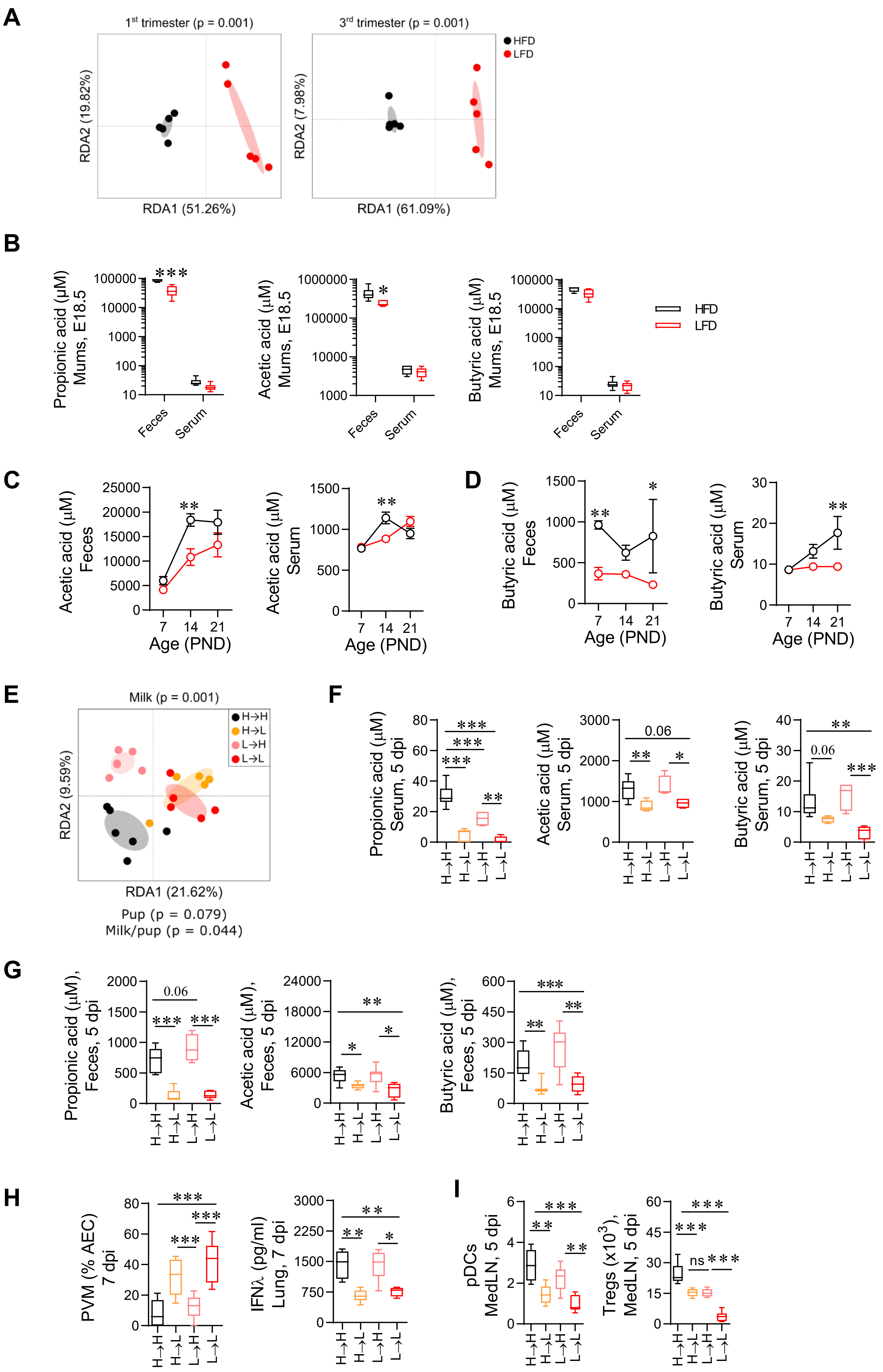
A
1st trimester (p = 0.001)
RDA2 (19.82%)
RDA1 (51.26%)
3rd trimester (p = 0.001)
RDA2 (7.98%)
RDA1 (61.09%)
HFD
LFD
B
Propionic acid (μM) Mums, E18.5
Acetic acid (μM) Mums, E18.5
Butyric acid (μM) Mums, E18.5
Feces
Serum
HFD
LFD
C
Acetic acid (μM) Feces
Acetic acid (μM) Serum
Age (PND)
D
Butyric acid (μM) Feces
Butyric acid (μM) Serum
Age (PND)
E
Milk (p = 0.001)
H→H
H→L
L→H
L→L
RDA2 (9.59%)
RDA1 (21.62%)
Pup (p = 0.079)
Milk/pup (p = 0.044)
F
Propionic acid (μM) Serum, 5 dpi
Acetic acid (μM) Serum, 5 dpi
Butyric acid (μM) Serum, 5 dpi
0.06
G
Propionic acid (μM), Feces, 5 dpi
Acetic acid (μM), Feces, 5 dpi
Butyric acid (μM), Feces, 5 dpi
0.06
H
PVM (% AEC) 7 dpi
IFNλ (pg/ml) Lung, 7 dpi
I
pDCs MedLN, 5 dpi
Tregs (x10³), MedLN, 5 dpi
ns

**Figure S8**

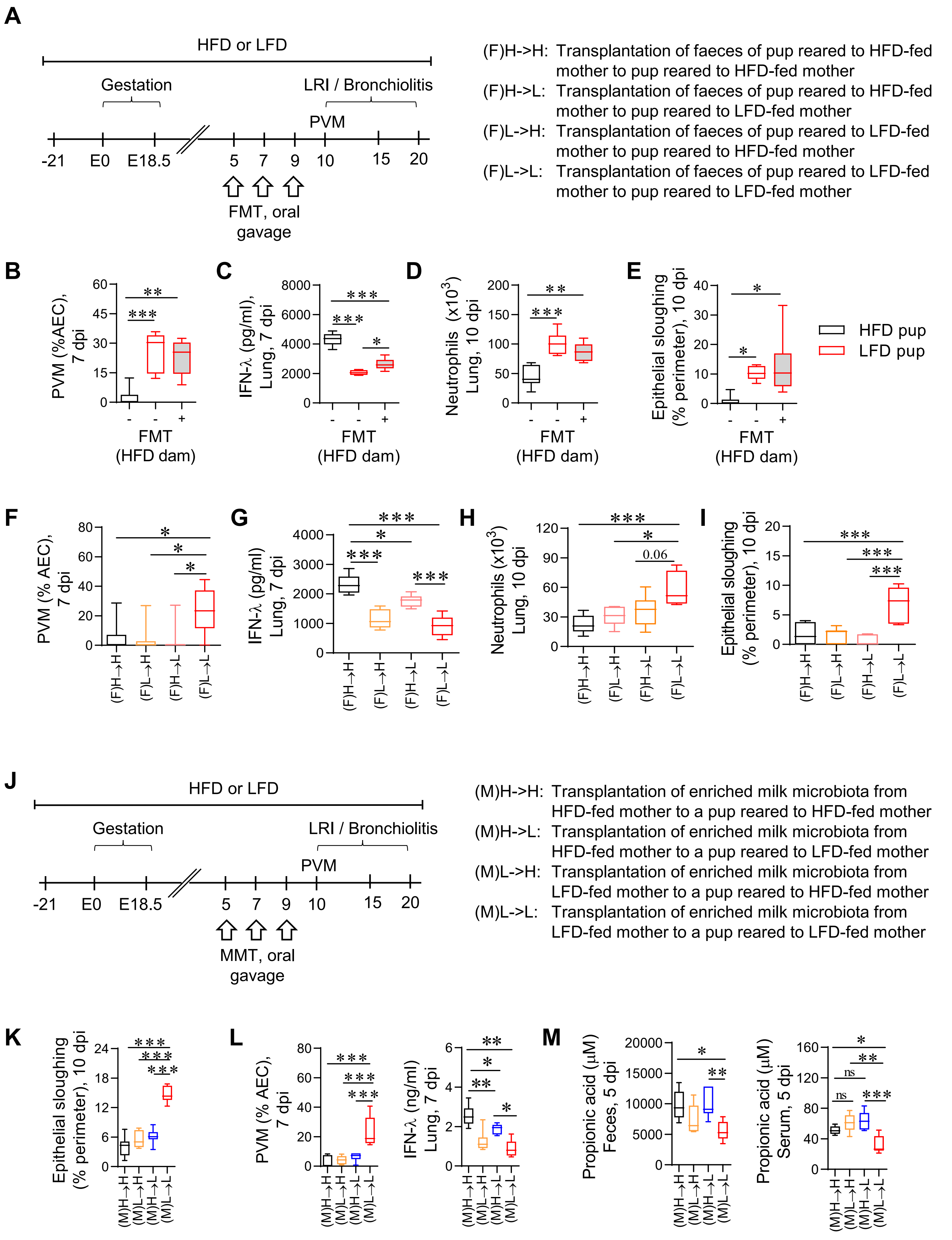
A
HFD or LFD
Gestation
LRI / Bronchiolitis
PVM
-21
E0
E18.5
5
7
9
10
15
20
FMT, oral gavage
(F)H->H: Transplantation of faeces of pup reared to HFD-fed mother to pup reared to HFD-fed mother
(F)H->L: Transplantation of faeces of pup reared to HFD-fed mother to pup reared to LFD-fed mother
(F)L->H: Transplantation of faeces of pup reared to LFD-fed mother to pup reared to HFD-fed mother
(F)L->L: Transplantation of faeces of pup reared to LFD-fed mother to pup reared to LFD-fed mother
B
PVM (%AEC), 7 dpi
FMT (HFD dam)
C
IFN-λ (pg/ml), Lung, 7 dpi
D
Neutrophils (x10³) Lung, 10 dpi
E
Epithelial sloughing (% perimeter), 10 dpi
HFD pup
LFD pup
F
PVM (% AEC), 7 dpi
G
IFN-λ (pg/ml) Lung, 7 dpi
H
Neutrophils (x10³) Lung, 10 dpi
0.06
I
Epithelial sloughing (% perimeter), 10 dpi
(F)H→H
(F)L→H
(F)H→L
(F)L→L
J
HFD or LFD
Gestation
LRI / Bronchiolitis
PVM
MMT, oral gavage
(M)H->H: Transplantation of enriched milk microbiota from HFD-fed mother to a pup reared to HFD-fed mother
(M)H->L: Transplantation of enriched milk microbiota from HFD-fed mother to a pup reared to LFD-fed mother
(M)L->H: Transplantation of enriched milk microbiota from LFD-fed mother to a pup reared to HFD-fed mother
(M)L->L: Transplantation of enriched milk microbiota from LFD-fed mother to a pup reared to LFD-fed mother
K
Epithelial sloughing (% perimeter), 10 dpi
L
PVM (% AEC), 7 dpi
IFN-λ (ng/ml) Lung, 7 dpi
M
Propionic acid (μM) Feces, 5 dpi
Propionic acid (μM) Serum, 5 dpi
ns
(M)H→H
(M)L→H
(M)H→L
(M)L→L

**Figure S9**

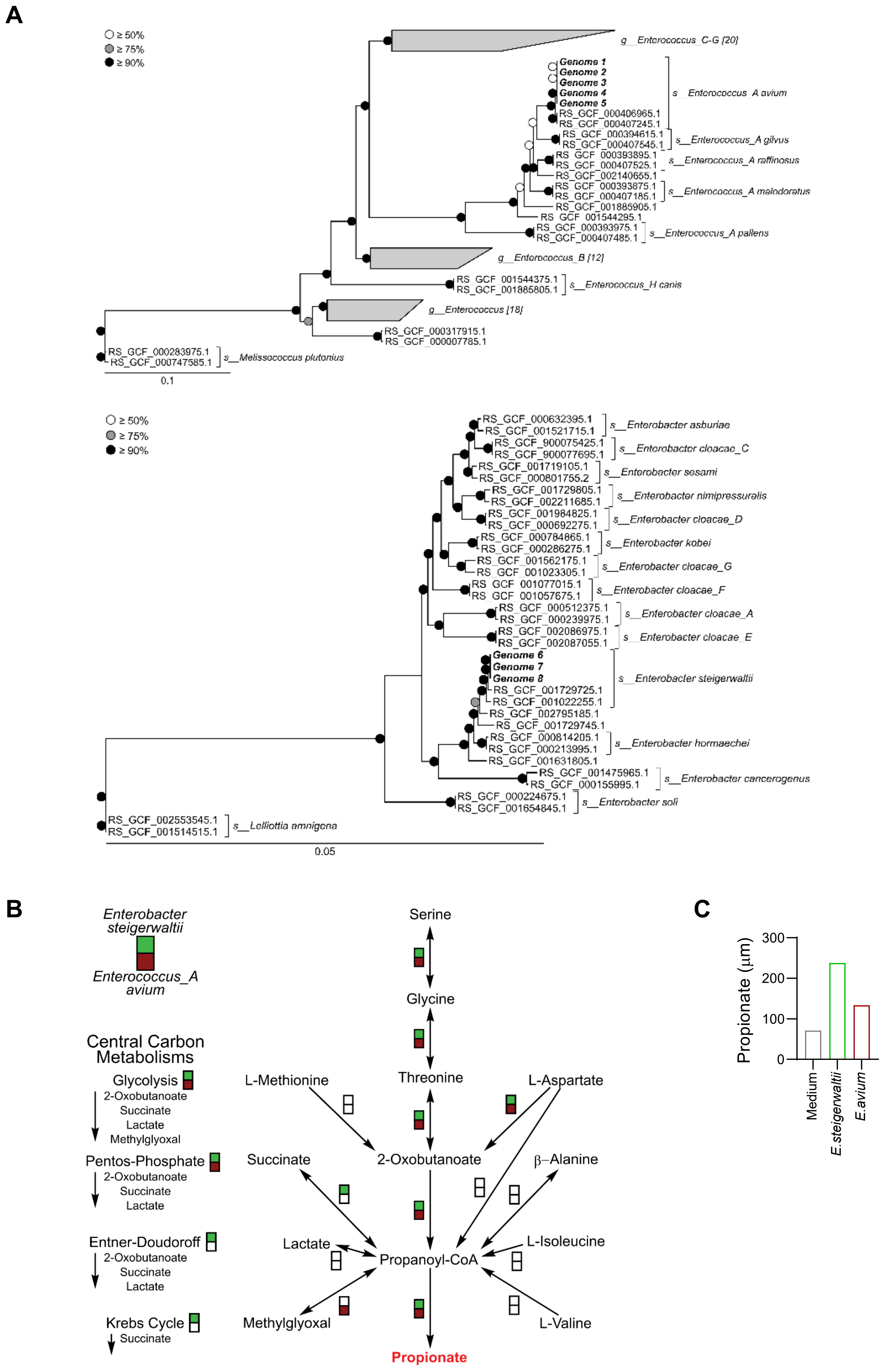
A
≥ 50%
≥ 75%
≥ 90%
g__Enterococcus_C-G [20]
Genome 1
Genome 2
Genome 3
Genome 4
Genome 5
RS_GCF_000406965.1
RS_GCF_000407245.1
s__Enterococcus_A avium
RS_GCF_000394615.1
RS_GCF_000407545.1
s__Enterococcus_A gilvus
RS_GCF_000393895.1
RS_GCF_000407525.1
s__Enterococcus_A raffinosus
RS_GCF_002140655.1
RS_GCF_000393875.1
RS_GCF_000407185.1
s__Enterococcus_A malodoratus
RS_GCF_001885905.1
RS_GCF_001544295.1
RS_GCF_000393975.1
RS_GCF_000407485.1
s__Enterococcus_A pallens
g__Enterococcus_B [12]
RS_GCF_001544375.1
RS_GCF_001885805.1
s__Enterococcus_H canis
g__Enterococcus [18]
RS_GCF_000317915.1
RS_GCF_000007785.1
RS_GCF_000283975.1
RS_GCF_007747585.1
s__Melissococcus plutonius
0.1
≥ 50%
≥ 75%
≥ 90%
RS_GCF_000632395.1
RS_GCF_001521715.1
s__Enterobacter asburiae
RS_GCF_900075425.1
RS_GCF_900077695.1
s__Enterobacter cloacae_C
RS_GCF_001719105.1
RS_GCF_000801755.2
s__Enterobacter sesami
RS_GCF_001729805.1
RS_GCF_002211685.1
s__Enterobacter nimipressuralis
RS_GCF_001984825.1
RS_GCF_000692275.1
s__Enterobacter cloacae_D
RS_GCF_000784865.1
RS_GCF_000286275.1
s__Enterobacter kobei
RS_GCF_001562175.1
RS_GCF_001023305.1
s__Enterobacter cloacae_G
RS_GCF_001077015.1
RS_GCF_001057675.1
s__Enterobacter cloacae_F
RS_GCF_000512375.1
RS_GCF_000239975.1
s__Enterobacter cloacae_A
RS_GCF_002086975.1
RS_GCF_002087055.1
s__Enterobacter cloacae_E
Genome 6
Genome 7
Genome 8
RS_GCF_001729725.1
RS_GCF_001022255.1
s__Enterobacter steigerwaltii
RS_GCF_002795185.1
RS_GCF_001729745.1
RS_GCF_000814205.1
RS_GCF_000213995.1
s__Enterobacter hormaechei
RS_GCF_001631805.1
RS_GCF_001475965.1
RS_GCF_000155995.1
s__Enterobacter cancerogenus
RS_GCF_000224675.1
RS_GCF_001654845.1
s__Enterobacter soli
RS_GCF_002553545.1
RS_GCF_001514515.1
s__Lelliottia amnigena
0.05
B
Enterobacter steigerwaltii
Enterococcus_A avium
Central Carbon Metabolisms
Glycolysis
2-Oxobutanoate
Succinate
Lactate
Methylglyoxal
Pentos-Phosphate
2-Oxobutanoate
Succinate
Lactate
Entner-Doudoroff
2-Oxobutanoate
Succinate
Lactate
Krebs Cycle
Succinate
Serine
Glycine
Threonine
L-Methionine
L-Aspartate
Succinate
2-Oxobutanoate
β–Alanine
Lactate
Propanoyl-CoA
L-Isoleucine
Methylglyoxal
L-Valine
Propionate
C
Propionate (µm)
300
200
100
0
Medium
E.steigerwaltii
E.avium

**Figure S10**

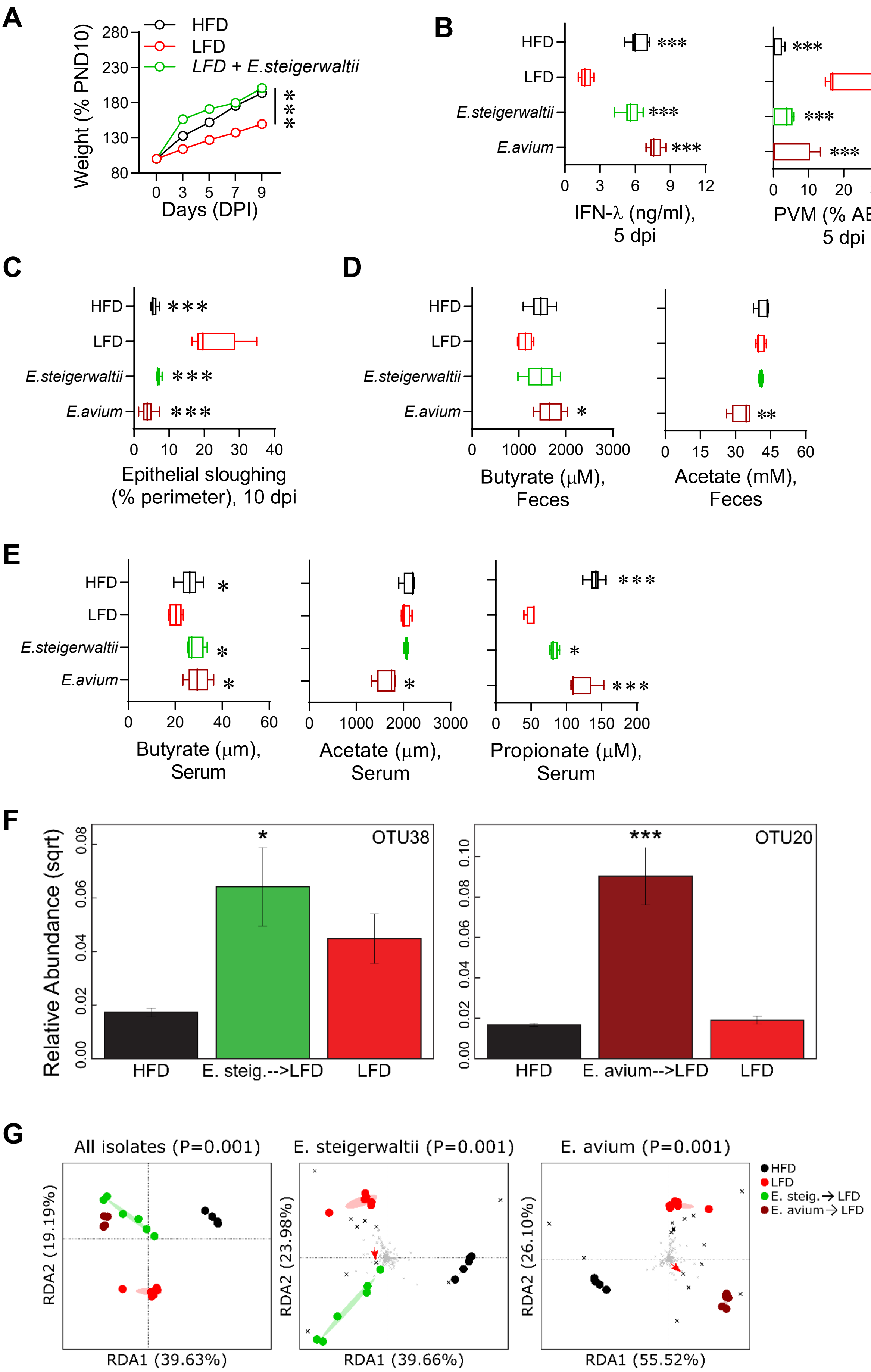
A
Weight (% PND10)
Days (DPI)
HFD
LFD
LFD + E.steigerwaltii
B
IFN-λ (ng/ml), 5 dpi
PVM (% AECs), 5 dpi
HFD
LFD
E.steigerwaltii
E.avium
C
Epithelial sloughing (% perimeter), 10 dpi
D
Butyrate (μM), Feces
Acetate (mM), Feces
E
Butyrate (μm), Serum
Acetate (μm), Serum
Propionate (μM), Serum
F
Relative Abundance (sqrt)
OTU38
OTU20
HFD
E. steig.-->LFD
LFD
E. avium-->LFD
G
All isolates (P=0.001)
E. steigerwaltii (P=0.001)
E. avium (P=0.001)
RDA1 (39.63%)
RDA2 (19.19%)
RDA1 (39.66%)
RDA2 (23.98%)
RDA1 (55.52%)
RDA2 (26.10%)
HFD
LFD
E. steig.→ LFD
E. avium→ LFD

#### Figure S11

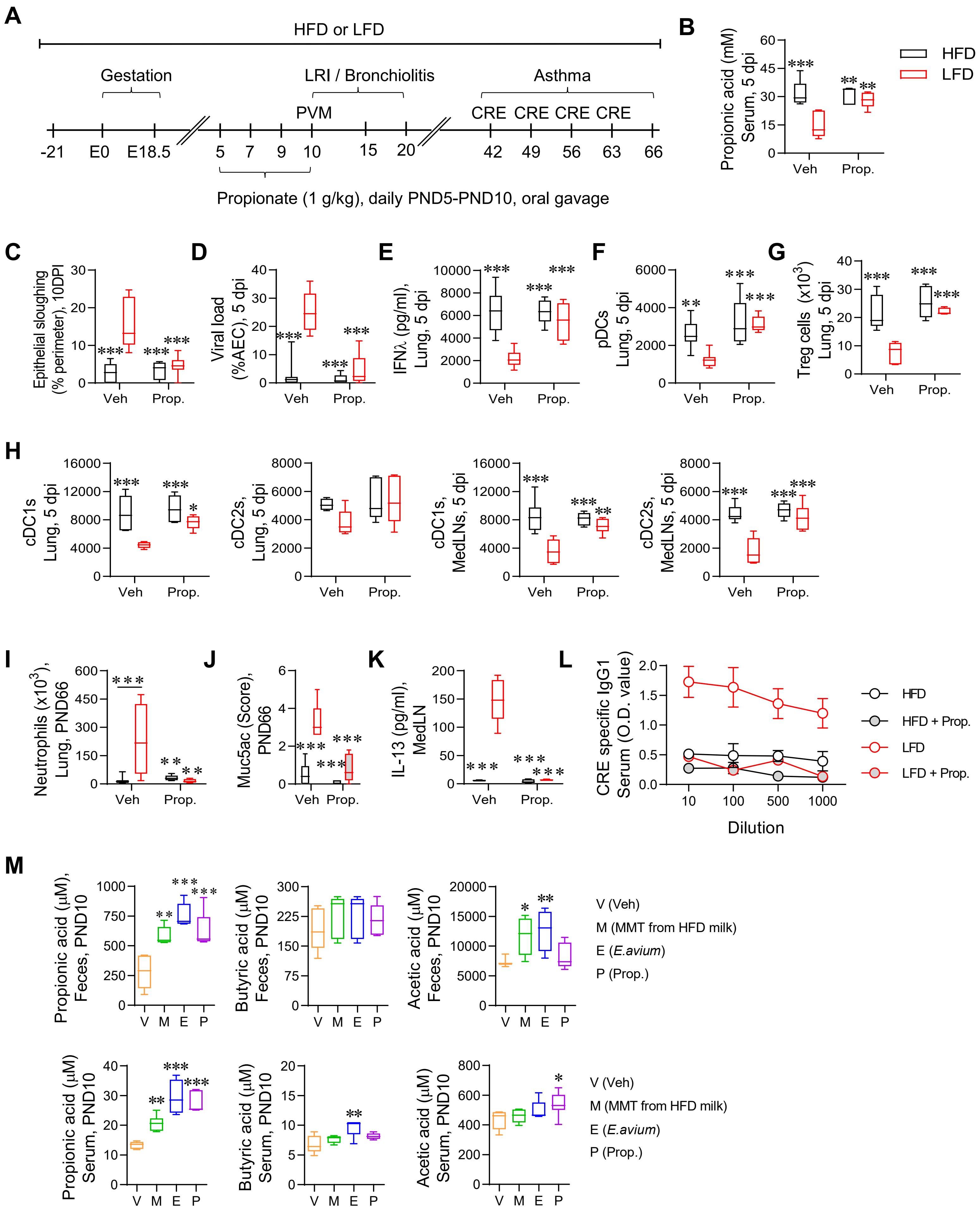
A
HFD or LFD
Gestation
LRI / Bronchiolitis
Asthma
PVM
CRE CRE CRE CRE
-21 E0 E18.5 5 7 9 10 15 20 42 49 56 63 66
Propionate (1 g/kg), daily PND5-PND10, oral gavage
B
Propionic acid (mM) Serum, 5 dpi
HFD
LFD
Veh
Prop.
C
Epithelial sloughing (% perimeter), 10DPI
D
Viral load (%AEC), 5 dpi
E
IFNλ (pg/ml), Lung, 5 dpi
F
pDCs Lung, 5 dpi
G
Treg cells (x10$^3$) Lung, 5 dpi
H
cDC1s Lung, 5 dpi
cDC2s, Lung, 5 dpi
cDC1s, MedLNs, 5 dpi
cDC2s, MedLNs, 5 dpi
I
Neutrophils (x10$^3$), Lung, PND66
J
Muc5ac (Score), PND66
K
IL-13 (pg/ml), MedLN
L
CRE specific IgG1 Serum (O.D. value)
Dilution
HFD
HFD + Prop.
LFD
LFD + Prop.
M
Propionic acid (μM), Feces, PND10
Butyric acid (μM), Feces, PND10
Acetic acid (μM), Feces, PND10
V (Veh)
M (MMT from HFD milk)
E (E.avium)
P (Prop.)
Propionic acid (μM) Serum, PND10
Butyric acid (μM) Serum, PND10
Acetic acid (μM) Serum, PND10
V (Veh)
M (MMT from HFD milk)
E (E.avium)
P (Prop.)

### Figure S12

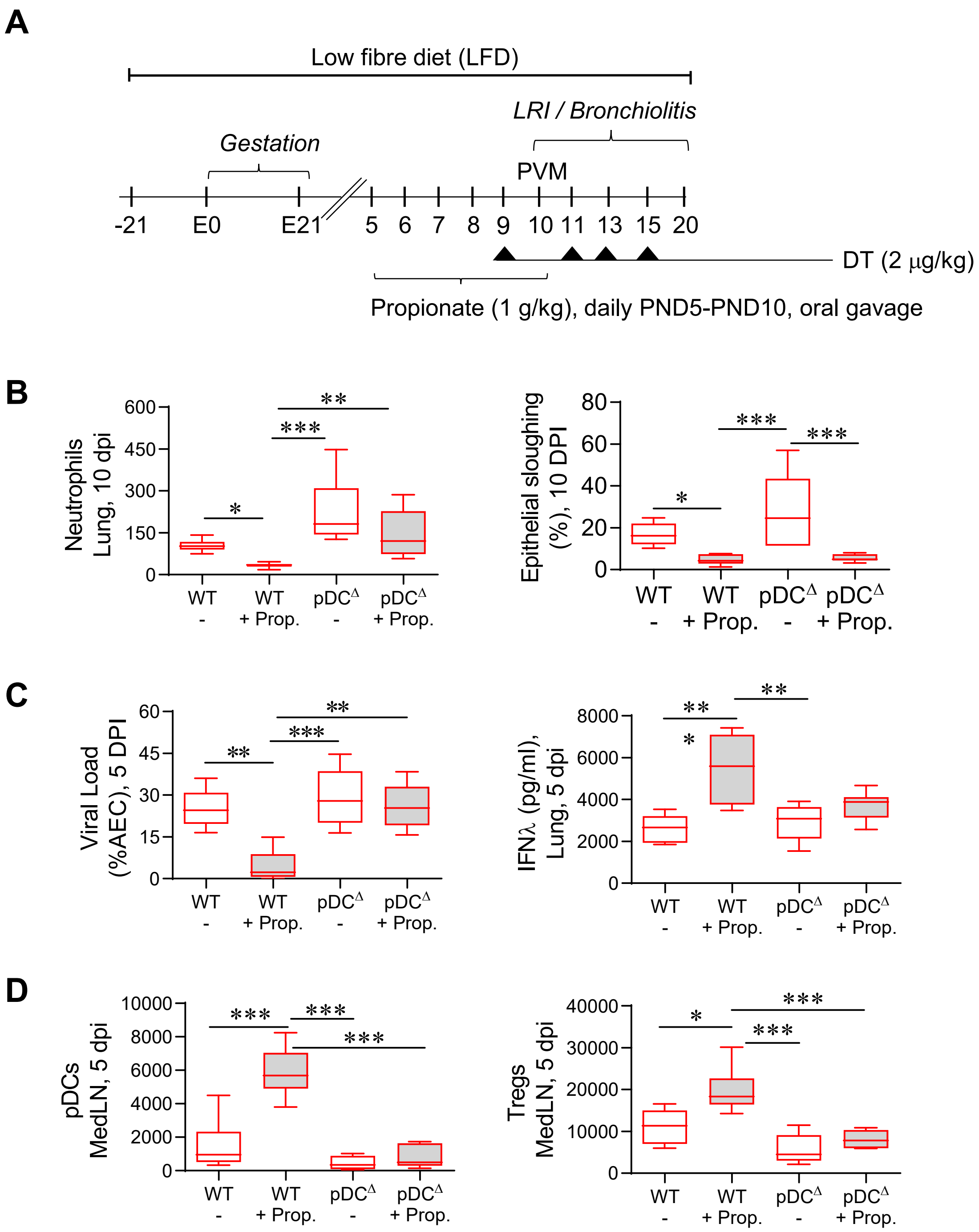
A
Low fibre diet (LFD)
LRI / Bronchiolitis
Gestation
PVM
-21
E0
E21
5
6
7
8
9
10
11
13
15
20
DT (2 μg/kg)
Propionate (1 g/kg), daily PND5-PND10, oral gavage
B
Neutrophils Lung, 10 dpi
Epithelial sloughing (%), 10 DPI
C
Viral Load (%AEC), 5 DPI
IFNλ (pg/ml), Lung, 5 dpi
D
pDCs MedLN, 5 dpi
Tregs MedLN, 5 dpi
WT
-
WT
+ Prop.
pDC^Δ
-
pDC^Δ
+ Prop.

### Figure S13

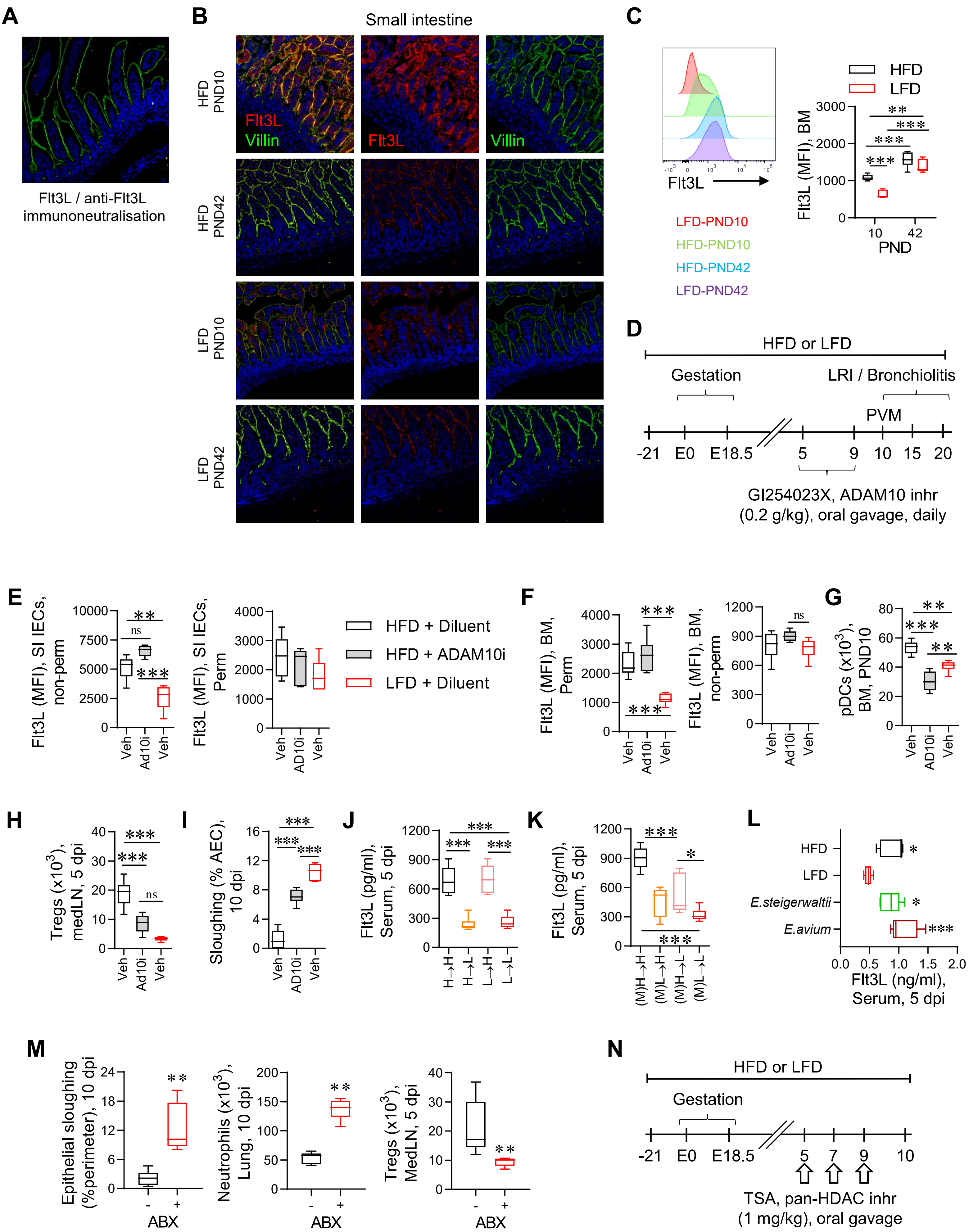

A
Flt3L / anti-Flt3L immunoneutralisation
B
Small intestine
HFD PND10
HFD PND42
LFD PND10
LFD PND42
Flt3L
Villin
C
Flt3L
LFD-PND10
HFD-PND10
HFD-PND42
LFD-PND42
HFD
LFD
Flt3L (MFI), BM
PND
D
HFD or LFD
Gestation
LRI / Bronchiolitis
PVM
-21 E0 E18.5 5 9 10 15 20
GI254023X, ADAM10 inhr (0.2 g/kg), oral gavage, daily
E
Flt3L (MFI), SI IECs, non-perm
Flt3L (MFI), SI IECs, Perm
HFD + Diluent
HFD + ADAM10i
LFD + Diluent
F
Flt3L (MFI), BM, Perm
Flt3L (MFI), BM, non-perm
G
pDCs (x10^3), BM, PND10
H
Tregs (x10^3), medLN, 5 dpi
I
Sloughing (% AEC), 10 dpi
J
Flt3L (pg/ml), Serum, 5 dpi
K
Flt3L (pg/ml), Serum, 5 dpi
L
HFD
LFD
E.steigerwaltii
E.avium
Flt3L (ng/ml), Serum, 5 dpi
M
Epithelial sloughing (%perimeter), 10 dpi
Neutrophils (x10^3), Lung, 10 dpi
Tregs (x10^3), MedLN, 5 dpi
ABX
N
HFD or LFD
Gestation
-21 E0 E18.5 5 7 9 10
TSA, pan-HDAC inhr (1 mg/kg), oral gavage

**Figure S14**

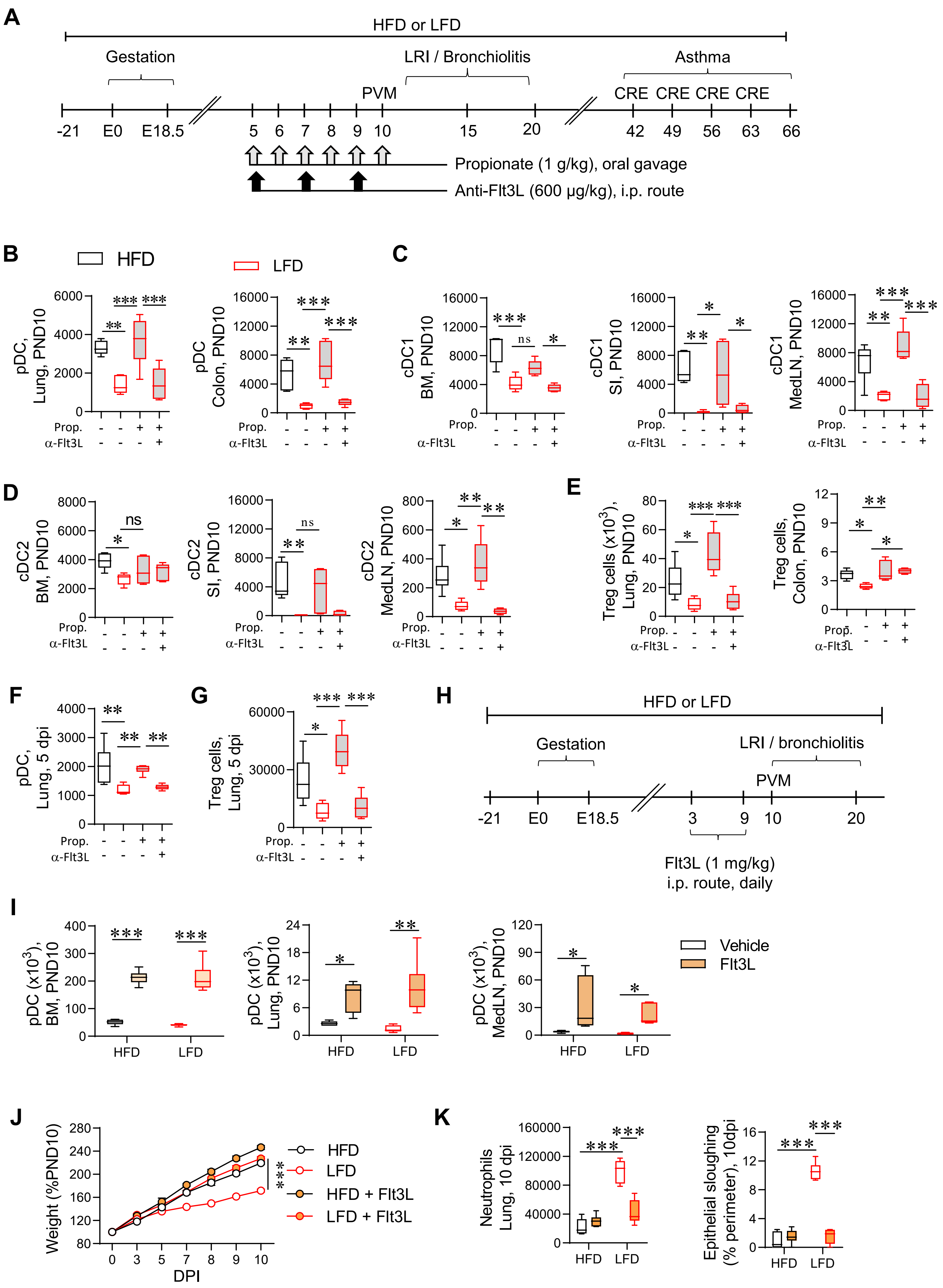
A
HFD or LFD
Gestation
LRI / Bronchiolitis
Asthma
PVM
CRE CRE CRE CRE
-21 E0 E18.5 5 6 7 8 9 10 15 20 42 49 56 63 66
Propionate (1 g/kg), oral gavage
Anti-Flt3L (600 µg/kg), i.p. route
B
HFD
LFD
pDC, Lung, PND10
pDC Colon, PND10
C
cDC1 BM, PND10
cDC1 SI, PND10
cDC1 MedLN, PND10
D
cDC2 BM, PND10
cDC2 SI, PND10
cDC2 MedLN, PND10
E
Treg cells (x10^3), Lung, PND10
Treg cells, Colon, PND10
Prop.
α-Flt3L
F
pDC, Lung, 5 dpi
G
Treg cells, Lung, 5 dpi
H
HFD or LFD
Gestation
LRI / bronchiolitis
PVM
-21 E0 E18.5 3 9 10 20
Flt3L (1 mg/kg)
i.p. route, daily
I
pDC (x10^3), BM, PND10
pDC (x10^3), Lung, PND10
pDC (x10^3), MedLN, PND10
Vehicle
Flt3L
HFD
LFD
J
Weight (%PND10)
DPI
HFD
LFD
HFD + Flt3L
LFD + Flt3L
K
Neutrophils Lung, 10 dpi
Epithelial sloughing (% perimeter), 10dpi
